# Genetic elucidation of complex biochemical traits mediating maize innate immunity

**DOI:** 10.1101/2020.03.04.977355

**Authors:** Yezhang Ding, Philipp R. Weckwerth, Elly Poretsky, Katherine M. Murphy, James Sims, Evan Saldivar, Shawn A. Christensen, Si Nian Char, Bing Yang, Anh-dao Tong, Zhouxin Shen, Karl A. Kremling, Edward S. Buckler, Tom Kono, David R. Nelson, Jörg Bohlmann, Matthew G. Bakker, Martha M. Vaughan, Ahmed S. Khalil, Mariam Betsiashvili, Steven P. Briggs, Philipp Zerbe, Eric A. Schmelz, Alisa Huffaker

**Affiliations:** Section of Cell and Developmental Biology, University of California at San Diego, La Jolla, CA USA; Department of Plant Biology, University of California Davis, One Shields Avenue, Davis, CA, USA; ETH Zurich, Institute of Agricultural Sciences, Zurich, Switzerland; Chemistry Research Unit, Center for Medical, Agricultural, and Veterinary Entomology, Department of Agriculture–Agricultural Research Service, Gainesville, FL, USA; Division of Plant Sciences, Bond Life Sciences Center, University of Missouri, Columbia, MO, USA; Donald Danforth Plant Science Center, St. Louis, MO, USA; Department of Plant Breeding and Genetics, Cornell University, Ithaca, NY, USA; United States Department of Agriculture-Agricultural Research Service, Robert W. Holley Center for Agriculture and Health, Ithaca, New York, USA; Minnesota Supercomputing Institute, University of Minnesota, Minneapolis, MN USA; University of Tennessee Health Science Center, Memphis, TN, USA; Michael Smith Laboratories, University of British Columbia, Vancouver, British Columbia, Canada; National Center for Agricultural Utilization Research, United States Department of Agriculture-Agricultural Research Service, Peoria, IL, USA; Department of Microbiology, University of Manitoba, Winnipeg, Manitoba, Canada.

## Abstract

Specialized metabolites constitute key layers of immunity underlying crop resistance; however, challenges in resolving complex pathways limit our understanding of their functions and applications. In maize (*Zea mays*) the inducible accumulation of acidic terpenoids is increasingly considered as a defense regulating disease resistance. To understand maize antibiotic biosynthesis, we integrated association mapping, pan-genome multi-omic correlations, enzyme structure-function studies, and targeted mutagenesis. We now define ten genes in three zealexin (Zx) gene clusters comprised of four sesquiterpene synthases and six cytochrome P450s that collectively drive the production of diverse antibiotic cocktails. Quadruple mutants blocked in the production of β-macrocarpene exhibit a broad-spectrum loss of disease resistance. Genetic redundancies ensuring pathway resiliency to single null mutations are combined with enzyme substrate-promiscuity creating a biosynthetic hourglass pathway utilizing diverse substrates and *in vivo* combinatorial chemistry to yield complex antibiotic blends. The elucidated genetic basis of biochemical phenotypes underlying disease resistance demonstrates a predominant maize defense pathway and informs innovative strategies for transferring chemical immunity between crops.

## Introduction

Currently 50% of global arable land is allocated to agriculture. As the world’s largest annually harvested crop, maize (Zea mays) contributes to the significant footprint of poaceous cereals. In the absence of yearly improvements in germplasm and cereal productivity, comparable land consumption today would be greater than 60%^1, 2^. Given human reliance on a few related grasses, genetically encoded mechanisms providing crop stress protection have long been sought^3–5^. In particular, fungal diseases, such as those caused by *Fusarium* species including *F. graminearum,* are widely devastating to poaceous crops, resulting in both significant yield losses and grain contamination with harmful mycotoxins^6, 7^. The understanding of innate immune responses, crop genetic variation and endogenous pathway interactions underlying broad-spectrum disease resistance^8^ represents foundational knowledge necessary for sustained improvement and crop trait optimization.

Plants are protected from pest and pathogen attack by interconnected layers of physical barriers, pattern-recognition receptors, defense proteins and bioactive specialized metabolites^9–11^. Specialized metabolic pathways are often unique to individual species, display specificity in regulated production and mediate cryptic yet impactful phenotypes^9, 12^. Benzoxazinoids are the most broadly shared and widely studied poaceous chemical defenses. Constitutively produced in seedlings, benzoxazinoids contribute to resistance against insects and fungi such as northern corn leaf blight (*Setosphaeria turcica*)^5, 13–15^. In contrast to benzoxazinoids and other largely constitutive defenses present prior to attack, many specialized metabolites are produced exclusively on demand, display extreme localization and often evade analytical detection^16^. Maize relies on a combination of dynamically-regulated benzoxazinoids, phenylpropanoids and terpenoids for biotic and abiotic stress protection^5, 17–19^. While the biosynthesis and roles of benzoxazinoids and terpene volatiles in anti-herbivore defenses are increasingly understood^13, 20^, the genetic and biochemical complexities underlying maize protection against fungal pathogens have remained a challenge to resolve^18, 21–23^.

Terpenoids are the most structurally diverse class of plant specialized metabolites and are typically produced from the combined activities of terpene synthase (TPS) and cytochrome P450 monooxygenase (P450) enzymes^24, 25^. Known maize terpenoid antibiotics include α/β-costic acids, dolabralexins and kauralexins^18, 26, 27^. Despite advances in diterpenoid pathway elucidation^18, 28^, acidic sesquiterpenoid derivatives of β-macrocarpene, termed zealexins, represent the single largest class of defensive terpenoids known in the genus *Zea*^29^, and yet remain the least understood. The endogenous accumulation of zealexins correlates with the expression of genes encoding β-macrocarpene synthases, namely ZmTPS6 and ZmTPS11, which likewise display dramatic transcriptional increases following challenge with diverse fungal pathogens^29–32^. Consistent with crop protection roles, viral silencing of *ZmTPS6* ⁄*11* revealed the first identified maize genes required to restrict smut fungus (*Ustilago maydis*) infection and tumor formation^33^. While correlations between *ZmTPS6* ⁄*11* transcripts, zealexin production and fungal resistance exist, all known maize lines produce zealexins and no single biosynthetic pathway node has been proven *in planta*. Many catalytic activities and biological roles have been assigned to the 43 TPS encoded in the maize B73 genome^34, 35^; however, the structural diversity of zealexins, combinations of underlying TPS/P450 terpenoid-diversifying genes, and the endogenous protective function of the zealexin pathway remains unresolved^27, 36^. Recent advances in omic tools, co-regulation analyses, genetic resources, *in vivo* protein biochemistry and gene editing approaches now enable the critical examination and engineering of complex protective pathways underlying crop resistance.

To define the genetic basis, pan-genome complexity and biochemical layers of immunity mediating maize disease resistance, this study identifies 17 metabolites as products of the core zealexin (Zx) pathway (*Zx1* to *Zx10*) that consists of three functionally distinct gene clusters encoding ZmTPS responsible for hydrocarbon olefin production and P450s in the ZmCYP71Z and ZmCYP81A families facilitating oxygenation and desaturation. Enzyme promiscuity within the zealexin pathway enables formation of an expansive cocktail of terpenoid antibiotics through conversion of multiple endogenous precursors via a biosynthetic hourglass pathway. While fungal challenge is complex and results in the large-scale alteration of greater than half the measurable proteome, *zx1 zx2 zx3 zx4* quadruple mutants demonstrate that zealexins are important biochemical defenses significantly contributing to protection against *F. graminearum* stalk rot. Given that all plants produce terpenoid precursors, the promiscuous enzyme activities described are amenable to genetic transfer. A foundational understanding of genetic and biochemical mechanisms in maize lays the groundwork for diversifying chemical defenses underlying disease resistance phenotypes in phylogenetically distant grain crops^37^.

## RESULTS

### Maize harbors a functionally variable gene cluster of four β-macrocarpene synthases

Two pathogen regulated β-macrocarpene synthases, termed Terpene Synthases (TPS) 6 and 11, were previously assigned as the B73 (RefGen_V4) genes *Zm00001d024207* and *Zm00001d024210* (www.maizegdb.org), respectively^30, 31^. Current indirect evidence supports ZmTPS6/11 in the production of diverse antibiotics, termed zealexins^29, 32^. As a visual aid, all biochemicals and genes with examined relevance to the zealexin (Zx) pathway present in the genus *Zea* are now summarized (Supplementary Fig. 1 and 2; Supplementary Tables 1 to 4). Analyses of the B73 genome for all TPS reveal that *ZmTPS6/11* are components of a four-gene cluster on chromosome 10 (Fig. 1a), sharing >84% protein identity one to another (Supplementary Figs. 3 and 4). Following benzoxazinoid pathway nomenclature^5^, we adopted unified B73 Zx pathway abbreviations starting with *Zx1* (*Zm00001d024207*), *Zx2* (*Zm00001d024208*), *Zx3* (*Zm00001d024210*) and *Zx4* (*Zm00001d024211*) based on sequential chromosome order (Fig. 1a-b). Unless otherwise noted, gene and protein abbreviations refer to B73 (RefGen_V4) reference sequences. RNA-seq analyses of *Fusarium*-elicited stem tissues in the inbred lines B73, Mo17 and W22 demonstrate that fungal-elicited transcript accumulation occurs for each of the four genes in an inbred specific manner (Fig. 1c-e; Supplementary Table 2). To understand the contribution of *Zx* gene cluster I to the production of β-macrocarpene and the pathway intermediate β-bisabolene, individual genes *Zx1* to *Zx4* from both B73 and W22 (Supplementary Fig. 4-5 and Supplementary Table 4) were functionally analyzed using transient, *Agrobacterium*-mediated expression in *Nicotiana benthamiana*. B73 (Zx1, Zx3, Zx4) and W22 (Zx2, Zx3, Zx4) shared similar yet different combinations of functional β-macrocarpene synthases yielding low levels of the β-bisabolene intermediate (Fig. 1f-g).

**Fig. 1.**
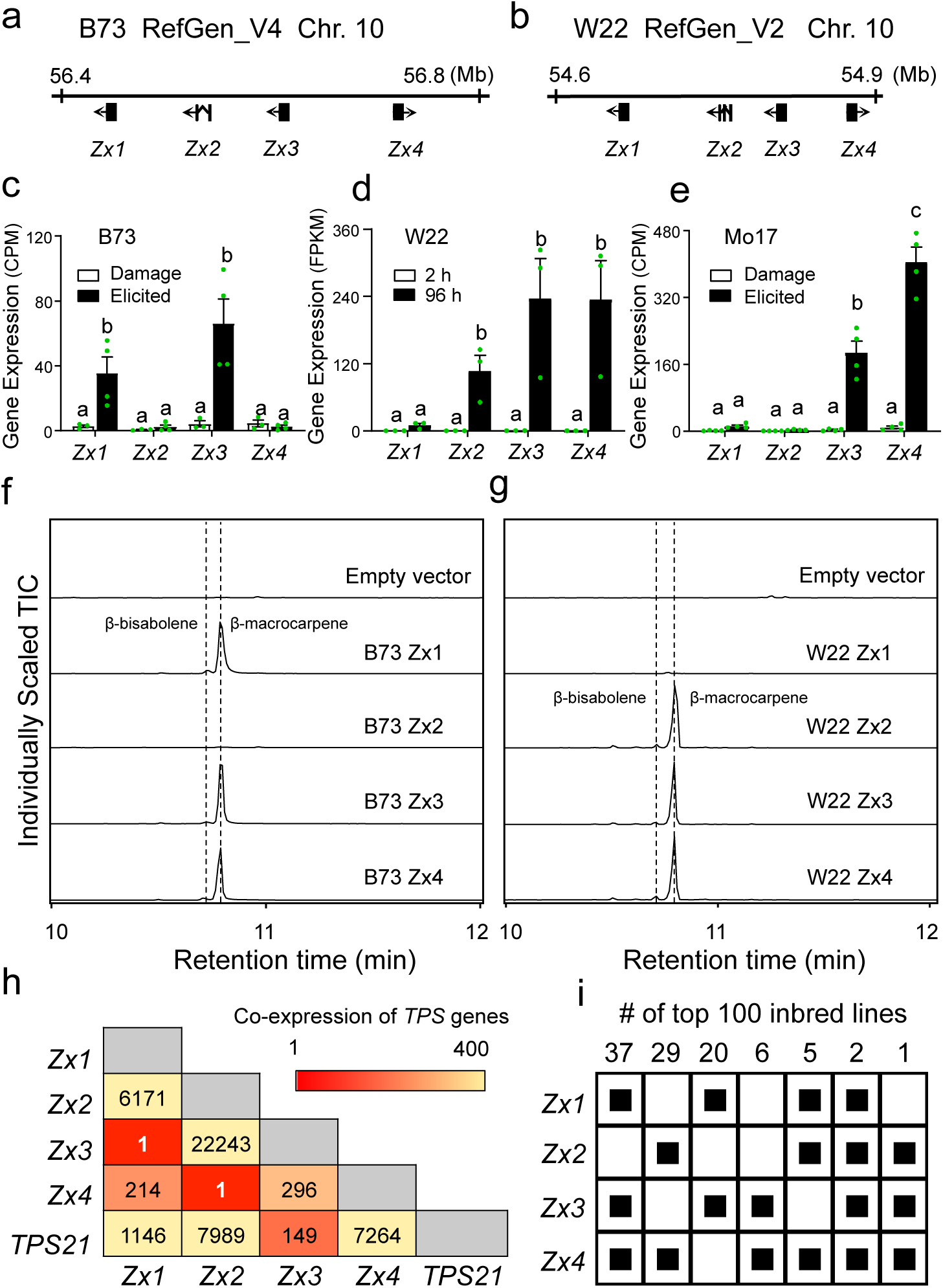
A genetically variable cluster of four terpene synthases ensure the production of zealexin precursors, β-bisabolene and β-macrocarpene. Array of four β-macrocarpene synthase genes, termed *Zx1* to *Zx4*, on chromosome 10 of the (a) B73 genome (RefGen_V4) and the (b) W22 genome (RefGen_V2). *Zx1* to *Zx4* transcript abundance derived from RNA-seq analyses of 5-week old (c) B73, (d) W22 and (e) Mo17 stems damaged (Damage) or additionally treated with heat-killed *F. venenatum* hyphae (Elicited). For Mo17 and B73, harvests occurred at 36 h and 3’-RNA-seq gene expression is given as Counts Per Million mapped reads (CPM). For W22, results average 3 early (0, 2 and 4 h) and late (72, 96, 120 h) elicitation time points with gene expression given as Reads Per Kilobase of transcript per Million mapped reads (RPKM). Error bars in c, d and e indicate mean ± s.e.m. (*n* = 3-4 biologically independent replicates). Within plots, different letters (a–c) represent significant differences (one-way ANOVA followed by Tukey’s test corrections for multiple comparisons, *P* < 0.05). f, g, Total ion chromatograms (TIC) are shown for leaf volatiles emitted following *Agrobacterium*-mediated transient *N. benthamiana* expression assays of B73 and W22 encoded Zx1, Zx2, Zx3 and Zx4. An empty vector was used for the *Agrobacterium*-infiltrated control. h, Heat map depicting the co-expression of genes, *Zx1* to *Zx4* and the β-selinene synthase (*ZmTPS21*), present in 1960 RNA-seq samples. Low numbers indicate supportive mutual rank (MR) scores. i, Expression matrix of *Zx1* to *Zx4* in the top 100 most highly expressing inbred maize lines from 1960 RNA-seq samples. Individual genes with expression levels ≥ 5% of total sum expression of *Zx1* to *Zx4* were counted as expressed and represented by filled black squares (■).

**Fig. 2.**
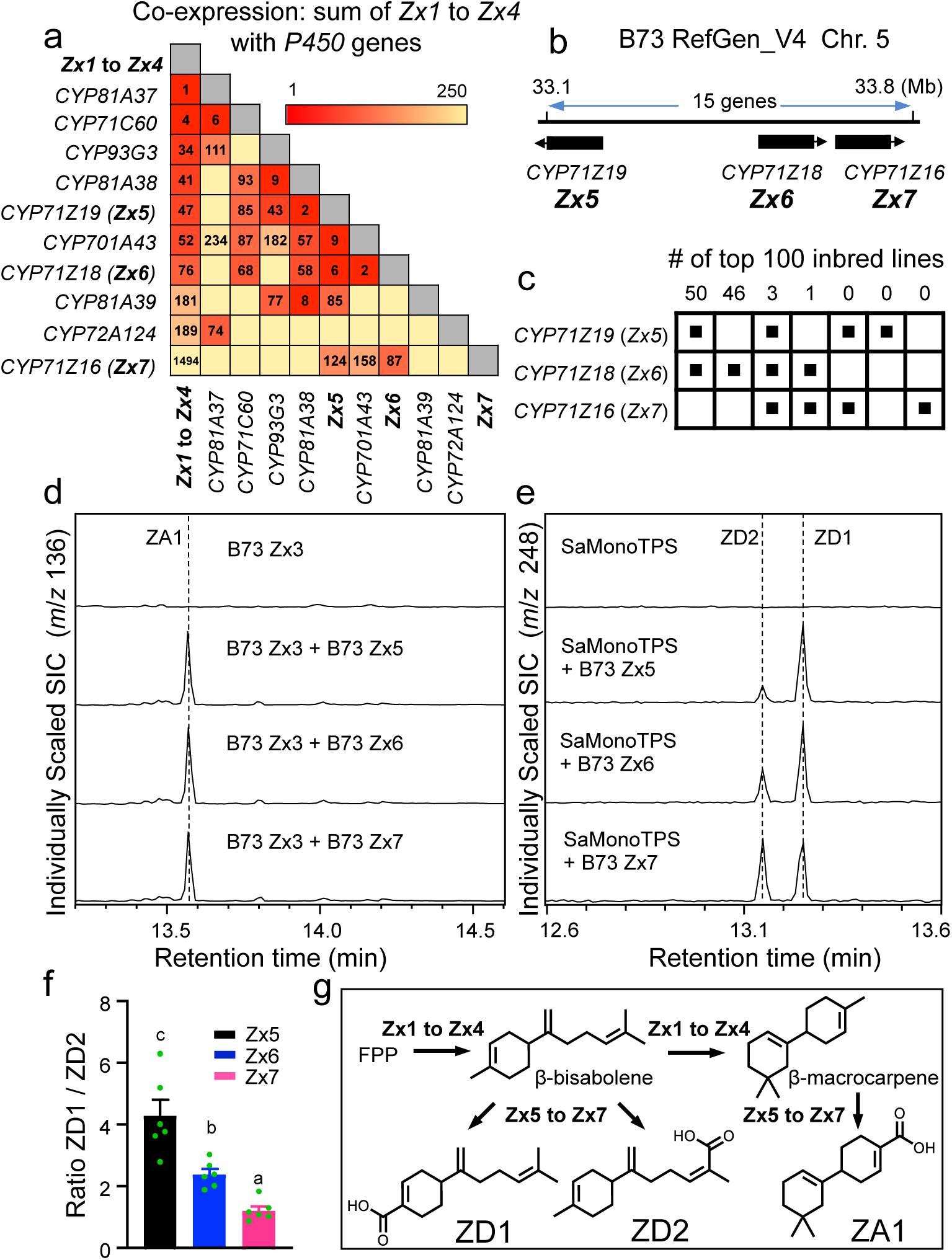
Zealexin gene cluster II contains three 71Z family cytochrome (CYP) P450s that catalyze the production of A and D-series zealexins. **a**, Heatmap depicting the summed expression of *Zx1* to *Zx4* and co-expression with all maize P450s in a dataset of 1960 RNA-seq samples. Low numbers indicate supportive scores while weak MR correlations > 250 were omitted; **b**, Physical position of gene cluster II containing *ZmCYP71Z19* (*Zx5*), *ZmCYP71Z18* (*Zx6*) and *ZmCYP71Z16* (*Zx7*) referenced to the B73 genome (RefGen_V4); **c**, Expression matrix of *Zx5*, *Zx6* and *Zx7* in the top 100 most highly expressing inbred lines present in a dataset of 1960 RNA-seq samples. Genes with expression levels ≥ 5% of *Zx5 to Zx7* summed expression were counted as expressed (filled black squares, ■); **d**, GC-MS selected ion chromatograms (SIC) of extracts derived from *Agrobacterium*-mediated transient *N. benthamiana* co-expression assays of Zx3 individually paired with Zx5, Zx6, or Zx7 all resulted in the production of ZA1; **e**, Parallel co-expression assays of the β-bisabolene synthase from *Santalum album* (SaMonoTPS) with Zx5, Zx6, or Zx7 all resulted in the production of zealexin D1 (ZD1) and zealexin D2 (ZD2) in variable proportions. **f**, Average ratios of ZD1 to ZD2 from co-expression assays of SaMonoTPS with Zx5, Zx6, or Zx7. Error bars indicate mean ± s.e.m. (*n* = 4 biologically independent replicates) and different letters (a–c) represent significant differences (one-way ANOVA followed by Tukey’s test corrections for multiple comparisons, *P* < 0.05). **g**, Schematic representation of farnesyl diphosphate (FPP) cyclization reactions catalyzed by Zx3 and Zx5 to Zx7 yield the acidic zealexins ZD1, ZD2 and ZA1.

**Fig. 3.**
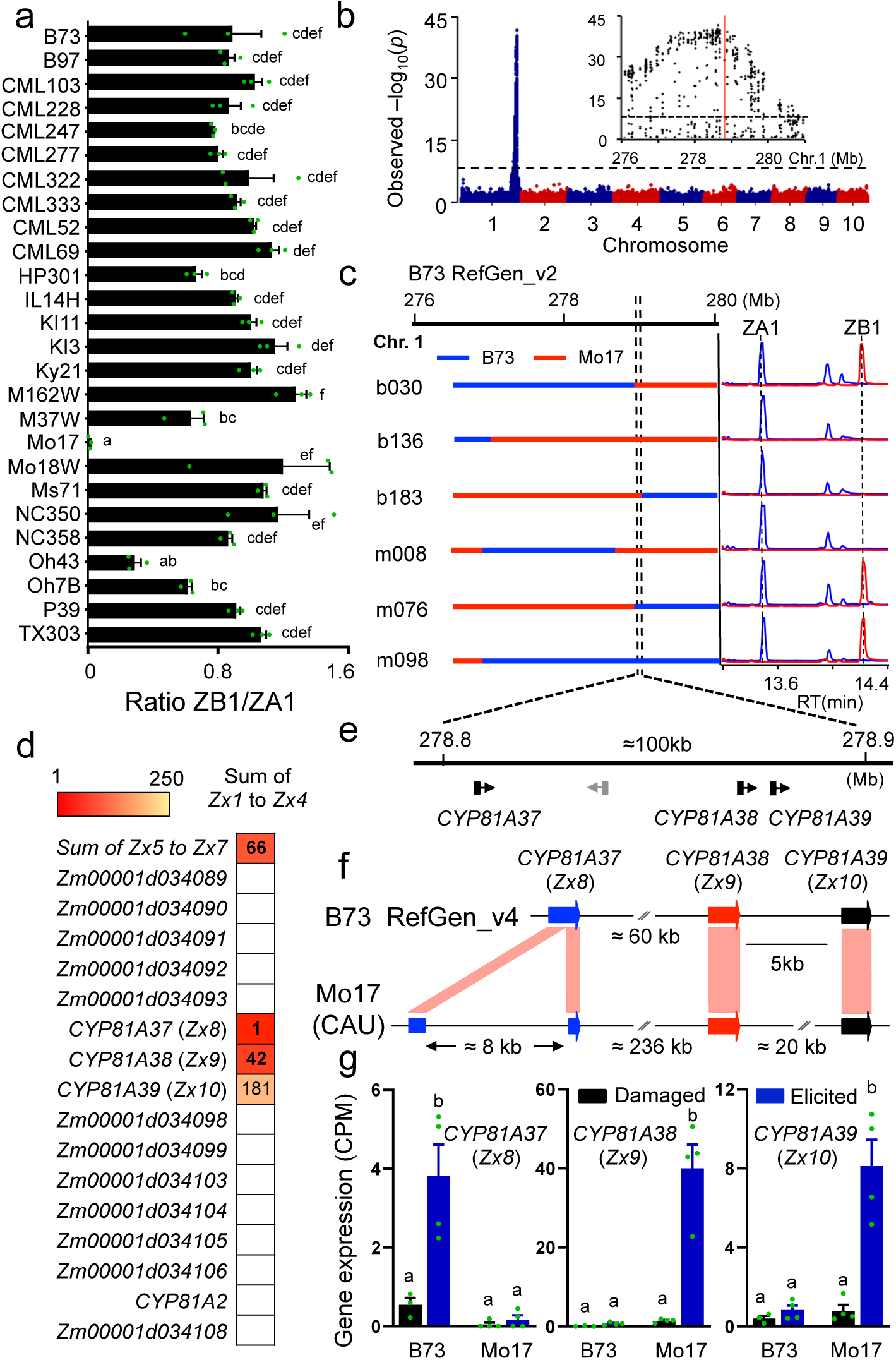
Association mapping reveals zealexin gene cluster III containing three CYP81A family P450s. **a**, ZB1/ZA1 ratio in stems of diverse inbred lines treated with heat-killed *F. venenatum* for 3 days identified Mo17 as a unique parent. **b**, Association analysis of the ratio of ZB1 to ZA1 using the Intermated B73 x Mo17 (IBM) recombinant inbred lines (RILs) with the general linear model (GLM) and 173,984 single nucleotide polymorphisms (SNPs). The most statistically significant SNPs are located on chr. 1 (B73 RefGen_v2). The dashed line denotes a 5% Bonferroni correction. Insert: local Manhattan plot on chr. 1. **c**, B73 and Mo17 chromosomal segments in IBM near isogenic lines (NILs) represented by blue and red, respectively, paired with chemotypes indicated as GC-MS selected-ion chromatograms (SIC) (ZA1 = *m*/*z* 136 blue; ZB1 = *m*/*z* 246 red). **d,** Heat map depicting the co-expression of genes in the mapping region with the summed expression of *Zx1* to Zx*4* and *Zx5* to Zx*7* present in 1960 RNA-seq samples. Low numbers indicate supportive scores while weak MR correlations > 250 were omitted. **e,** The locus was fine-mapped to a 100 kb region on B73 BACs AC202436 and AC196018 on chr. 1 containing four genes. **f**, Tandem array of *CYP81A37* (*Zx8*), *CYP81A38* (*Zx9*), *CYP81A39* (*Zx10*) on chr. 1 of the B73 genome (RefGen_V4) and the Mo17 genome (China Agricultural University, CAU). Compared with B73, Mo17 *Zx8* contains an 8 kb insertion. **g**, 3’ RNA-seq results derived from B73 and Mo17 stems either Damaged or additionally treated with heat-killed *F. venenatum* (Elicited) and harvested 36 h later. Gene expression is given as Counts Per Million mapped reads (CPM). Error bars in **a** and **g** indicate mean ± s.e.m. (*n* = 3-4 biologically independent replicates). Within plots, different letters (a–f) represent significant differences (one-way ANOVA, Tukey’s corrections for multiple comparisons, *P* < 0.05)

**Fig. 4.**
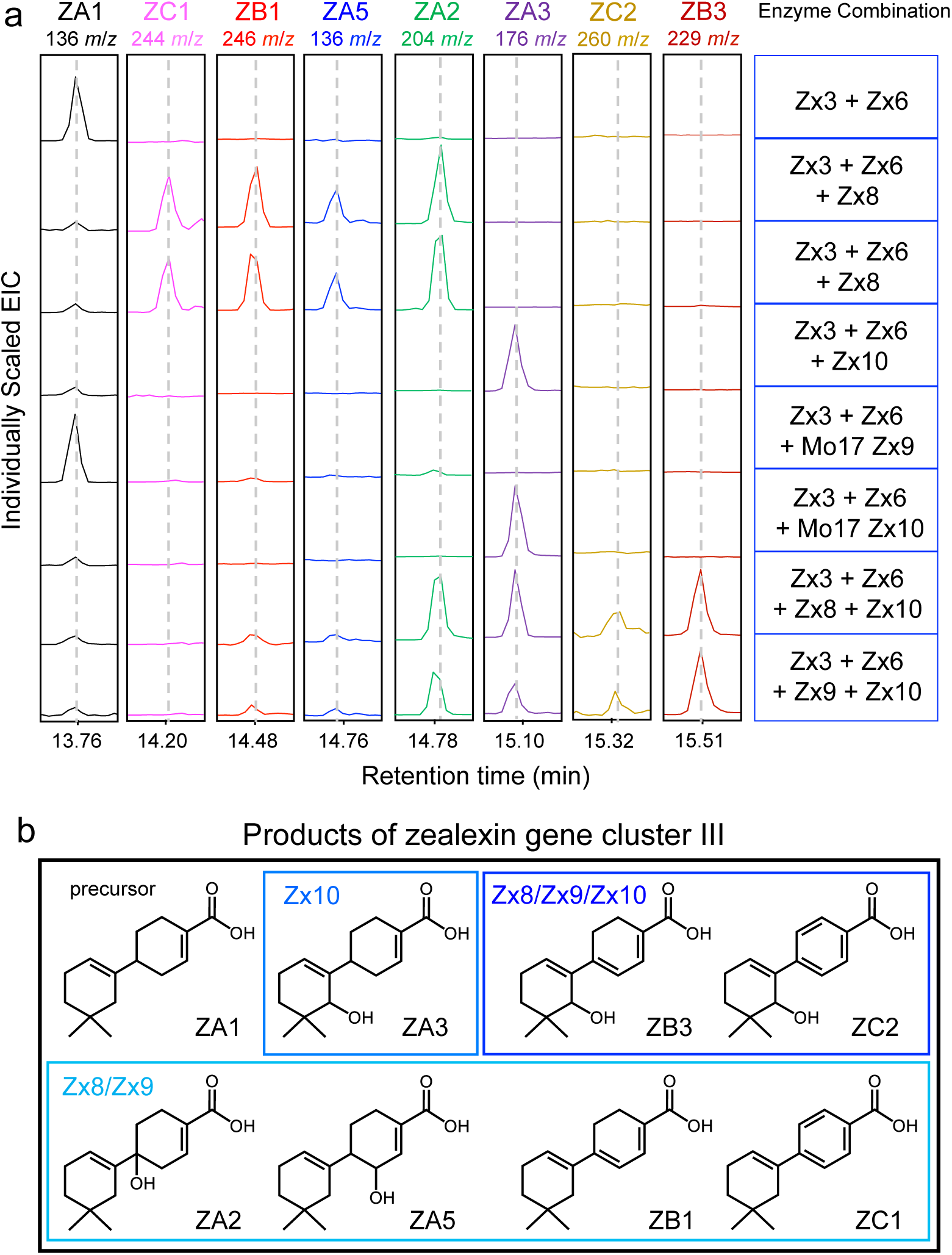
Enzyme co-expression defines the role of zealexin gene cluster III in antibiotic biosynthesis. **a**, GC-MS extracted ion chromatograms (EIC) of extracts derived from *Agrobacterium*-mediated transient *N. benthamiana* co-expression assays using representative zealexin pathway genes from gene cluster I (*Zx3*), gene cluster II (*Zx6*), and combinations from gene cluster III including *Zx8*, *Zx9*, and *Zx10*. Combinatorial *in vivo* enzyme assays in the presence of ZA1 (*m*/*z* 136, product of Zx3 and Zx6) yielded 7 additional zealexins with diagnostic EI *m*/*z* ions as follows: ZC1 (*m*/*z* 244), ZB1 (*m*/*z* 246), ZA5 (*m*/*z* 136), ZA2 (*m*/*z* 204), ZA3 (*m*/*z* 176), ZC2 (*m*/*z* 260) and ZB3 (*m*/*z* 229). ZA5 and ZB3 represent novel compounds. To address the association mapping results, functionality of Mo17 Zx9 was included and supported highly impaired activity in ZB1 synthesis. Unlike Mo17 Zx9, Mo17 Zx10 remains functional. Four independent experiments were preformed and showed similar results. **b**, Structures of zealexins derived from the activity of Zx8, Zx9 and Zx10 on ZA1 as a substrate.

**Fig. 5.**
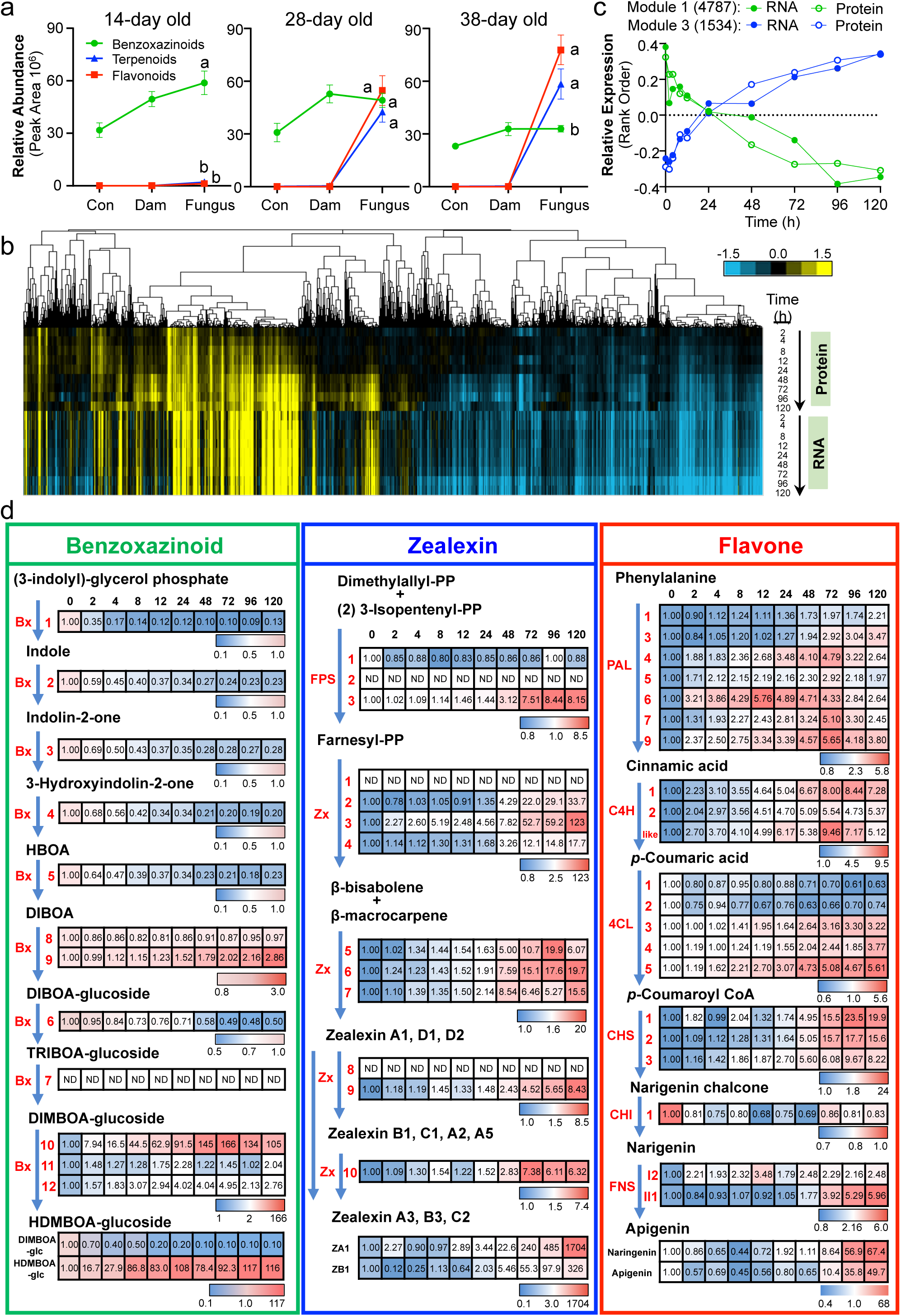
Predominant zealexin pathway activation occurs during the reprograming of fungal-induced defenses. **a**, Defense activation based on relative amounts estimated from LC-MS peak areas of major benzoxazinoids (Bx), acidic terpenoids and flavonoids present in intact maize stem tissues (control; Con) or those slit and treated with either H_2_O (damaged; Dam) or heat-killed *F. venenatum* hyphae (Fungus) at age 11, 25 and 35 days and harvested 3 days later. Error bars indicate mean ± s.e.m. (*n* = 4 biologically independent replicates). Within plots, different letters (a–b) represent significant differences for the *F. venenatum* treatment (one-way ANOVA followed by Tukey’s test corrections for multiple comparisons, *P* < 0.05). **b**, Complete linkage hierarchical clustering of 10,508 unique protein fold changes paired with mapping of log2 RNA-fold changes during a 120 h *F. venenatum* elicitation time course in W22 stems using 38 day old plants. Vertical lines correspond to individual gene IDs. Rows are organized by time point, data type and colors (blue, under-expressed; yellow, over-expressed). **c,** Weighted Gene Co-Expression Network Analysis (WGCNA) of the W22 *F. venenatum* elicitation time course proteomic and transcriptomic data identifies modules with distinct regulation patterns. Module 1 (4787 gene IDs) includes early steps in benzoxazinoid (Bx) biosynthesis while module 3 (1534 gene IDs) contains the Zx pathway. **d,** Heat maps of normalized protein fold changes in the W22 stem *F. venenatum* elicitation time course for benzoxazinoid, zealexin and flavone pathways. Corresponding metabolite fold changes for representative benzoxazinoids [2,4-dihydroxy-7-methoxy-1,4-benzoxazin-3-one glucoside (DIMBOA-glc); 2-hydroxy-4,7-dimethoxy-1,4-benzoxazin-3-one-Glc (HDMBOA-glc) and flavone pathway metabolites (naringenin, apigenin) were analyzed by LC-MS while zealexins (ZA1, ZB1) were analyzed by GC-MS. B73 RefGen_V4 gene IDs and abbreviations are defined in Supplemental Table 2).

To understand common mutations causing a loss of function in Zx1 to Zx4, we examined amino acid (AA) sequence variations in W22 Zx1 and observed a W274R substitution predicted to negatively impact the catalytic site (Supplementary Fig. 5)^38^. Reversion of inactive W22 Zx1 back to R274W or mutation of active W22 Zx4 (R274) to W274R respectively reactivated and inactivated the enzymes in *N. benthamiana* in transient expression assays (Supplementary Fig. 6). W22-like Zx1 non-synonymous SNPs at chromosome 10 position 56448050 (A to G) underlying the Zx1 W274R null mutation are common in maize germplasm and present in >10% of examined inbreds (Supplementary Table 5)^39, 40^. To demonstrate endogenous relationships, mutual rank (MR)-based global gene co-expression analyses were used to associate transcriptional patterns of maize sesquiterpene synthases in a large RNA-seq dataset^40^. Analyses revealed the highest degree of co-regulation between *Zx1*/*Zx3* and *Zx2*/*Zx4* with partial *ZmTPS21* co-regulation responsible for β-selinene derived antibiotics (Fig. 1h)^26^. Genome-wide analyses of *Zx1* to *Zx4* expression levels in diverse inbreds are consistent with the complex patterns (Fig. 1i) witnessed in B73, Mo17 and W22 (Fig. 1c-e). Functional analyses of *Zx* gene cluster I demonstrate that inbred specific combinations of functional Zx1 to Zx4 proteins contribute to β-macrocarpene and β-bisabolene production (Fig. 1a-i).

### A second zealexin pathway gene cluster contains three promiscuous CYP71Z family cytochrome P450s

Increases in *Zx1* to *Zx4* accumulation are among the largest fold transcriptional changes following pathogen challenge^29, 30^. In an early analysis of stems, we observed a member of the cytochrome P450 CYP71 family, *ZmCYP71Z18* (NM_001147894), to be among the most *F. graminearum* up-regulated genes that co-occurred with *Zx1* to *Zx4* members^29^. Subsequently, we demonstrated that both ZmCYP71Z18 and the adjoining ZmCYP71Z16 are catalytically active in the oxidation of β-macrocarpene to zealexin A1^27^. To consider roles for additional P450s, we performed a global gene co-expression analysis of the summed expression *Zx1* to *Zx4* with all predicted maize *P450* transcripts and revealed nine candidates with low MR scores (<250) including *ZmCYP71Z18* and *ZmCYP71Z19* (Fig. 2a) that were further supported by replicated RNA-seq data (Supplementary Table 2). *ZmCYP71Z19* is phylogenetically most closely related to *ZmCYP71Z16/18*^18^ and is located within the same 15 gene interval on chromosome 5 (Fig. 2b; Supplementary Fig. 7). Like *Zx1* to *Zx4*, *ZmCYP71Z16/18/19* each display variable relative expression between inbreds (Fig. 2c). We name the B73 P450 genes *ZmCYP71Z19* (*Zm00001d014121*) *Zx5*, *ZmCYP71Z18* (Zm00001d014134) *Zx6* and *ZmCYP71Z16* (*Zm00001d014136*) *Zx7* based on chromosome order with each sharing >71% protein sequence identity (Supplementary Fig. 8). Consistent with a shared role in zealexin biosynthesis, *Agrobacterium*-mediated enzyme co-expression assays with B73 Zx3 in *N. benthamiana* demonstrate that Zx5, Zx6 and Zx7 each independently catalyze the oxidization of β-macrocarpene to zealexin A1 (ZA1; Fig. 2d and 2g) providing pathway redundancy.

As the initial product of Zx1 to Zx4 (Fig. 1f-g), β-bisabolene predictably contributes to the array of 13 established candidate zealexins^29^. Purification efforts from diseased maize sheath tissue enabled the isolation and NMR identification of 2 acidic β-bisabolene derivatives, namely zealexin D1 (4-(6-methylhepta-1,5-dien-2-yl)cyclohex-1-ene-1-carboxylic acid) and zealexin D2 (2-methyl-6-(4-methylcyclohex-3-en-1-yl)hepta-2,6-dienoic acid) that produce diagnostic GC/MS electron ionization (EI) spectra as methyl ester derivatives (Supplementary Table 6 and Supplementary Fig. 9). To examine the catalytic oxidation of β-bisabolene, we performed *N. benthamiana* co-expression assays using *Santalum album* monoTPS (SaMonoTPS, EU798692) which utilizes the precursor *E*/*E*-farnesyl diphosphate (FDP) to produce β-bisabolene^41^. Similar to ZA1 biosynthesis, Zx5, Zx6 and Zx7 each catalyzed the complete oxidation of β-bisabolene at the C1 and C15 positions yet resulted in significant differences in the final ratios of zealexin D1/D2 produced (Fig. 2d-f; Supplementary Fig. 10). Our collective findings demonstrate that Zx6 and Zx7 support promiscuous catalytic activity on established Zx1 to Zx4 products (Fig. 1f-g) and also the diterpenoid defense precursors dolabradiene and *ent*-isokaurene^18, 27^. Zx5 enzyme co-expression analyses demonstrate activity on sesquiterpene olefins (Fig. 2d-e) including the substrate β-selinene to produce β-costic acid; however, no appreciable activity in kauralexin biosynthesis was observed (Supplementary Fig. 11)^18, 26^.

To consider the evolutionary origin and catalytic potential of zealexin gene cluster II, we examined the single copy *Sorghum bicolor* gene (*Sobic.001G235500*) *SbCYP71Z19* that exists as a Zx5 syntenic ortholog sharing 89% AA identity (Supplementary Fig. 7) despite at least 12 million years^42^ of phylogenetic divergence between the two genera. Both maize and sorghum gene evolution estimates (Supplementary Fig. 12) and enzyme co-expression studies, demonstrating that SbCYP71Z19 can oxidize both sesquiterpene and diterpene precursors (Supplementary Fig. 11), are consistent with the existence of a CYP71Z progenitor gene that possessed sufficient promiscuity to produce diverse terpenoid defenses prior to gene duplication and divergence in maize.

To determine if gene cluster II provides endogenous Zx pathway redundancy, we examined the W22 *zx5 Ds* insertion mutant (*dsgR102G05*) for fungal-elicited zealexins and found no measurable deficits (Supplementary Fig. 13). Of *Zx5* to *Zx7*, *Zx5* displays the highest degree of genome wide co-regulation with *Zx1* to *Zx4* (Fig. 2b) and the greatest degree of catalytic specificity towards sesquiterpene substrates (Supplementary Fig. 11). Despite signatures of specificity, the W22 *zx5* mutant supports endogenous gene cluster II redundancy enabling partially interchangeable enzymes to be shared by at least four different maize defense pathways^18, 27^.

### Forward genetics reveals a third zealexin biosynthetic gene cluster

Beyond carboxylic acid derivatives, zealexins contain additional oxidations, desaturations and aromatized variants^29^. To identify enzyme(s) responsible for these modifications, we screened established biparental mapping lines^43, 44^ for significant differences in the fungal-elicited ratios of ZB1 to ZA1. Compared to other examined inbreds, Mo17 uniquely displays low ZB1/ZA1 ratios (Fig. 3a). Using the Intermated B73 x Mo17 (IBM)-recombinant inbred lines (RILs)^43^, we utilized the ratio of ZB1/ZA1 as an association mapping trait in mature field roots and identified highly significant SNPs on chromosome 1 (Fig. 3b, Supplementary Table 7). Similarly, a genome wide association study (GWAS) using the Goodman association panel^39^ further supported co-localized SNPs (Supplementary Fig. 14) spanning the same interval.

To systematically narrow candidates, IBM near isogenic lines (NILs)^45^ were used for fine mapping and resulted in a narrow 100-kb region containing three B73 *CYP81A* genes (Fig. 3c-e) named *ZmCYP81A37* (*Zm00001d034095*) *Zx8*, *ZmCYP81A38* (*Zm00001d034096*) *Zx9* and *ZmCYP81A39* (*Zm00001d034097*) *Zx10*. Mutual Rank (MR) analyses of the combined expression of *Zx1* to *Zx4* and *Zx5* to *Zx7* in relation to the mapping interval confirmed strong zealexin pathway co-expression (Fig. 3d). Unlike B73, the Mo17 genome uniquely contains an 8 kb insertion in *Zx8* (Fig. 3f) and the Mo17 *Zx8* transcript displays no fungal-elicited accumulation (Fig. 3g). In the B73 genome, Zx8 shares 99% and 72% AA identity with Zx9 and Zx10 respectively (Supplementary Fig. 15).

To examine gene functions, combinations of representative B73 pathway genes *Zx3* and *Zx6* were co-expressed in *N. benthamiana* with combinations of B73 *Zx8*, B73 *Zx9* and B73 *Zx10*. Both Zx8/9 pairings resulted in the conversion of ZA1 to ZA2, zealexin A5 (ZA5; 2-hydroxy-5’,5’-dimethyl-[1,1’-bi(cyclohexane)]-1’,3-diene-4-carboxylic acid), ZB1 and ZC1, while Zx10 yielded exclusively ZA3 (Fig. 4a-b, Supplementary Fig. 9, Supplementary Table 6). Parallel microbial co-expression of Zx3, Zx7 with B73 Zx8 and B73 Zx9 in *E. coli* also demonstrated the conversion of ZA1 to ZB1, while Zx10 similarly produced ZA3 (Supplementary Fig. 16). In *N. benthamiana,* the combined activities of B73 Zx3, Zx6, Zx8/9 and Zx10 yielded the additional additive product termed ZB3 (6’-hydroxy-5’,5’-dimethyl-[1,1’-bi(cyclohexane)]-1,1’,3-triene-4-carboxylic acid) (Fig. 4a-b, Supplementary Fig. 9, Supplementary Table 6) and low levels of the aromatic variant ZC2^29, 46^. As novel compounds ZA5 and ZB3 produce characteristic (EI) spectra (Supplementary Fig. 9 and 10) and were identified in maize following purification and NMR elucidation (Supplementary Table 6). Zealexin biosynthetic lesions in Mo17 are partially explained by a loss of transcript accumulation in Zx8 but not Zx9 (Fig. 3g). *N. benthamiana* co-expression of B73 Zx3 and Zx6 with Mo17 Zx9 resulted in equal expression by quantitative real-time PCR (qrtPCR) yet a significant >10-fold reduction in ZA2, ZA5 and ZB1 production (Fig. 4a, Supplementary Fig. 17) consistent with separate deleterious mutations in both Mo17 *Zx8* and Mo17 *Zx9*.

**Fig. 6.**
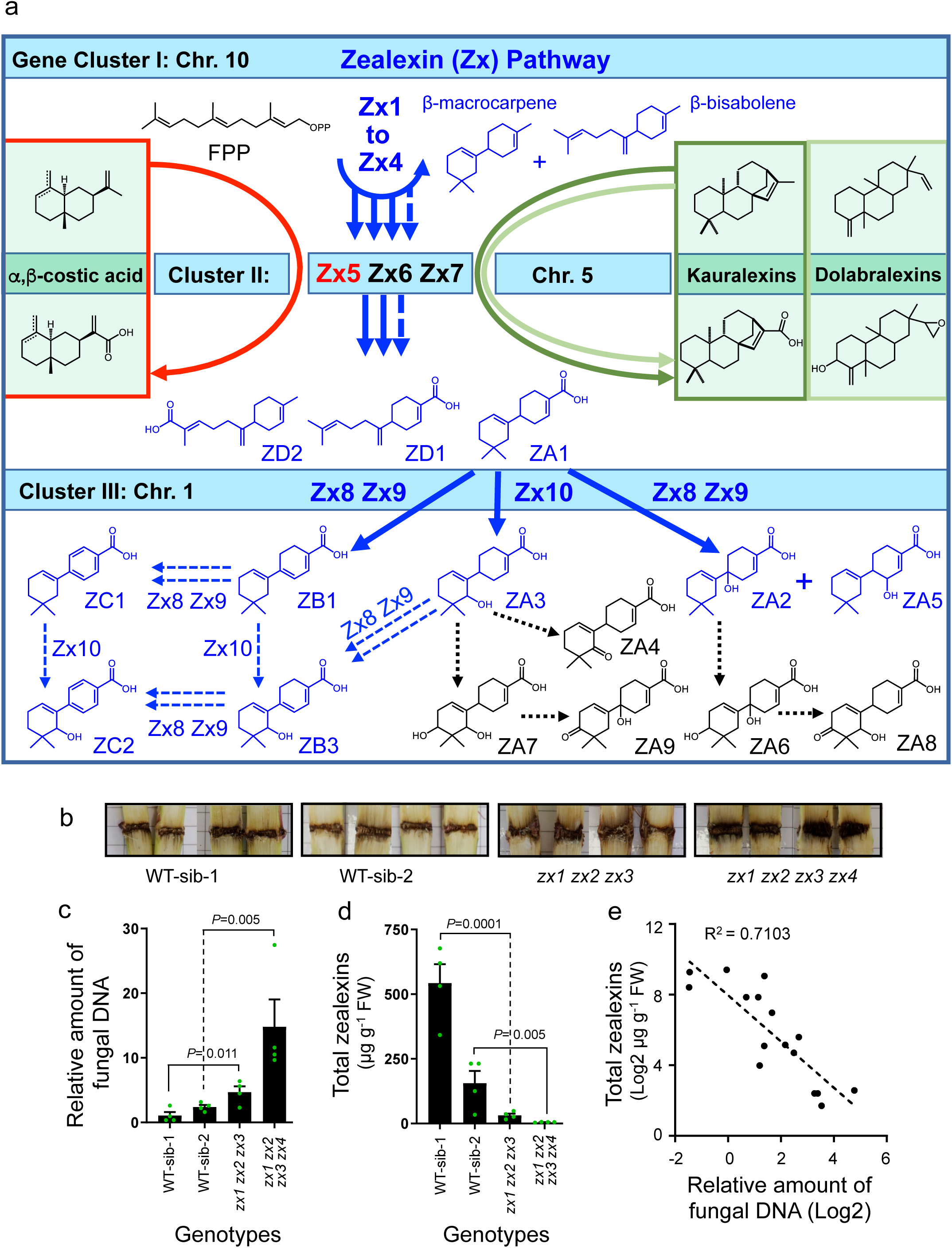
The zealexin pathway is a biosynthetic hourglass with genetic redundancy and enzyme promiscuity producing protective antibiotic cocktails. **a**, Schematic representation of enzyme activities encoded by zealexin gene cluster I (*Zx1* to *Zx4*), II (*Zx5* to *Zx7*) and III (*Zx8* to *Zx10*) including interactions with additional pathways, such as kauralexins (dark green), dolabralexins (light green) and costic acids (red). Solid blue line arrows represent enzyme catalysis supported by genetics and *in vivo* expression studies. Dashed blue line arrows indicate demonstrated activities of gene cluster III that require enzyme feeding studies to clarify the order of catalysis. Dashed black line arrows represent undefined enzyme activities. **b**, Representative disease levels in stems of CRISPR/ Cas9 derived triple (*zx1 zx2 zx3*) and quadruple (*zx1 zx2 zx3 zx4*) mutants and the respective wild type siblings (WT-sib), which were grown for 25 days and stem inoculated with *F. graminearum* (10 µl of 1.5 x 10^5^ conidia ml^-1^) for 10 days. Eight biological replicates were performed and showed similar results. **c**, Relative amount of fungal DNA in triple (*zx1 zx2 zx3*) and quadruple (*zx1 zx2 zx3 zx4*) mutants and the respective wild type siblings (WT-sib). qrtPCR was used to determine the change in relative amount of fungal DNA (*FgTri6*) in stems at 10 days after inoculation. **d**, Total zealexins (µg g^-1^ FW) present in triple (*zx1 zx2 zx3*) and quadruple (*zx1 zx2 zx3 zx4*) mutants and the respective wildtype siblings (WT-sib). **e**, Log2 correlation of endogenous zealexins with the relative amount of fungal DNA in triple (*zx1 zx2 zx3*) and quadruple (*zx1 zx2 zx3 zx4*) mutants and the respective wildtype siblings (WT-sib). Error bars in **c** and **d** indicate mean ± s.e.m. (*n* = 4 biologically independent replicates). *P* values represent Student’s *t* test, two-tailed distribution, equal variance.

To consider roles for Zx8 and Zx9, we purified ZB1 and observed significant antifungal activity against three important maize pathogens, namely *F. verticillioides*, *F. graminearum* and *Aspergillus flavus,* similar to ZA1 (Supplementary Fig. 18). Placed in context, gene cluster III containing *Zx8* to *Zx10* expands the established roles of CYP81 enzymes beyond glucosinolate, isoflavonoid, lignin and xanthone biosynthesis to now include sesquiterpenoid antibiotics (Supplementary Fig. 19).

### Elicited antibiotic production occurs during large-scale transcriptomic, proteomic and metabolomic reprogramming

For over 60 years benzoxazinoids (Bx) have been extensively examined as the predominant chemical defenses protecting maize seedlings from herbivores and pathogens^5, 15, 47, 48^. To understand the context where acidic terpenoids predominate, we applied heat-killed *F. venenatum* hyphae to wounded maize stem tissues at 13, 25 and 35 days after planting. Following three days of elicitation, 16 day-old seedlings maintained predominantly Bx metabolites while plants examined at 38 days displayed predominant complex mixtures of acidic terpenoids and flavonoids (Fig. 5a). To understand the complexity of defense activation in mature plants, we conducted a time course experiment over a period of 120 h following the application of heat-killed *Fusarium* hyphae to stem tissues of the W22 inbred and measured changes in levels of transcripts, proteins and defense metabolites (Supplementary Tables 1, 2, 8 and 9). Early (0-4 h) versus late (72-120 h) fold changes in protein levels were averaged to provide an estimate of genetic control at the transcriptional, post-transcriptional and translational levels. A combination of wounding and fungal elicitation resulted in 52% of proteins (5501 of 10508) displaying either significantly positive (2694) or negative (2807) changes in abundance following treatment (Supplementary Table 9). Protein-transcript pairs (10,508) were analyzed by complete linkage hierarchical clustering (Fig. 5b) and assigned to 11 modules (0-10) using Weighted Gene Co-Expression Network Analysis (WGCNA) of protein and RNA fold changes following rank order normalization (Fig. 5c, Supplementary Fig. 20; Supplementary Table 9). Many acidic terpenoid and flavonoid biosynthetic pathway genes grouped in module 3 (1534) containing an enrichment in gene ontology (GO) terms relating to response to stimulus, secondary and phenylpropanoid metabolic processes (Fig. 5b and d, Supplementary Table 9 and 10). Flavonoids are predominant protective biochemicals in nearly all plants, are fungal-regulated in maize and commonly co-occur with terpenoids^49–51^. The up-regulated production of simple flavonoids, such as narigenin and apigenin, are associated with increased protein levels of phenylalanine ammonia lyase (PAL), cinnamate 4-hydroxylase (C4H), 4-coumarate CoA ligase (4CL), chalcone synthase (CHS), chalcone isomerase (CHI) and flavone synthase (FNS) family members^52^ many of which parallel Zx pathway activation (Fig. 5d). Unlike terpenoid and flavonoid pathways, early Bx pathway (Bx1-5) transcripts and enzymes were assigned to Module 1 (4787) and display rapid co-suppression (Fig. 5c and 5d, Supplementary Table 2 and 10). In contrast, more terminal Bx biosynthetic enzymes, such as Bx10, are pathogen activated, exist in Module 3 and highlight a shift from general defenses to those with increased reactivity (Fig. 5d, Supplementary Table 10)^14^.

### Diverse maize antibiotics produced through an hourglass biosynthetic pathway drive pathogen resistance

To better understand zealexin diversity, we conducted large-scale stem inoculations with a necrotrophic fungal pathogen, namely southern leaf blight (SLB, *Cochliobolus heterostrophus*) and isolated additional related metabolites. Beyond known zealexins^29, 32^, ZD1-2, ZA5 and ZB3 (Fig. 2 and 4, Supplementary Table 6), we describe four additional structures namely ZA6 (1,4’-dihydroxy-5’,5’-dimethyl-[1,1’-bi(cyclohexane)]-1’,3-diene-4-carboxylic acid), ZA7 (4’,6’-dihydroxy-5’,5’-dimethyl-[1,1’-bi(cyclohexane)]-1’,3-diene-4-carboxylic acid), ZA8 (1-hydroxy-5’,5’-dimethyl-4’-oxo-[1,1’-bi(cyclohexane)]-1’,3-diene-4-carboxylic acid), and ZA9 (6’-hydroxy-5’,5’-dimethyl-4’-oxo-[1,1’-bi(cyclohexane)]-1’,3-diene-4-carboxylic acid) (Fig. 6a, Supplementary Table 6, Supplementary Fig. 9 and 21) derived from β-macrocarpene. In SLB elicitation experiments of maize stem tissues, at least 15 Zx pathway products are detectable, produce diagnostic EI spectra and significantly accumulate over time (Supplementary Figs. 9, 21-22). Our current and collective research^18, 26, 27, 29, 32^ enables construction of the maize Zx pathway and shared functions (Fig. 6a). Gene duplications resulting in gene clusters I, II and III combine with enzyme promiscuity (Fig. 1, 2 and 4; Supplementary Fig. 11) create a complex biosynthetic hourglass where diverse endogenous substrates of independent origins share enzymes. Gene cluster II (*Zx5* to *Zx7*) results in a cocktail of oxidized antibiotics further acted on by separate subsequent pathway enzymes (Fig. 6a).

Previous studies have indirectly linked β-macrocarpene synthases, zealexin production and disease resistance^29–33^. To examine endogenous relationships, we generated *zx1 zx2 zx3* triple and *zx1 zx2 zx3 zx4* quadruple insertion- and deletion-based mutants using CRISPR/Cas9 gene editing (Supplementary Fig. 23). Ten days after stalk inoculation with *F. graminearum*, *zx1 zx2 zx3 zx4* mutant plants displayed visible (Fig. 6b) and quantitative increases in disease susceptibility as estimated by the relative amount of fungal DNA (Fig. 6c). Unlike wild type plants and *zx1 zx2 zx3* plants containing a single β-macrocarpene synthase, *zx1 zx2 zx3 zx4* mutants consistently displayed a lack of detectable zealexins following *F. graminearum* inoculation (Fig. 6d) yet were not impaired in kauralexin production (Supplementary Fig. 24). Across all samples zealexin production negatively correlated (R^2^=0.71) with *F. graminearum* DNA levels (Fig. 6e). Enhanced disease susceptibility to Stewart’s wilt (*Pantoea stewartii*) bacteria was similarly observed in *zx1 zx2 zx3 zx4* mutants (Supplementary Fig. 25) demonstrating protective roles against diverse pathogens. Suppression of zealexin production in *zx1 zx2 zx3* and *zx1 zx2 zx3 zx4* mutants also significantly altered the root bacterial microbiome associated with plants grown in field soil, lowering evenness and altering the abundances of particular taxa (Supplementary Fig. 26, Supplementary Table 13). Collectively the zealexin pathway and complex array of resulting antibiotics plays a significant role in maize interactions with microorganisms.

## Discussion

An understanding of the genetic, biosynthetic and regulatory machinery controlling innate immunity is essential to optimize biochemical defenses and crop resistance traits. For insights into maize disease resistance, we leveraged multi omic approaches to elucidate hidden biochemical layers of immunity. Our current study defines ten genes present in three distinct gene clusters that ensure the production of 17 zealexin pathway metabolites with collective antibiotic action (Fig. 6a, Supplementary Table 2, Supplementary Fig. 1, 2 and 18). We demonstrate that maize antibiotic production relies on a biosynthetic hourglass pathway encoded by three ZmCYP71Z family genes (*Zx5* to *Zx7*) on chromosome 5 that contribute to multiple distinct families of sesquiterpenoids and diterpenoids. The highly interconnected nature of maize antibiotic biosynthesis demonstrates how complex combinatorial blends are biosynthesized and contribute to immunity against diverse microorganisms. Association mapping efforts to uncover genetic loci responsible for quantitative resistance to diverse maize pathogens commonly result in the discovery of multiple loci with comparatively small effects that explain 1-3% of trait variation^21, 53, 54^. Zealexin product complexity, pathway redundancy and overall resiliency to mutations are consistent with multiple disease resistance quantitative trait loci (QTLs) that are commonly too small to be detected individually^22^. Unlike qualitative resistance genes such as the wall-associated kinase (*ZmWAK*) that protects against head smut (*Sporisorium reilianum*)^55^, zealexin biosynthesis as a trait is not controlled by a single Mendelian locus. At each of the three zealexin gene clusters, pan-genome expression or sequence-level variation exists which can impair individual zealexin enzymes; however, genetic redundancies within gene clusters ensure zealexin biosynthesis.

Zealexin gene cluster I, encoded on chromosome 10, contains four tandem duplicate β-bisabolene/β-macrocarpene synthase genes, termed *Zx1* to *Zx4*. Comparative co-expression in *N. benthamiana* was used to prove pathway products (Fig. 1) and understand catalytic differences in Zx1 controlled by a single SNP common among inbreds (Supplementary Fig. 6 and Supplementary Table 5). With genetic variation driving exonic changes and observed differences in relative expression, varying functional copies of *Zx1* to *Zx4* are maintained in the maize pan-genome with inbred-specific patterns (Fig. 1a-i and Supplementary Fig. 6). Zealexin pathway redundancy contrasts both the benzoxazinoid (Bx) and kauralexin pathways, for which single gene mutations in indole-3-glycerol phosphate lyase (*benzoxazinless1*: *bx1*) and kaurene synthase-like 2 (*ksl2*) can reduce pathway metabolites to one percent of wild type levels and impair biotic stress resistance^5, 18, 56^. Zealexin pathway resiliency to single-gene mutations, coupled with the loss of pathogen resistance in *zx1 zx2 zx3 zx4* quadruple mutants, supports the hypothesis that maize relies on zealexins as key biochemical defenses.

To generate acidic non-volatile antibiotics, zealexin gene cluster II on chromosome 5 contains three neighboring duplicated CYP71Z genes (*Zx5* to *Zx7*) that each display variable co-expression with *Zx1* to *Zx4* (Fig. 2a) and drive the production of ZA1, ZD1 and ZD2. Previously associated with the synthesis of kauralexins, dolabralexins and ZA1, Zx6 and Zx7 contribute to a powerful *in vivo* system for combinatorial chemistry^18, 27^. We now demonstrate that Zx5 additionally acts on broader sesquiterpene olefins including the ZmTPS21 product β-selinene to produce β-costic acid; however, Zx5 lacks appreciable kauralexin biosynthetic activity (Supplementary Fig. 11), highlighting specific differences in the product-diversifying roles of zealexin gene cluster II. Syntenic to *Zx5*, the closely related sorghum gene *SbCYP71Z19* encodes an enzyme that similarly generates acids from β-selinene, β-bisabolene, and β-macrocarpene, demonstrating that the biosynthetic ability of related genes to oxidize diverse terpenoid precursors existed in a common ancestor of maize and sorghum (Supplementary Fig. 11 and 12). Our current systematic analyses of gene cluster II, coupled with the conserved activity encoded by *SbCYP71Z19* syntenic with *Zx5* (Supplementary Fig. 7 and 11), provides new insights into the origin of zealexins. The high degree of co-regulation between the sum expression of *Zx1* to *Zx4* with *Zx5,* comparative sesquiterpene substrate specificity yet pathway redundancy in *zx5* mutants is consistent with both a more selective role for Zx5 in zealexin biosynthesis and the maintained functional resiliency of gene cluster II to null mutations (Fig. 2a-b, Supplementary Fig. 11 and 13).

Genetic fine-mapping on chromosome 1 identified zealexin gene cluster III which unexpectedly revealed three related CYP81A family P450s. While the CYP81 subfamily have established roles in specialized metabolism surrounding glucosinolate, isoflavonoid, lignan and xanthone biosynthesis, none have been previously demonstrated to utilize terpenoid substrates (Supplementary Fig. 19, Supplementary Table 3). Zx8 and Zx9 are functionally redundant, acting on ZA1 to produce four oxidized products, namely ZA2, ZA5, ZB1 and ZC1 (Fig. 4a-b). Successive rounds of oxidation at the C6 position likely yield the observed C1-C6 desaturation in ZB1^57^. *Zx10* represents the sole non-redundant pathway gene, encoding ZmCYP81A39, which is responsible for zealexin C8 oxidation to an alcohol and the combined variants ZA3, ZB3 and ZC2 (Fig. 4a-b). Additional zealexins, namely ZA6 to ZA9, displaying C10 oxidations to alcohols and ketones, were further identified in maize tissues; however, the final enzymes responsible remain currently unknown. Collectively Zx1 to Zx10 account for production of 12 of the 17 identified zealexin pathway precursors and end products (Fig 6a). As a major product of Zx1 to Zx9 action, ZB1 exhibits significant antifungal activity at 25 μg ml^-1^ against two key *Fusarium* pathogens of maize (Supplementary Fig. 18).

Zealexin biosynthesis is a highly co-regulated pathway fully contained in WGCNA module 3 that includes 1534 transcript-protein pairs enriched for predominant gene ontology (GO) terms ’response to stimulus’ and ’secondary metabolic processes’ (Fig. 5c, Supplementary Table 10 and 11). Beyond terpenoids, phenylpropanoid defenses including naringenin chalcone, apigenin, and apigenin 7-O-methyl ether are known to accumulate in maize following anthracnose stalk rot (*Colletotrichum graminicola*) infection and reduce fungal growth^51^. Our current work demonstrates that fungal-elicited flavonoid pathway activation in maize is highly coordinated with terpenoid defenses (Fig. 5a and 5d). Multi-omic analyses place zealexin biosynthesis in the context of massive re-organization of the transcriptome and proteome including a 5-fold suppression of early Bx biosynthetic enzymes (Bx1 to Bx5) (Fig. 5d) and more generally an enrichment in GO terms defining processes surrounding ’DNA/RNA metabolism’ in module 1 (Fig. 5c, Supplementary Table 9 and 10). Together our experiments address a 15-year old hypothesis that sesquiterpenoids mediate maize disease resistance^30^, consider zealexins in the context of multiple biochemical defense pathways, and place zealexins among the predominant antibiotics contributing to defense (Fig. 5d and 6a-e).

Independent of genetic mechanisms pursued, the leveraged application of durable multiple disease resistance traits is a key goal in crop protection^22^. Efforts in sorghum have resulted in the identification of complete defense pathways, defined enzyme organization in biosynthetic metabolons, and enabled the relocation of pathways to specific organelles in heterologous plants^58, 59^. Capturing the full breadth of plant resistance traits endogenously provided by complex pathways requires an understanding of interconnections and biosynthetic nodes. Our results highlight extensive zealexin biosynthetic interactions with multiple terpenoid pathways mediated by gene cluster II that encodes three ZmCYP71Z family proteins with promiscuous activities. While the full transfer of maize terpenoid antibiotic defenses to a non-native crop model would require a series of pathway genes, leveraging single gene transfers and knowledge of enzyme promiscuity with existing modular pathways has recently provided enhanced levels of innate immunity in rice against fungal pathogens^37^.

Pathogen-elicited terpenoid antibiotics have been studied in crops for over 40 years and led to the discovery of TPS-mediated plant defenses^16, 60, 61^. Despite a massive growth of comparative omics, delineating clear connections between genotypes, chemotypes and phenotypes has remained a challenge due to genetic redundancies and enzyme promiscuity. Our use of association analyses paired with transcriptional co-regulation patterns, combinatorial biochemical studies and targeted mutant analyses using CRISPR/Cas9 has collectively provided powerful tools to narrow and interrogate metabolic pathways controlling maize innate immunity. Heterologous enzyme co-expression studies efficiently define candidate gene functions, impact of genomic variation, promiscuity, redundancy and endogenous pathway interactions leading to antibiotic complexity. Comprehensive proteomics confirms the existence of endogenous translation products and gives a more complete context to the regulation of multiple pathways known or suspected to impact defense phenotypes. Our current elucidation of highly interactive antibiotic pathways illuminates complex combinatorial strengths in the genus *Zea* which can now be considered in breeding and additional pathway engineering approaches to effectively enhance disease resistance in crops^37^.

## ONLINE METHODS

### Plant and fungal materials

Maize seeds for the Intermated B73 x Mo17 (IBM)-recombinant inbred lines (RILs)^43^ and the Goodman diversity panel^39^ were provided by Dr. Georg Jander (Boyce Thompson Institute, Ithaca, NY, USA) and Dr. Peter Balint-Kurti (U.S. Department of Agriculture-Agricultural Research Service [USDA-ARS]). Nested Association Mapping (NAM)^44^ parental line seeds were obtained from the Maize Genetic COOP Stock Center, Urbana, IL, USA. All maize lines used for genetic mapping efforts are listed (Supplementary Table 11). *Zea perennis* (Ames 21874), *Z. diploperennis* (PI 462368), *Z. luxurians* (PI 422162), *Z. m. parviglumis* (PI 384069), *Z. m. mexicana* (Ames 21851) were provided by the (USDA-ARS, North Central Regional Plant Introduction Station, Ames, IA). Maize inbreds used for replicated elicitation experiments were germinated in MetroMix 200 (Sun Gro Horticulture Distribution, Inc.) supplemented with 14-14-14 Osmocote (Scotts Miracle-Gro) and grown in a greenhouse as previously described^26^. Fungal cultures of *Fusarium graminearum* (NRRL 31084), *F. verticillioides* (NRRL 20956), *Aspergillus flavus* (NRRL 3357) and southern leaf blight (SLB; *Cochliobolus heterostrophus, C.h.*) were grown on V8 agar for 12 days before the quantification and use of spores^29^. Heat-killed *Fusarium venenatum* (strain PTA-2684) hyphae was commercially obtained (Monde Nissin Corporation Co.) and used as a non-infectious elicitor.

### Maize stem challenge with heat-killed *Fusarium* and live fungi

Using a scalpel, 35 day-old plants were slit in the center, spanning both sides of the stem, to create a 10 cm longitudinal incision. The incision wounded the upper nodes, internodes, and the most basal portion of unexpanded leaves. For replicated (n=3-4) 36 h experiments using B73 and Mo17, a ten point (n=1) 0-120 h time course with W22^18^ and the 3 day treatment of the Goodman diversity panel, approximately 500 µl of commercial heat-killed *F. venenatum* hyphae was introduced into each slit stem followed by sealing the site with clear plastic packing tape to minimize desiccation of the treated tissues. B73 and Mo17 experiments included parallel wound control plants lacking fungal hyphae treatment. For the quantification of zealexin diversity following *C. heterostrophus* inoculation, maize (*Z. mays* var. Golden Queen) plants were wounded as described above and treated with either 100 µl of H_2_O or an aqueous *C. heterostrophus* spore (1 × 10^7^ ml^−1^) suspension. Four damage controls and 4 *C. heterostrophus* treated plants were harvested each day for 3 consecutive days. For the stalk rot resistance assay, a 1-mm-diameter hole was created through the second aboveground node in the stalk of 35 day old plants and inoculated with either 10 μl of H_2_O or 10 μl of a *F. graminearum* spore (1.5 × 10^5^ ml^−1^) suspension. After 10 d, stems were longitudinally slit with a scalpel, photographed and harvested using pool of 2 individual plants for each of the 4 final harvested replicates. Within each experiment, treated maize stem tissues were harvested into liquid N_2_ at specific time points as indicated.

### Pantoea stewartii resistance assay

*P. stewartii* subsp. *stewartii* strain DC283 harboring the plasmid pHC60 encoding GFP S65T (DC283-GFP; nalR and tetR) was used as described^62^. Nalidixic acid (30 μg ml^-1^) and tetracycline (20 μg ml^-1^) were used for selection of DC283-GFP when grown in Luria-Bertani (LB) agar and LB broth at 28°C. Bacteria were subcultured by diluting 1:10 into 10 ml final volume with antibiotics and grown to an OD_600_ of 0.7. Bacteria were harvested by centrifugation at 2,800 x g for 10 min and re-suspended in phosphate buffered saline that included 0.01% Tween 20 (PBST buffer) three times. Final bacterial OD was adjusted to OD_600_ of 0.2 and used for infiltration. Twelve-day old maize seedlings were punctured with a 1 mm diameter needle in the internode between aboveground node 1 and node 2 and infiltrated with 10 µl PBS buffer (mock) or 10 µl *P. stewartii*. Plants were evaluated either after five days for bacterial growth by GFP-quantification or for wilting symptoms 16 dpi by counting the number of dying leaves per plant due to bacterial wilting. Progression of *P. stewartii*-GFP bacteria in veins was visualized by illumination with blue light using a Dark Reader Spot Lamp (DRSL; Clare Chemical Research, Dolores, CO) as previously described^63^. For quantification of *P. stewartii*-GFP, total protein from infected leaf tissue was extracted in PBST buffer. GFP fluorescence intensity was measured with Synergy H1 Multi-Mode microplate reader (BioTek, Winooski, VT) equipped with a green filter cube (excitation 485 /20 nm, emission 528 /20 nm) using total protein extract from mock-inoculated plants as blank^63^. GFP fluorescence intensity was normalized to the highest fluorescence value and is shown as relative fluorescence units (RFU). For each extraction two technical replicates were averaged and used for calculation of RFU.

### RNA-seq analyses of fungal-elicited genes

To examine B73 and Mo17 defense transcript changes following elicitation with heat-killed *Fusarium* hyphae, total RNA was isolated with the NucleoSpin^®^ RNA Plant Kit (Takara Bio USA) according to the manufacturer’s protocol. RNA quality was assessed based on RNA integrity number (RIN) using an Agilent Bioanalyzer. 3’ RNA-seq library construction and sequencing were performed at Cornell University’s Genomics Facility at the Institute of Biotechnology (Ithaca, NY, USA; http://www.biotech.cornell.edu/brc/genomics-facility/services). Approximately 500 ng of total RNA was used to construct the 3’ RNA-Seq libraries using the QuantSeq 3’ mRNA-Seq Library Prep Kit FWD (Lexogen, USA) according to the manufacturer’s instructions. All libraries, each with their own unique adapter sequences, were pooled together and sequenced on one lane of an Illumina NextSeq 500 to generate 90 bp single-end reads. Trimmomatic (v0.39) was used to remove Illumina Truseq adaptor sequences and trim the first 12 bp^64^. Trimmed reads were aligned to the maize B73 V4 reference genome (ensemble 4.44) using Hisat2 (v2.0.0)^65^ and sorted using Sambamba (v0.6.8)^66^. Raw mapped reads were quantified using featureCounts (v1.6.4)^67^. Counts were processed in R with DESeq2 to generate normalized read counts using the default method and transformation of the normalized counts using the rlogTransformation method^68^.

For the analyses of W22 tissues, RNA-seq library construction and sequencing were performed by Novogene Corporation Inc. (Sacramento, CA, USA). The mRNA was first enriched from total RNA using oligo (dT) magnetic beads and then fragmented randomly into short sequences followed by first-strand cDNA synthesis with random hexamer-primed reverse transcription. Second-strand cDNA synthesis was done by nick-translation using RNaseH and DNA polymerase I. After adaptor end-repair and ligation, cDNA was amplified via PCR and purified to create the final cDNA library. cDNA concentration was quantified using a Qubit 2.0 fluorometer (Life Technologies) and then diluted to 1 ng µl^-1^ before assessing insert size on an Agilent Bioanalyzer 2100. Library preparations were sequenced on an Illumina platform and paired-end reads were obtained. Image analysis and base calling were performed with the standard Illumina pipeline. Raw reads were filtered to remove reads containing adapters or reads of low quality. Qualified reads were then aligned to *Zea mays* AGPv4 reference genome using TopHat v2.0.12^69^. Gene expression values calculated as fragments per kilo base per million reads (FPKM) were analyzed using HTSeq v0.6.1^70^. RNA-seq data was deposited in the NCBI Gene Expression Omnibus (GEO; http://www.ncbi.nlm.nih.gov/geo/) and is accessible through accession numbers GSE138961 (W22) and GSE138962 (B73 and Mo17)

### 5’ RACE cDNA library construction and cloning of Zx pathway cDNAs

Total RNA was isolated from 35-day-old B73, Mo17 and W22 meristem tissues elicited with heat-killed *F. venenatum* hyphae collected at 48 h as described above. Approximate 2 µg total RNA was used for the construction of a 5’ rapid amplification of cDNA ends (RACE) cDNA library with the SMARTer RACE 5’/3’ Kit (Clontech) in accordance with the manufacturer’s protocol. Genes with full-length open reading frames (ORFs) were amplified using gene-specific oligonucleotides (Supplementary Table 12). For *Agrobacterium*-mediated transient expression in *N. benthamiana*, full-length ORFs, including B73 *Zx3, B73 Zx4*, W22 *Zx3,* W22 Zx*4*, B73 *Zx5*, B73 *Zx7*, Mo17 *Zx9, and* Mo17 *Zx10* were amplified from cDNA library and cloned into the expression vector pLIFE33. Genes, including B73 *Zx1 B73 Zx2*, W22 *Zx1,* W22 *Zx2*, B73 *Zx6*, and Sorghum homolog of maize *Zx5* (*Sobic.001G235500*) were synthesized and subcloned into pLIFE33. In addition, *SaMonoTPS* (*Santalum album*, EU798692) on the plasmid pESC Leu2d was subcloned into pLIFE33^20^. Native and synthetic gene sequences used in this study for enzyme characterization are detailed (Supplementary Table 4).

### Transient co-expression assays in *N. benthamiana*

For transient expression in *N. benthamiana*, pLIFE33 constructs carrying individual target genes and pEarleyGate100 with *ElHMGR*^159–582^ construct^35^ were electroporated into *Agrobacterium tumefaciens* strain GV3101. To ensure detectable production of sesquiterpenoid pathway products, all assays utilized co-expression of the coding sequence for truncated cytosolic *Euphorbia lathyris* 3-hydroxy-3-methylglutaryl-coenzyme A reductase (HMGR; ElHMGR^159–582^, JQ694150.1)^71^. An *A. tumefaciens* strain encoding the P19 protein was also equally added in order to suppress host gene silencing. *Agrobacterium* cultures were separately prepared at OD_600_ of 0.8 in 10 mM MES pH 5.6, 10 mM MgCl_2_, mixed together in equal proportion, and then infiltrated into the newly fully expanded leaves of six week old *N. benthamiana* plants using a needleless syringe^72^. Three days post infiltration (dpi), sesquiterpene volatiles from *Agrobacterium*-inoculated tobacco leaves were collected by passing purified air over the samples at 600 ml min^-1^ and trapped on inert filters containing 50 mg of HayeSep Q (80- to 100-μm mesh) polymer adsorbent (Sigma-Aldrich). Individual samples were then eluted with 150 µl of methylene chloride and analyzed by GC/EI-MS. For analyzing non-volatile sesquiterpenoids, *Agrobacterium*-inoculated leaves were harvested at 5 dpi for further metabolite analysis.

### Co-expression of TPSs and P450s in *E. coli*

Microbial co-expression of TPS and P450 enzymes was conducted using an established *E. coli* system engineered for enhanced terpenoid production^73, 74^. For functional analysis, an N-terminally truncated Zx3 gene (lacking the predicted plastid transit peptide) and a full-length gene of the maize farnesyl diphosphate synthase (*ZmFPS3*, *Zm0001d043727*) were inserted into the pCOLA-Duet1 expression vector (EMD Millipore) to generate the construct. For co-expression of the P450s Zx6, Zx7, Zx8, Zx9 and Zx10, N-terminally modified^27^ and codon-optimized genes were synthesized and subcloned into the pET-Duet1 expression vector (EMD Millipore) carrying the maize cytochrome reductase (ZmCPR2, Zm00001d026483), resulting in the constructs pET-Duet1:ZmCPR2/Zx6/Zx10. These individual constructs were then co-expressed with pCOLA-Duet1:ZmFPS3/Zx3 using the expression of pCOLA-Duet1:ZmFPPS/Zx3 only as a control. For further functional analysis of Zx7 with other P450s an additional pACYC-Duet1:ZmFPS3/Zx7 construct was generated and co-expressed with pET-Duet1:ZmCPR2/Zx8, Zx9 or Zx10. The desired construct combinations were co-transformed into *E. coli* strain BL21DE3-C41 cells (Lucigen) together with pCDFDuet:IRS for enhanced precursor formation^73^.

Cultures were grown in 50 ml Terrific Broth (TB) medium to an OD_600_ of ∼0.6 at 37°C and cooled to 16°C before protein expression was induced by adding 1 mM isopropyl-thio-galactoside (IPTG), followed by incubation for 72 h with supplement of 25 mM sodium pyruvate, 4 mg l^-1^ riboflavin, and 75 mg l^-1^ δ-aminolevulinic acid as previously described^74^. Organic solvent extraction of enzyme products was performed with 50 ml of 1:1 ethyl acetate:hexane (v/v), followed by sample concentration under N_2_ stream. Samples were resuspended in 200 µl methanol, treated with 10 µl of 1M (trimethylsilyl)diazomethane for one hour to methylate the compounds, then concentrated under N_2_ stream again. Samples were then re-suspended in 1 ml hexane for mass spectral analysis.

### Gas chromatography/mass spectrometry (GC/MS) analyses of metabolites

Maize and *N. benthamiana* tissue samples were frozen in liquid N_2_, ground to a powder and stored at -80°C until further analyses. Tissue aliquots were weighed to 50 mg, solvent extracted in a bead homogenizer, derivatized using trimethylsilyldiazomethane, and collected using vapor phase extraction as described previously^29, 75^. For metabolite extraction from *N. benthamiana*, tissue aliquots were subjected to β-glucosidase treatment (Sigma-Aldrich, Co, LLC, USA) in 250 µl 0.1 M sodium acetate buffer (pH=5.5) at a concentration of 100 units ml^-1^ at 37°C for 30 minutes before solvent extraction. GC-MS analysis was conducted using an Agilent 6890 series gas chromatograph coupled to an Agilent 5973 mass selective detector (interface temperature, 250°C; mass temperature, 150°C; source temperature, 230°C; electron energy, 70 eV). The gas chromatograph was operated with a DB-35MS column (Agilent; 30 m, 250 µm i.d., 0.25 µm film). The sample was introduced as a splitless injection with an initial oven temperature of 45°C. The temperature was held for 2.25 min, then increased to 300°C with a gradient of 20°C min-1, and held at 300°C for 5 min. Unless otherwise noted, GC/EI-MS quantification of zealexins pathway products was based on the slope of an external standard curve constructed from β-costic acid (Ark Pharm; no. AK168379) spiked into 50-mg aliquots untreated maize stem tissues identically processed using vapor phase extraction^75^. With consideration of relative retention times on a DB35 column, diagnostic EI fragments used in this study are as follows β-bisabolene (*m*/*z* 248 parent ion, *m*/*z* 93 fragment ion), β-macrocarpene (*m*/*z* 204 parent ion, *m*/*z* 136 fragment ion), ZD1 (*m*/*z* 248 parent ion, *m*/*z* 93/69 fragment ions), ZD1

(*m*/*z* 248 parent ion, *m*/*z* 93 fragment ion), ZA1 (*m*/*z* 248 parent ion, *m*/*z* 136 fragment ion), ZB1 (*m*/*z* 246 parent ion), ZA5 (*m*/*z* 136 fragment ion), ZA2 (*m*/*z* 204 fragment ion), ZA3 (*m*/*z* 176 fragment ion), ZC2 (*m*/*z* 260 parent ion), and ZB3 (*m*/*z* 229 fragment ion). To analyze complex zealexin profiles following *C.h.* inoculation, we used an isobutane-chemical ionization-GC/MS method better suited for deconvolution of co-chromatography challenges^29, 75^. In this situation analytes of interest produced the following diagnostic ions (*m*/*z*) and retention times (RT) in order; β-bisabolene [M+H]^+^ *m*/*z* 205, RT 9.81 min; β-macrocarpene [M+H] ^+^ *m*/*z* 205, RT, 9.85 min; α/β-costic acids [M+H]^+^ *m*/*z* 249, RT, 12.86 min; ZD2 [M+H]^+^ *m*/*z* 249, RT 13.14 min; ZD1 [M+H]^+^ *m*/*z* 249, RT 13.27 min; ZA1 [M+H]^+^ *m*/*z* 249, RT 13.55 min; ZC1 [M+H]^+^ *m*/*z* 245, RT 14.12, ZB1 [M+H]^+^ *m*/*z* 247, RT 14.51; ZA5 fragment [M-H_2_O]^+^ *m*/*z* 247 RT 14.88; ZA2 [M-H_2_O]^+^ *m*/*z* 247, RT 14.93; ZA3 [M-H_2_O]^+^ *m*/*z* 247, RT 15.32; ZC2 [M+H]^+^ *m*/*z* 261, RT 15.65 min; ZB3 [M+H]^+^ *m*/*z* 263, RT min 15.87; ZA4 [M+H]^+^ *m*/*z* 263, RT 16.17 min; ZA6 fragment [M-2H_2_O]^+^ *m*/*z* 245, RT 16.75; ZA7 fragment [M-2H_2_O]^+^ *m*/*z* 245, RT 17.20 min; ZA8 [M+H]^+^ *m*/*z* 279, RT 17.21 min; ZA9 [M+H]^+^ *m*/*z* 279, RT 17.48 min. Analytes were quantified based on an U-^13^C-linolenic acid internal standard as previously described^29^.

GC-MS analysis of *E. coli* expressed enzyme products was performed on an Agilent 7890B GC with a 5977 Extractor XL MS Detector at 70 eV and 1.2 ml min^-1^ He flow, using a HP5-MS column (30 m, 250 µm i.d., 0.25 µm film) with a sample volume of 1 µl and the following GC parameters: Pulsed splitless injection at 250°C and 50°C oven temperature; hold at 50°C for 3 min, 20°C min^-1^ to 300°C, hold 3 min. MS data from 90 to 600 *m*/*z* ratio were collected after a 9 min solvent delay. Product identification was conducted using authentic standards and by comparison of reference mass spectra with Wiley, National Institute of Standards and Technology and the Adams libraries.

### Liquid chromatography/mass spectrometry (LC/MS) analyses of maize benzoxazinoids, flavonoids and acidic terpenoids

LC/MS analyses were used to estimate the relative abundance of defense metabolite classes present following stem elicitation with heat killed *F. venenatum* in different aged plants. Stem tissues where ground to a fine powder with liquid N_2_ and 50 mg samples were sequentially and additively bead homogenized in 1) 100 µl 1-propanol: acetonitrile: formic acid (1:1:0.01), 2) 250 µl acetonitrile: ethyl acetate (1:1), and 3) 100 µl of H_2_O. The co-miscible acidified solvent mixture of contained 1-propanol: acetonitrile: ethyl acetate: H_2_O (11:39:28:22) which following centrifugation (15,000 rpm, 20 min) 5 µl was used for LC/MS analysis. The LC consisted of an Agilent 1260 Infinitely series HiP Degasser (G4225A), 1260 binary pump (G1312B), and a 1260 autosampler (G1329B). The binary gradient mobile phase consisted of 0.1% formic acid in H_2_O (solvent A) and 0.1% formic acid in MeOH (solvent B). Analytical samples were chromatographically separated on a Zorbax Eclipse Plus C18 Rapid Resolution HD column (Agilent: 1.8 µm, 2.1 x 50 mm) using a 0.35 ml min^-1^ flow rate. The mobile phase gradient was: 0–2 min, 5% B constant ratio; 3 min, 24% B; 28 min, 98% B, 35 min, 98% B, and 36 min 5% B for column re-equilibration before the next injection. Eluted analytes underwent electrospray ionization (ESI) via an Agilent Jet Stream Source with thermal gradient focusing using the following parameters: nozzle voltage (500 V), N_2_ nebulizing gas (flow 12 l min-1, 55 psi, 225°C) and sheath gas (350°C, 12 l min^-1^). The transfer inlet capillary was 3500V and both MS1 and MS2 heaters were at 100°C. Negative ionization mode scans (0.1 amu steps, 2.25 cycles s^-1^) from *m*/*z* 100 to 1000 were acquired. Using the conditions defined above the following retention times (min) and ions (m/z) were used to estimate relative changes in the abundance of maize defense metabolites. Benzoxazinoids included DIMBOA-Glc (split peak 5.63/6.58 min) [M-H]^-^ *m*/*z* 372; HDMBOA-Glc (split peak 7.27/7.89 min) [M+46 (formate)-H]^-^ *m*/*z* 432; and HDM_2_BOA-Glc (split peak 6.66/7.65 min) [M+46 (formate)-H]^-^ *m*/*z* 462. Flavonoids included genkwanin (18.25 min) [M-H]^-^ *m*/*z* 282; naringenin (12.98 min) [M-H]^-^ *m*/*z* 271; naringenin chalcone (13.99 min) [M-H]^-^ *m*/*z* 271; apigenin (14.94 min) [M-H]^-^ *m*/*z* 269; tetrahydroxyflavanone (9.33 min) [M-H]^-^ *m*/*z* 287; and a dimethoxytetrahydroxyflavanone candidate (split peak 10.54/13.28 min) [M-H]^-^ *m*/*z* 315. Estimates of total acidic terpenoids included A- and B-series kauralexin diterpenoids KB1 (26.59 min) [M-H]^-^ *m*/*z* 301; KA1 (26.95 min) [M-H]^-^ *m*/*z* 303; KB3 (21.13 min) [M-H]^-^ *m*/*z* 315; KA3 (22.21 min) [M-H]^-^ m/z 317, KA2 (21.14 min) [M-H]^-^ *m*/*z* 333; KA4 (21.14 min) [M-H]^-^ *m*/*z* 319 and acidic sesquiterpenoids α/β-costic acids (23.13 min) [M-H]^-^ *m*/*z* 233; ZC1 (22.60 min) [M-H]^-^ *m*/*z* 229; ZB1 (22.78 min) [M-H]^-^ *m*/*z* 231; ZD1+ZD2 (combined peak 23.45 min) [M-H]^-^ *m*/*z* 233; ZA1 (23.71 min) [M-H]^-^ *m*/*z* 233; ZB3 (17.16 min) [M-H]^-^ *m*/*z* 247; ZC2 (17.20 min) [M-H]^-^ *m*/*z* 245; ZA3 (18.00 min) [M-H]^-^ *m*/*z* 249; and ZA2 (19.59 min) [M-H]^-^ *m*/*z* 249.

### Homology modeling and Zx1 site-directed mutagenesis

A homology model of B73 β-macrocarpene synthase Zx1 was generated by using the SWISS-MODEL server(https://swissmodel.expasy.org/) based on the template for *Nicotiana tabacum* 5-*epi*-aristolochene synthase (5-EAT)^76^. Protein variants were generated by whole-plasmid PCR amplification with site-specific sense and anti-sense oligonucleotides (Supplementary Table 12), followed by Dpn I treatment to remove the parental template. All genes encoding variant proteins were sequence-verified before co-expression in *N. benthamiana*.

### Mutual Rank (MR) analyses of coregulated transcripts

The Goodman diversity panel RNA-seq dataset (B73 RefGen_V4) was composed of 300 inbred lines constituting 1960 developmentally diverse tissues samples previously deposited in the NCBI SRA project ID SRP115041^40^. Using 1960 samples, calculations of Mutual Rank (MR) were used as a measure of coexpression by calculating the geometric mean of the product of two-directional ranks derived from Pearson correlation coefficients across gene pairs^18, 77^.

### Genetic mapping of zealexin biosynthetic genes

To consider genetic variation in biparental mapping lines, we first screened the NAM parent founders, B73 and Mo17 for differences in zealexin production after 3 d of heat-killed *Fusarium* stem elicitation. Based on a selective deficit of ZB1 in Mo17, a field grown population of 216 IBM RILs^43^ was employed using naturally occurring necrotic root tissues collected 30 days after pollination for analysis of the ratio of ZA1 to ZB1 as a mapping trait. The locus responsible for zealexin B1 biosynthesis was further fine-mapped using select B73 x Mo17 NILs^45^. In effort utilize genetic diversity in a larger population, the Goodman diversity panel^39^ was grown in the greenhouse and stem tissues were harvested 3 d after elicitation with heat-killed *Fusarium*. Association analyses were conducted in TASSEL 5.0^78^ using the General Linear Model (GLM) for the IBM RILs and the unified Mixed Linear Model (MLM) to effectively control for false positives arising from the differential population structure and familial relatedness in the Goodman diversity panel^79^. Differential population structure and familial relatedness are generally not significant features in biparental RIL populations; thus, GLM analyses were selected for the IBM RILS. A list of NAM parents, IBM RILs, IBM NILs, and specific diversity panel lines used for mapping in this study are given (Supplementary Table 11). Genotypic data from imputed IBM RIL SNP markers (July 2012 All Zea GBS final build; www.panzea.org) with less than 20% missing genotypes and a >15% minor allele frequency greater were used to generate 173,984 final SNP markers. GWAS analyses utilized the B73 version 2 referenced HapMap consisting of 246,477 SNPs as described^80^. Final GWAS analyses were conducted with the R package GAPIT^81, 82^ and compressed MLM parameters to identify genomic regions putatively associated with the trait. The kinship matrix (K) was derived from the 246,477 SNPs and used jointly with population structure (Q) to improve association analysis^83^. Manhattan plots were constructed in the R package qqman (http://cran.rproject.org/web/packages/qqman)^84^.

### Gene duplication date estimation

Coding sequences for *Zea mays* (B73 RefGenv4) were fetched from Ensembl Plants and *Sorghum bicolor* (v.3.1.1) coding sequences were fetched from Phytozome V12. The coding sequences were translated to AAs with the standard translation table and aligned with clustal-omega 1.2.4^85^. Clustal-omega was run with up to 10 refinement iterations (--iterations=10) and using the full distance matrix during iterations (--full-iter). The resulting AA alignments were back-translated to nucleotides using the original coding sequences as guides. The back-translated alignments were used to estimate gene duplication dates. Date estimation was carried out using BEAST 2.6.1^86^ and a general time reversible nucleotide substitution model and a random local clock^87^. We used a calibrated Yule model as the prior for the gene tree. The maize genes were set to form a monophyletic clade in the tree, and the distribution for the common ancestor of the maize genes was set to be normal with a mean of 11.9 million years ago ^42^ and a standard deviation of 1. The MCMC routine in BEAST was run for 10 million steps, and runtimes were improved by using the beagle phylogenetics library (https://github.com/beagle-dev/beagle-lib). Trees from BEAST were visualized with DensiTree 2.2.7^88^. Scripts to perform translation, alignment, and backtranslation are available [in supplement/in GitHub/upon request]. Coding sequence alignments and BEAST XML control files for gene families are available [in supplement/in GitHub/upon request].

### Nucleic Acid Isolation and qrtPCR

Total RNA was isolated with a NucleoSpin^®^ RNA Plant Kit (Takara Bio USA) from *N. benthamiana* leaves 2 days post infiltration with the *Agrobacterium tumefaciens* strain (GV3101) according to the manufacturer’s protocol. First-strand cDNA was synthesized with SuperScript III First-Strand Synthesis SuperMix (Invitrogen, Grand Island, NY, USA). Quantitative real-time PCR (qrtPCR) was performed using Power SYBR Green Master mix (Applied Biosystems, Waltham, MA, USA), and 250 nM primers on a Bio-Rad CFX96TM Real-Time PCR Detection System. Mean cycle threshold values were normalized to the *N. benthamiana EF-1α*^89^. Fold-change calculations were performed using the equation 2^-ΔΔCt^. The sequences of qrtPCR primers used in the study are listed (Supplementary Table 12). For quantification of the fungal biomass, total DNA was extracted from fungal-inoculated maize stem tissues and subjected to qrtPCR using the *F. graminearum*-specific primers for a deoxynivalenol mycotoxin biosynthetic gene (*FgTri6*) (Supplementary Table 12)^90^. Plant DNA quantification was analyzed using specific primers (Supplementary Table 12) for the maize ribosomal protein *L17* gene (*ZmRLP17b*, *Zm00001d049815*). The relative amounts of fungal DNA were calculated by the 2^-ΔΔCt^ method, normalized to *ZmRLP17b* and expressed relative to those in damage-treated maize stems.

### *In vitro* bioassays of zealexin B1 activity as an antifungal agent

*In vitro* antifungal assays using purified ZA1 and ZB1 were performed using the Clinical and Laboratory Standards Institute M38-A2 guidelines as detailed^91^. In brief, a 96-well microtiter plate-based method using a Synergy4 (BioTech Instruments) reader was used to monitor fungal growth at 30°C in broth medium through periodic measurements of changes in optimal density (OD_600_ nm) for 48 h. Each well contained 200 μl of initial fungal inoculum (2.5 × 10^4^ conidia ml^-1^) with 1 µl of either pure dimethyl sulfoxide (DMSO) or DMSO containing 5 μg ZA1 or ZB1.

### Identification of the *Zmcyp71z19* (*zx5*) mutant

The *Dsg* insertion (dsgR102G05) in W22 *Zx5* (Zm00001d014121, B73 RefGen_V4) was verified by designing PCR primer pairs, with one gene-specific pair (Supplementary Table 12) from W22 Zx5 and one primer from the *Dsg GFP* insertion (GFP_AC-DS: TTCGCTCATGTGTTGAGCAT)^92^.

### Proteomic analysis of W22 stem tissues

As part of a previously described effort, W22 maize plants were grown individually in 1G pots for 35 days^18^. All plants were stem elicited with heat-killed with *F. venenatum* hyphae with staged timing to enable 10 time points (0, 2, 4, 8, 12, 24, 48, 72, 96, and 120 h) to be harvested within the same hour and age. Stem tissues from four plants were harvested and pooled to generate a single homogenous sample per time point, ground in liquid N_2_ and stored at -80°C. Briefly, extracted proteins were digested with Lys-C (Wako Chemicals, 125-05061) for 15 min and secondarily digested with trypsin (Roche, 03 708 969 001) for 4 h as described^18^. TMT-10 labeling was performed and checked by LC–MS/MS to confirm >99% efficiency. Labeled peptides from each time point sample were pooled together for 2D-nanoLC–MS/MS analysis as described^18^. Spectra were acquired on a Q-Exactive-HF mass spectrometer (Thermo Electron Corporation, San Jose, CA) and raw data was extracted and searched using Spectrum Mill vB.06 (Agilent Technologies)^18^. MS/MS spectra were searched against maize B73 V4 genome (Ensembl v36) with a concatenated 1:1 decoy database of 263,022 total protein sequences. Peptides shared among different protein groups were removed before quantitation. False discovery rates (FDR) were set to 0.1% at the peptide level and 1% at the protein level, respectively. In a large-scale expansion an earlier effort, we now combine analyses of 2 separate technical LC/MS replicates, assign peptide sequences to the B73 genome, include 4 new intermediate time points (8, 12, 24 and 48 h) and report 10,749 unique protein groups beyond the original 13 previously reported^18^. Raw mass spectra have been deposited and archived at the Mass Spectrometry Interactive Virtual Environment (MassIVE) repository (ftp://massive.ucsd.edu/MSV000084285).

### Analyses of paired transcriptome and proteome changes following *Fusarium* elicitation

Analyses of proteome changes utilized two technical replicates with intensity values summed between runs. In cases where the fold change at a time point for one technical replicate had ≥ 5-fold difference from the same time point for other replicate, values from the run with the higher total intensities were used. To generate a co-expression heat map of 10,508 proteins and transcripts, we performed complete linkage hierarchical clustering of the W22 fold-change protein data using Cluster 3.0 ^93^ and combined the corresponding fold-change RNA-seq values to the cluster table.

Uncentered Pearson correlations were used as the similarity metric and results were visualized in Java Treeview^94^. The W22 time course data was analyzed using the Weighted Correlation Network Analysis (WGCNA) R package^95^ to cluster genes with similarly expressed proteins and similarly expressed RNA into modules following rank ordering. One-step network construction was performed using the blockwise modules function with a high sensitivity (deep split 4)^96^. Networks and topological overlap matrices were assigned with the minimum module size set to 30 and soft thresholding power of 5. The tree cutting algorithm was adaptive-height tree cut (Dynamic Tree Cut) and average linkage hierarchical clustering was used. To find functional enrichment, we used the R package topGO^97^ (R package version 2.36.0) in conjunction with the maize-GAMER data set^98^. The R package system PipeR was used to predict maize upstream open reading frames^99^.

### Isolation and NMR identification of zealexins

Field grown maize (*Z. mays* var. Golden Queen) stems (6 kg) 20 days post pollination were harvested, slit in half lengthwise with a scalpel, inoculated with *C. heterostrophus* (SLB) hyphae and allowed to incubate for 5 d at room temperature in the dark at 100% humidity. Husks from the same plants were coated with a thin slurry of heat-killed *F. venenatum* for 5 days. Following incubation, tissues were frozen in liquid N_2_ and crushed to a coarse powder with dry ice in a hammer mill and stored at -20°C prior to extraction. Equal portions of the stem and husk tissues were combined (1 kg) and ground to a fine powder in liquid N_2_. The powder was then allowed to thaw for 2 min and further ground in 2 l of ethyl acetate. The suspension was filtered through a Buchner funnel with Whatman#1 filter paper and resulting solvent concentrated *en vacuo* on a Buchi rotoevaporator until 20 ml remained. The remaining solution was directly absorbed onto 20 g C18 resin (Discovery®, DSC-18; Sigma Aldrich) by the evaporation of residual solvent in vacuo. The resulting oil was then dry loaded and separated by preparative flash chromatography (CombiFlash^®^Rf, Teledyne ISCO, Inc, Lincoln, NE, USA) on a 5g C18 flash column (Teledyne, RediSepRf High Performance Gold). The mobile phase consisted of solvent A (acetonitrile:H_2_O, 20:80) and solvent B (acetonitrile: 100) with A held constant the 5 min followed by a linear ramp to 100% B at 60 min using a flow rate of 18 ml min^-1^ and resulted in enriched mixtures containing distinct related zealexin classes. Carboxylic acids present in fraction aliquots were derivatized with trimethylsilyldiazomethane and screened using GC/EI-MS analyses. Simple sesquiterpene acids lacking further oxygenation (ZA1, ZB1, ZD1, ZD2), those with an additional ketone or alcohol (ZA2-5, ZB3, ZC3), and those with two additional sites of oxygenation (ZA6-9) separated into 3 distinct fractions based on polarity. Each enriched flash fraction was further separated to yield pure compounds by preparative HPLC on a Dionex Ultimate 3000 instrument equipped with a YMC-Pack OD-AQ column (250 x 20mm, s-10 µm, 12 nm). Enriched flash fractions were dried under a N_2_ stream (20 mg) dissolved in 200 μl methanol and re-chromatographed using a H_2_O:ACN gradient and flow rate of 25 ml min^-1^. The steepness of the gradient employed varied dependent on the target compounds polarity. More polar compounds, such ZA6-ZA9, had shallow gradients from 100% H_2_O to 30% ACN over 45 min; whereas, the less polar zealexin D-series metabolites required a gradient of 30% ACN to 70% ACN over 45 min. Manual monitoring of ultraviolet (UV; 210 nm) signals and corresponding collection of narrow fractions enabled the final purification of previously unidentified zealexins. Structures were elucidated using ^1^H and ^13^C APT 1D NMR experiments, as well as correlated spectroscopy (COSY), heteronuclear single quantum correlation (HSQC) and heteronuclear multiple bond correlation (HMBC) 2D experiments. Additional 2D experiments were performed to help resolve overlaying signals such as nuclear Overhauser effect spectroscopy (NOESY), total correlation spectroscopy (TOCSY), HSQC-TOCSY and H2BC. NMR experiments were performed in the McKnight Brain Institute at the National High Magnetic Field Laboratory’s AMRIS Facility, which is supported by National Science Foundation Cooperative Agreement No. DMR-1157490 and the State of Florida. Purified zealexins were dissolved in chloroform-d (Cambridge Isotope Laboratories) and NMR spectra were collected on a Bruker Avance II 600-MHz cryoprobe as well as an Agilent 600-MHz ^13^C direct detect cryoprobe. Data was analyzed using Mnova (MestreLab) software. Chemical shifts were calculated by reference to chemical shifts as follows: ^1^H 7.26 ppm and 13C 77.4 ppm for CDCl_3_; ^1^H 7.16 ppm and ^13^C 128.1 ppm for benzene-d_6_; and ^1^H 1.94 ppm and ^13^C 1.4 ppm for acetonitrile-d_3_ (Supplementary Table 6). Assignments were made directly from ^1^H and ^13^C APT data when possible, or inferred through 2D experiments such as HSQC or HMBC.

### Sequence analysis and phylogenetic tree construction

Protein sequence alignments derived from UniProtKB and Genbank IDs (Supplementary Table 3) were performed using Clustal W as implemented in the BioEdit software package (http://www.mbio.ncsu.edu/BioEdit/bioedit.html). The maximum-likelihood phylogenetic trees were constructed using MEGA7 (http://www.megasoftware.net/megabeta.php) with bootstrap values based on 1,000 iterations.

### Creation of *zx1 zx2 zx3* and *zx1 zx2 zx3 zx4* mutants using CRISPR/Cas9

Zx3 guide RNA (gRNA) target site selection was based on the B73 reference genome sequence and criteria as described^100^. Flanking regions with the target site at the middle were PCR-amplified from the maize genotype Hi-II and Sanger sequenced for accuracy of genomic sequence including the gRNA complementary sequence. The gRNA gene was constructed in the intermediate vector and the expression cassette was mobilized through a gateway reaction into the Cas9-expressing binary vector for maize Hi-II transformation at the Iowa State University Plant Transformation Facility as previously described^101^. A total of ten independent T0 transgenic plants were obtained. To examine if the target gene sequence was edited, the PCR amplicons encompassing the gRNA target site (Supplementary Fig. 23) from each plant were sequenced. Early in this effort, it was revealed that *ZmTPS6/11* were part of a 4 gene cluster evident in the B73 V4 genome^102^ which reduced the frequency of complete null mutants. Ultimately, one *zx1 zx2 zx3* triple mutant and one *zx1 zx2 zx3 zx4* quadruple mutant were obtained. The homozygous mutant plants were outcrossed with B73 and the resulting F1 plants were self-pollinated to generate F2 progenies. Following genotyping, homozygous mutant plants without the CRISPR transgene were selected and backcrossed to B73. Two homozygous mutant plants, *zx1 zx2 zx3* and *zx1 zx2 zx3 zx4*, and two wild-type siblings were selected for bioassays by genotyping from self-pollinated plants after B73 backcrossing two additional times.

### Root microbiome profiling

To investigate bacterial microbiomes associated with maize zealexin knock out lines (*zx1 zx2 zx3* and *zx1 zx2 zx3 zx4*) and their corresponding wild type lines greenhouse grown plants were germinated in individual 1 G pots mixed 1:1 with commercial potting soil (BM2; American Horticultural Supply, Inc) and field soil from the UCSD Biology Field Station (La Jolla, CA) where maize has been planted each year for 3 decades. After 8 weeks, all soil was gently shaken from the roots of mature plants, and sequentially rinsed with water, 70% ethanol (30 seconds) and distilled water to preferentially remove the external rhizosphere communities.

Cleaned root tissues were then frozen in liquid N_2_, ground to a fine powder and stored at -80C. DNA was isolated from 100 mg aliquots of freeze-dried root samples using the PureLink^TM^ Plant Total DNA Purification Kit (Invitrogen) as per kit protocols. Microbiome profiling was accomplished via amplicon sequencing, using primers 515F and 806R^103^ to amplify the v4 region of bacterial 16S rRNA genes. Primers were modified with 5’ overhangs for compatibility with the MiSeq workflow, and to create a frameshifted mixture of oligos to provide signal diversity when sequencing through the primer regions. Each PCR reaction mix consisted of 0.5 U Phusion High-Fidelity DNA polymerase with associated Phusion Green HF reaction buffer (Thermo Fisher), dNTPs at 200 µM final concentration, forward and reverse primers at 0.5 µM each, peptide nucleic acid blockers ((PNA, Bio Inc) at 1 µM to prevent amplification of plastid and mitochondrial templates, 1-10 µl template DNA (depending on measured concentration) and nuclease free water to a total volume of 25 µl per reaction. Thermocycling consisted of 98 °C for 60 s, 25 cycles of (98 °C for 10 s, 75 °C for 10 s, 57 °C for 20 s, 72 °C for 15 s), final extension at 72 °C for 5 min. PCR products were cleaned using the SequalPrep Normalization Plate Kit (Thermo Fisher). An 8 cycle second round PCR was used to add sample-specific barcode indices, using the Nextera XT Index Kit (Illumina). The manufacturer’s protocol was followed, except that we substituted Phusion High-Fidelity DNA polymerase for the suggested polymerase. The sequencing library also included negative control samples (i.e., DNA extractions performed without any plant tissue, and PCRs run without any template DNA), and mock community control samples of known composition (20 Strain Staggered Mix Genomic Material; ATCC^®^ MSA-1003™, American Type Culture Collection). Indexed amplicons were cleaned and normalized with the SequalPrep kit (ThermoFisher Scientific) ahead of sample pooling. Library quality and concentration were assessed with the TapeStation instrument (Agilent) and with the Library Quantification Kit for Illumina Platforms (Kapa Biosystems). Sequencing was performed with a MiSeq instrument (Illumina), using a version 2 (500 cycle) sequencing kit. Raw sequence data are available at NCBI BioProject PRJNA580260. Amplicon sequences were processed with the DADA2 pipeline in R v.3.5^104, 105^. Briefly, primer sequences were located and trimmed using the tool Cutadapt^106^, permitting a single mismatch. Reads were culled if no primer sequence was found, when lengths <50, or when they contained ambiguous base calls. Reads were trimmed at the trailing end, where quality tends to drop (20 bases for R1, 50 bases for R2), and filtered to permit a maximum of 2 expected errors^107^. True sequence variants were inferred from the observed sequences with the DADA2 algorithm ^108^. Forward and reverse reads were merged, permitting one mismatch in the overlapping region. Chimeras were detected and removed using the DADA2 method. Sequence variants were assigned to taxonomic bins using a naïve Bayesian classifier^109^, with the Silva reference alignment v. 132^110^. Reads were culled if they could not be classified below the rank of domain, or were classified as chloroplast or mitochondria. For assessment of phylogenetic diversity, a phylogenetic tree was constructed using the package phangorn ^111^, with a neighbor-joining tree as the starting point for a maximum likelihood tree (generalized time-reversible with Gamma rate variation). Further manipulations, visualization, and analyses used the package phyloseq ^112^. Differential abundance of sequence variants was tested using the DESeq2 package for R ^113^, with taxon counts modeled on genotype [i.e., wild type vs. knock out] + locus [i.e., *zx1 zx2 zx3* vs. *zx1 zx2 zx3 zx4*). The packages ggplot2 ^114^ and pheatmap ^115^ were used for visualizations.

### Statistical analyses

Statistical analyses were conducted using JMP Pro 13.0 (SAS Institute Inc.) and GraphPad Prism 8.0 (GraphPad Software, Inc.). One-way analyses of variance (ANOVAs) were conducted to evaluate statistical differences. Tukey tests were used to correct for multiple comparisons between control and treatment groups. Student *t*-tests (unpaired, two tailed) were conducted for pairwise comparisons. *P* values < 0.05 were considered significant.

## ACKNOWLEDGMENTS

We thank A. Steinbrenner, K. Dressano, J. Chan, K. O’Leary, M. Broemmer, H. Riggleman, S. Reyes, and S. Delgado for help in planting, treatments and sampling (UCSD). Dr. Laurie Smith (UCSD) is thanked for shared UCSD Biology Field Station management. Research was supported by a grants from the USDA NIFA AFRI (1758976 to A.H. and E.S.) for sesquiterpenoids, National Science Foundation Plant-Biotic Interactions Program (grant no. 1758976 to E.S. and P.Z.) for diterpenoids, by a DOE Joint Genome Institute Community Science Program (JGI-CSP) grant (CSP2568 to P.Z., ES and AH), and by a fellowships provided by the NSF Graduate Research Fellowship Program (to KMM) and Fulbright Research Grant (E0581299; to MB).

## AUTHOR CONTRIBUTIONS

Y.D., P.W., E.P., P.Z., J.S., E.A.S. and A.H. designed the experiments and analyzed the data. Y.D., E.P., S.A.C., P.Z., K.A.K. and E.S.B. designed, performed and analyzed the transcriptome data. Y.D., E.S., A.S.K, K.M.M., P.Z., A.H. and E.A.S. performed MS experiments and MS-related metabolite data analysis. Y.D., E.S., K.M.M., P.Z., E.A.S and A.H. performed and analyzed the enzyme co-expression data. Z.S., A.T. and S.P.B. analyzed the combined proteome and transcriptome dataset. T.K. calculated estimates of gene evolution dates. D.R.N. assigned subfamily names for P450 proteins. M.M.V. and M.G.B. generated and analyzed the root microbiome data, B.Y., S.N.C. and P.W. designed gRNA constructs and generated the *zx1 zx2 zx3* and *zx1 zx2 zx3* zx4 maize mutants. J.S. and M.B. performed metabolite purifications and analyzed the NMR data. Y.D. and P.W. performed the *in vitro* and *in vivo* antibiotic resistance assays. Y.D., P.W., E.P., P.Z. E.A.S. and A.H. wrote the manuscript with input from all authors.

## References

1. Roser, H. R. a. M. Land Use. Our World in Data (2020).

2. Evenson, R. E. & Gollin, D. Assessing the impact of the green revolution, 1960 to 2000. Science 300, 758–762 (2003).

3. Cartwright, D., Langcake, P., Pryce, R. J., Leworthy, D. P. & Ride, J. P. Chemical activation of host defence mechanisms as a basis for crop protection. Nature 267, 511–513 (1977).

4. Snyder, B. A. & Nicholson, R. L. Synthesis of phytoalexins in sorghum as a site-specific response to fungal ingress. Science 248, 1637–1639, doi:10.1126/science.248.4963.1637 (1990).

5. Frey, M. et al. Analysis of a chemical plant defense mechanism in grasses. Science 277, 696–699 (1997).

6. Goswami, R. S. & Kistler, H. C. Heading for disaster: *Fusarium graminearum* on cereal crops. Molecular Plant Pathology 5, 515–525(2004).

7. Mueller, D. et al. Corn yield loss estimates due to diseases in the United States and Ontario, Canada from 2012 to 2015. Plant Health Progress 17, PHP-RS-16-0030 (2016).

8. Genetics for a warming world. Nat. Genet. 51, 1195–1195 (2019).

9. Dixon, R. A. Natural products and plant disease resistance. Nature 411, 843–847, doi:10.1038/35081178 (2001).

10. Jones, J. D. G. & Dangl, J. L. The plant immune system. Nature 444, 323–329, doi:10.1038/nature05286 (2006).

11. van Loon, L. C., Rep, M. & Pieterse, C. M. J. in Annual Review of Phytopathology Vol. 44 Annual Review of Phytopathology 135–162 (2006).

12. Moghe, G. D. & Kruse, L. H. The study of plant specialized metabolism: Challenges and prospects in the genomics era. Am. J. Bot. 105, 959–962 (2018).

13. Wouters, F. C., Blanchette, B., Gershenzon, J. & Vassao, D. G. Plant defense and herbivore counter-defense: benzoxazinoids and insect herbivores. Phytochemistry Reviews 15, 1127–1151 (2016).

14. Meihls, L. N. et al. Natural variation in maize aphid resistance is associated with 2,4-dihydroxy-7-methoxy-1,4-benzoxazin-3-one glucoside methyltransferase activity. Plant Cell 25, 2341–2355 (2013).

15. Fraenkel, G. S. The raison d’etre of secondary plant substances; these odd chemicals arose as a means of protecting plants from insects and now guide insects to food. Science 129, 1466–1470 (1959).

16. Schmelz, E. A. et al. Biosynthesis, elicitation and roles of monocot terpenoid phytoalexins. Plant J. 79, 659–678 (2014).

17. Vaughan, M. M. et al. Accumulation of terpenoid phytoalexins in maize roots is associated with drought tolerance. Plant Cell and Environment 38, 2195–2207 (2015).

18. Ding, Y. et al. Multiple genes recruited from hormone pathways partition maize diterpenoid defences. Nature Plants, doi:10.1038/s41477-019-0509-6 (2019).

19. Casas, M. I. et al. Identification and Characterization of Maize salmon silks Genes Involved in Insecticidal Maysin Biosynthesis. Plant Cell 28, 1297–1309 (2016).

20. Degenhardt, J. Indirect Defense Responses to Herbivory in Grasses. Plant Physiol. 149, 96–102 (2009).

21. Zila, C. T., Samayoa, L. F., Santiago, R., Butron, A. & Holland, J. B. A genome-wide association study reveals genes associated with Fusarium ear rot resistance in a maize core diversity panel. G3-Genes Genom Genet 3 (2013).

22. Wiesner-Hanks, T. & Nelson, R. Multiple Disease Resistance in Plants. Annu Rev Phytopathol 54, 229–252 (2016).

23. Stagnati, L. et al. A Genome Wide Association Study Reveals Markers and Genes Associated with Resistance to Fusarium verticillioides Infection of Seedlings in a Maize Diversity Panel. G3-Genes Genomes Genetics 9, 571–579 (2019).

24. Banerjee, A. & Hamberger, B. P450s controlling metabolic bifurcations in plant terpene specialized metabolism. Phytochem Rev 17, 81–111 (2018).

25. Karunanithi, P. S. & Zerbe, P. terpene synthases as metabolic gatekeepers in the evolution of plant terpenoid chemical diversity. Frontiers in Plant Science 10, doi:10.3389/fpls.2019.01166 (2019).

26. Ding, Y. Z. et al. Selinene volatiles are essential precursors for maize defense promoting fungal pathogen resistance. Plant Physiol. 175, 1455–1468 (2017).

27. Mafu, S. et al. Discovery, biosynthesis and stress-related accumulation of dolabradiene-derived defenses in maize. Plant Physiol. 176, 2677–2690 (2018).

28. Christensen, S. A. et al. Commercial hybrids and mutant genotypes reveal complex protective roles for inducible terpenoid defenses in maize. Journal of Experimental Botany 69, 1693–1705 (2018).

29. Huffaker, A. et al. Novel acidic sesquiterpenoids constitute a dominant class of pathogen-induced phytoalexins in maize. Plant Physiol. 156, 2082–2097 (2011).

30. Basse, C. W. Dissecting defense-related and developmental transcriptional responses of maize during *Ustilago maydis* infection and subsequent tumor formation. Plant Physiol. 138, 1774–1784 (2005).

31. Kollner, T. G. et al. Protonation of a neutral (*S*)-beta-bisabolene intermediate is involved in (*S*)-beta-macrocarpene formation by the maize sesquiterpene synthases TPS6 and TPS11. J. Biol. Chem. 283, 20779–20788 (2008).

32. Christensen, S. A. et al. Fungal and herbivore elicitation of the novel maize sesquiterpenoid, zealexin A4, is attenuated by elevated CO_2_. Planta 247, 863–873 (2018).

33. van der Linde, K., Kastner, C., Kumlehn, J., Kahmann, R. & Doehlemann, G. Systemic virus-induced gene silencing allows functional characterization of maize genes during biotrophic interaction with *Ustilago maydis*. New Phytologist 189, 471–483 (2011).

34. Block, A. K., Vaughan, M. M., Schmelz, E. A. & Christensen, S. A. Biosynthesis and function of terpenoid defense compounds in maize (*Zea mays*). Planta 249, 21–30 (2019).

35. Springer, N. M. et al. The maize W22 genome provides a foundation for functional genomics and transposon biology. Nat. Genet. 50, 1282–1288 (2018).

36. Mao, H., Liu, J., Ren, F., Peters, R. J. & Wang, Q. Characterization of CYP71Z18 indicates a role in maize zealexin biosynthesis. Phytochemistry 121, 4–10 (2016).

37. Shen, Q. et al. CYP71Z18 overexpression confers elevated blast resistance in transgenic rice. Plant Mol. Biol. 100, 579–589 (2019).

38. Kersten, R. D., Diedrich, J. K., Yates, J. R., 3rd & Noel, J. P. Mechanism-based post-translational modification and inactivation in terpene synthases. ACS Chem. Biol. 10, 2501–2511 (2015).

39. Flint-Garcia, S. A. et al. Maize association population: a high-resolution platform for quantitative trait locus dissection. Plant J. 44, 1054–1064 (2005).

40. Kremling, K. A. G. et al. Dysregulation of expression correlates with rare-allele burden and fitness loss in maize. Nature 555, 520–523 (2018).

41. Jones, C. G. et al. Sandalwood fragrance biosynthesis involves sesquiterpene synthases of both the terpene synthase (TPS)-a and TPS-b subfamilies, including santalene synthases. J. Biol. Chem. 286, 17445–17454 (2011).

42. Swigonova, Z. et al. Close split of sorghum and maize genome progenitors. Genome Res. 14, 1916–1923 (2004).

43. Lee, M. et al. Expanding the genetic map of maize with the intermated B73 x Mo17 (IBM) population. Plant Mol. Biol. 48, 453–461 (2002).

44. McMullen, M. D. et al. genetic properties of the maize nested association mapping population. Science 325, 737–740 (2009).

45. Eichten, S. R. et al. B73-Mo17 near-isogenic lines demonstrate dispersed structural variation in maize. Plant Physiol. 156, 1679–1690 (2011).

46. Suzuki, R., Iijima, M., Okada, Y. & Okuyama, T. Chemical constituents of the style of *Zea mays* L. with glycation inhibitory activity. Chem. Pharm. Bull. (Tokyo) 55, 153–155 (2007).

47. Ahmad, S. et al. Benzoxazinoid metabolites regulate innate immunity against aphids and fungi in maize. Plant Physiol. 157, 317–327 (2011).

48. Smissman, E. E., Lapidus, J. B. & Beck, S. D. Corn plant resistance factor. J. Org. Chem. 22, 220–220 (1957).

49. Pollastri, S. & Tattini, M. Flavonols: old compounds for old roles. Ann Bot 108, 1225–1233 (2011).

50. Lange, B. M. The evolution of plant secretory structures and emergence of terpenoid chemical diversity. Annu. Rev. Plant Biol. 66, 139–159 (2015).

51. Balmer, D., de Papajewski, D. V., Planchamp, C., Glauser, G. & Mauch-Mani, B. Induced resistance in maize is based on organ-specific defence responses. Plant J. 74, 213–225 (2013).

52. Yang, F. et al. A Maize Gene Regulatory Network for Phenolic Metabolism. Mol Plant 10, 498–515 (2017)

53. Benson, J. M., Poland, J. A., Benson, B. M., Stromberg, E. L. & Nelson, R. J. Resistance to gray leaf spot of maize: genetic architecture and mechanisms elucidated through nested association mapping and near-isogenic line analysis. PLoS Genet 11, e1005045, doi:10.1371/journal.pgen.1005045 (2015).

54. Kump, K. L. et al. Genome-wide association study of quantitative resistance to southern leaf blight in the maize nested association mapping population. Nat. Genet. 43, 163–168 (2011).

55. Zuo, W. et al. A maize wall-associated kinase confers quantitative resistance to head smut. Nat. Genet. 47, 151–157 (2015).

56. Hamilton, R. H. A corn mutant deficient in 2,4-dihydroxy-7-methoxy-1,4-benzoxazin-3-one with an altered tolerance of atrazine. Weeds 12, 27–30 (1964).

57. Meunier, B., de Visser, S. P. & Shaik, S. Mechanism of oxidation reactions catalyzed by cytochrome p450 enzymes. Chem. Rev. 104, 3947–3980 (2004).

58. Henriques de Jesus, M. P. R. et al. Tat proteins as novel thylakoid membrane anchors organize a biosynthetic pathway in chloroplasts and increase product yield 5-fold. Metab Eng 44, 108–116 (2017).

59. Laursen, T. et al. Characterization of a dynamic metabolon producing the defense compound dhurrin in sorghum. Science 354, 890–893 (2016).

60. Chappell, J. & Hahlbrock, K. transcription of plant defense genes in response to uv-light or fungal elicitor. Nature 311, 76–78 (1984).

61. Facchini, P. J. & Chappell, J. Gene family for an elicitor-induced sesquiterpene cyclase in tobacco. Proc. Natl. Acad. Sci. U. S. A. 89, 11088–11092 (1992).

62. Koutsoudis, M. D., Tsaltas, D., Minogue, T. D. & von Bodman, S. B. Quorum-sensing regulation governs bacterial adhesion, biofilm development, and host colonization in *Pantoea stewartii* subspecies stewartii. Proc. Natl. Acad. Sci. U. S. A. 103, 5983–5988 (2006).

63. Doblas-Ibanez, P. et al. Dominant, heritable resistance to Stewart’s wilt in maize is associated with an enhanced vascular defense response to infection with *Pantoea stewartii*. Mol. Plant. Microbe Interact., doi:10.1094/mpmi-05-19-0129-r (2019).

64. Bolger, A. M., Lohse, M. & Usadel, B. Trimmomatic: a flexible trimmer for Illumina sequence data. Bioinformatics 30, 2114–2120 (2014).

65. Kim, D., Landmead, B. & Salzberg, S. L. HISAT: a fast spliced aligner with low memory requirements. Nature Methods 12, 357–360 (2015).

66. Tarasov, A., Vilella, A. J., Cuppen, E., Nijman, I. J. & Prins, P. Sambamba: fast processing of NGS alignment formats. Bioinformatics 31, 2032–2034 (2015).

67. Liao, Y., Smyth, G. K. & Shi, W. FeatureCounts: an efficient general purpose program for assigning sequence reads to genomic features. Bioinformatics 30, 923–930 (2014).

68. Anders, S. & Huber, W. Differential expression analysis for sequence count data. Genome Biol. 11, R106, doi:10.1186/gb-2010-11-10-r106 (2010).

69. Kim, D. et al. TopHat2: accurate alignment of transcriptomes in the presence of insertions, deletions and gene fusions. Genome Biol. 14, doi:10.1186/gb-2013-14-4-r36 (2013).

70. Anders, S., Pyl, P. T. & Huber, W. HTSeq--a Python framework to work with high-throughput sequencing data. Bioinformatics 31, 166–169 (2015).

71. Sadre, R. et al. Cytosolic lipid droplets as engineered organelles for production and accumulation of terpenoid biomaterials in leaves. Nat Commun 10, 853, (2019).

72. Bach, S. S. et al. High-throughput testing of terpenoid biosynthesis candidate genes using transient expression in Nicotiana benthamiana. Methods Mol. Biol. 1153, 245–255 (2014).

73. Morrone, D. et al. Increasing diterpene yield with a modular metabolic engineering system in *E. coli*: comparison of MEV and MEP isoprenoid precursor pathway engineering. Appl. Microbiol. Biotechnol. 85, 1893–1906 (2010).

74. Murphy, K. M. et al. A customizable approach for the enzymatic production and purification of diterpenoid natural products. J Vis Exp, doi:10.3791/59992 (2019).

75. Schmelz, E. A., Engelberth, J., Tumlinson, J. H., Block, A. & Alborn, H. T. The use of vapor phase extraction in metabolic profiling of phytohormones and other metabolites. Plant J. 39, 790–808 (2004).

76. Starks, C. M., Back, K., Chappell, J. & Noel, J. P. Structural basis for cyclic terpene biosynthesis by tobacco 5-*epi*-aristolochene synthase. Science 277, 1815–1820 (1997).

77. Wisecaver, J. H. et al. A global co-expression network approach for connecting genes to specialized metabolic pathways in plants. The Plant Cell, 29, 944–95 (2017).

78. Bradbury, P. J. et al. TASSEL: software for association mapping of complex traits in diverse samples. Bioinformatics 23, 2633–2635 (2007).

79. Yu, J. M. et al. A unified mixed-model method for association mapping that accounts for multiple levels of relatedness. Nat. Genet. 38, 203–208 (2006).

80. Samayoa, L. F., Malvar, R. A., Olukolu, B. A., Holland, J. B. & Butron, A. Genome-wide association study reveals a set of genes associated with resistance to the Mediterranean corn borer (*Sesamia nonagrioides* L.) in a maize diversity panel. BMC Plant Biol. 15, doi:10.1186/s12870-014-0403-3 (2015).

81. Zhang, Z. W. et al. Mixed linear model approach adapted for genome-wide association studies. Nat. Genet. 42, 355–360 (2010).

82. Lipka, A. E. et al. GAPIT: genome association and prediction integrated tool. Bioinformatics 28, 2397–2399 (2012).

83. VanRaden, P. M. Efficient methods to compute genomic predictions. J. Dairy Sci. 91, 4414–4423 (2008).

84. Turner, S. D. qqman: an R package for visualizing GWAS results using Q-Q and manhattan plots. bioRxiv, doi:10.1101/005165. (2014).

85. Sievers, F. et al. Fast, scalable generation of high-quality protein multiple sequence alignments using Clustal Omega. Mol. Syst. Biol. 7, 539 (2011).

86. Bouckaert, R. et al. BEAST 2.5: An advanced software platform for Bayesian evolutionary analysis. PLoS Comput. Biol. 15, e1006650, doi:10.1371/journal.pcbi.1006650 (2019).

87. Drummond, A. J. & Suchard, M. A. Bayesian random local clocks, or one rate to rule them all. BMC Biol. 8, 114 (2010).

88. Bouckaert, R. R. DensiTree: making sense of sets of phylogenetic trees. Bioinformatics 26, 1372–1373 (2010).

89. Liu, D. et al. Validation of reference genes for gene expression studies in virus-infected *Nicotiana benthamiana* using quantitative real-time PCR. PLoS One 7, e46451 (2012).

90. Horevaj, P., Milus, E. A. & Bluhm, B. H. A real-time qPCR assay to quantify *Fusarium graminearum* biomass in wheat kernels. J. Appl. Microbiol. 111, 396–406 (2011).

91. Schmelz, E. A. et al. Identity, regulation, and activity of inducible diterpenoid phytoalexins in maize. Proc. Natl. Acad. Sci. U. S. A. 108, 5455–5460 (2011).

92. Garcia, N., Li, Y., Dooner, H. K. & Messing, J. Maize defective kernel mutant generated by insertion of a Ds element in a gene encoding a highly conserved TTI2 cochaperone. Proc. Natl. Acad. Sci. U. S. A. 114, 5165–5170 (2017).

93. de Hoon, M. J., Imoto, S., Nolan, J. & Miyano, S. Open source clustering software. Bioinformatics 20, 1453–1454 (2004).

94. Saldanha, A. J. Java Treeview--extensible visualization of microarray data. Bioinformatics 20, 3246–3248 (2004).

95. Langfelder, P. & Horvath, S. WGCNA: an R package for weighted correlation network analysis. BMC Bioinformatics 9, 559 (2008).

96. Zhang, B. & Horvath, S. A general framework for weighted gene co-expression network analysis. Stat. Appl. Genet. Mol. Biol. 4, Article17, doi:10.2202/1544-6115.1128 (2005).

97. Alexa, A. & Rahnenfuhrer, J. Gene set enrichment analysis with topGO. (2007).

98. Wimalanathan, K., Friedberg, I., Andorf, C. M. & Lawrence-Dill, C. J. Maize GO Annotation-Methods, Evaluation, and Review (maize-GAMER). Plant Direct 2, e00052, doi:10.1002/pld3.52 (2018).

99. Tw, H. B. & Girke, T. systemPipeR: NGS workflow and report generation environment. BMC Bioinformatics 17, 388 (2016).

100. Brazelton, V. A. et al. A quick guide to CRISPR sgRNA design tools. Gm Crops & Food-Biotechnology in Agriculture and the Food Chain 6, 266–276 (2015).

101. Char, S. N. et al. An *Agrobacterium*-delivered CRISPR/Cas9 system for high-frequency targeted mutagenesis in maize. Plant Biotechnology Journal 15, 257–268 (2017).

102. Jiao, Y. et al. Improved maize reference genome with single-molecule technologies. Nature 546, 524–527 (2017).

103. Caporaso, J. G. et al. Global patterns of 16S rRNA diversity at a depth of millions of sequences per sample. Proc. Natl. Acad. Sci. U. S. A. 108 **Suppl 1**, 4516–4522 (2011).

104. R Development Core Team, R. (R foundation for statistical computing Vienna, Austria, 2011).

105. Callahan, B. J., Sankaran, K., Fukuyama, J. A., McMurdie, P. J. & Holmes, S. P. Bioconductor Workflow for Microbiome Data Analysis: from raw reads to community analyses. F1000R es 5, 1492 (2016).

106. Martin, M. Cutadapt removes adapter sequences from high-throughput sequencing reads. 2011 17, 3, (2011).

107. Edgar, R. C. & Flyvbjerg, H. Error filtering, pair assembly and error correction for next-generation sequencing reads. Bioinformatics 31, 3476–3482 (2015).

108. Callahan, B. J. et al. DADA2: High-resolution sample inference from Illumina amplicon data. Nat Methods 13, 581–583 (2016).

109. Wang, Q., Garrity, G. M., Tiedje, J. M. & Cole, J. R. Naive Bayesian classifier for rapid assignment of rRNA sequences into the new bacterial taxonomy. Appl. Environ. Microbiol. 73, 5261–5267 (2007).

110. Quast, C. et al. The SILVA ribosomal RNA gene database project: improved data processing and web-based tools. Nucleic Acids Res. 41, D590–596 (2013).

111. Schliep, K. P. Phangorn: phylogenetic analysis in R. Bioinformatics 27, 592–593 (2011).

112. McMurdie, P. J. & Holmes, S. Phyloseq: an R package for reproducible interactive analysis and graphics of microbiome census data. PLoS One 8, e61217 (2013).

113. Love, M. I., Huber, W. & Anders, S. Moderated estimation of fold change and dispersion for RNA-seq data with DESeq2. Genome Biol. 15, 550 (2014).

114. Wickham, H. ggplot2: Elegant Graphics for Data Analysis. (Springer Publishing Company, Incorporated, 2009).

115. Kolde, R. Pheatmap: pretty heatmaps. R package version 61, 915 (2012).

