## Supplemental Figures 1 to 26 for "Genetic elucidation of complex biochemical traits mediating maize innate immunity"

Supplementary Fig. 1

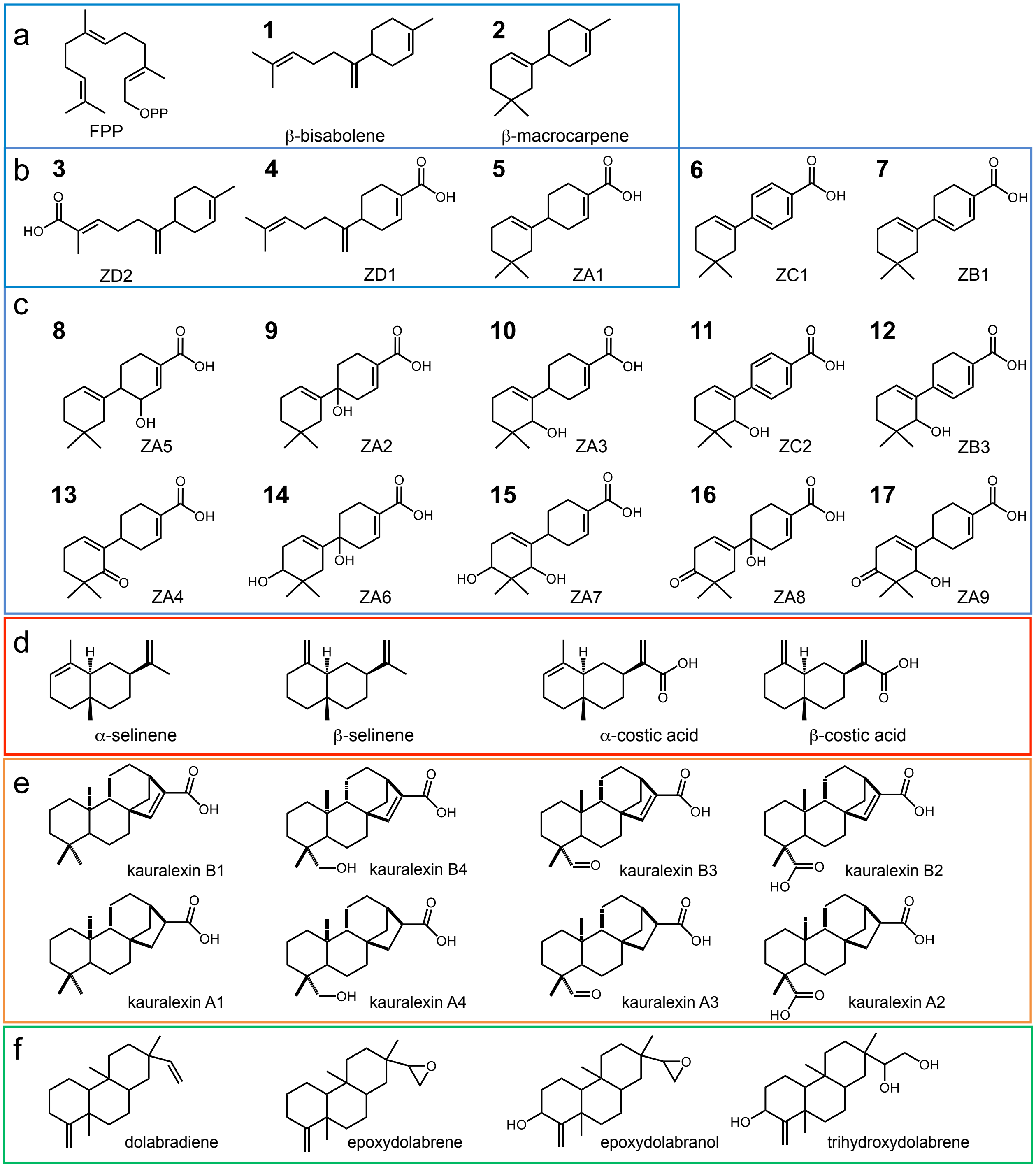


**Supplementary Fig. 1 | Maize isoprenoid precursors and products in the current study relevant to zealexin pathway elucidation. a,** Precursors for the zealexin pathway, including farnesyl diphosphate (FPP) and products of zealexin gene cluster I enzymes encoded by β-macrocarpene synthases *Zx1* to *Zx4* are (1) β-bisabolene and (2) β-macrocarpene. **b,** Zealexin gene cluster II enzymes encoded by *ZmCYP71Z19* (*Zx5*), *ZmCYP71Z18* (*Zx6*) and *ZmCYP71Z16* (*Zx7*) act on Zx1 to Zx4-derived products to produce (3) ZD2, (4) ZD1 and (5) ZA1. **c,** Zealexin gene cluster III enzymes encoded by *ZmCYP81A37* (*Zx8*), *ZmCYP81A38* (*Zx9*), *ZmCYP81A39* (*Zx10*) act on ZA1 to produce (6) ZC1, (7) ZB1, (8) ZA5, (9) ZA2, (10) ZA3, (11) ZC2 and (12) ZB3. Further modified zealexin gene cluster III products include (13) ZA4, (14) ZA6, (15) ZA7, (16) ZA8 and (17) ZA9. **d,** Products of the β-selinene synthase ZmTPS21 include α and β-selinene with further catalysis by ZmCYP71Z19 (Zx5) yielding α and β-costic acid. **e,** Representative maize *ent*-kaurene related diterpenoids utilizing the pathway genes *ZmCYP71Z18* (*Zx6*) and *ZmCYP71Z16* (*Zx7*) include B-series kauralexins (KB; KB1, KB4, KB3, KB2) and A-series kauralexins (KA; KA1, KA4, KA3, KA2). **f,** Known dolabralexin precursor dolabradiene and pathway products derived from ZmCYP71Z18 (Zx6) and ZmCYP71Z16 (Zx7) include epoxydolabrene, epoxydolabranol, and trihydroxydolabrene. **Note:** Numbered compounds (1-17) in approximate order of occurrence represent those directly considered in the present study, unnumbered compounds represent intermediates or enzyme products that interact with the zealexin pathway.

Supplementary Fig. 2

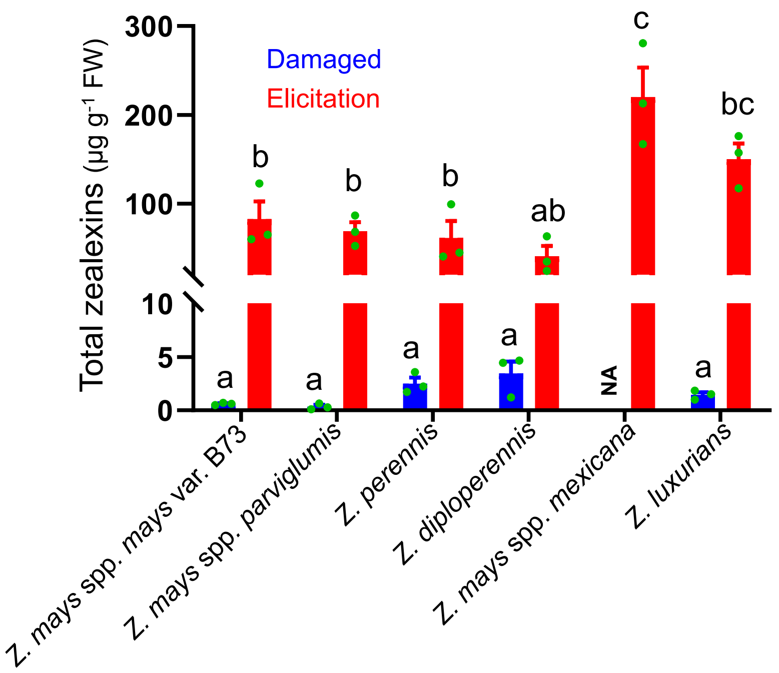


**Supplementary Fig. 2 │ Fungal-elicited zealexin accumulation occurs in all species of the genus *Zea* examined**. Average total zealexins in stems of 4-week old *Zea mays* spp. *mays* var. B73*, Zea mays* spp. *parviglumis* (AMES21889), *Zea diploperennis* (PI462368), *Zea luxurians* (PI422162), *Zea perennis* (Ames21874), and *Zea mays* spp. *mexicana* (AMES21851), treated with a heat-killed *F. venenatum* hyphae preparation. All stem tissues were harvested 5 days after treatment and analyzed by GC-MS. Total zealexins include ZA1, ZB1, ZD1 and ZD2. Error bars in indicate mean ± s.e.m. (*n* = 3 biologically independent replicates) and different letters (a–c) represent significant differences (one-way ANOVA followed by Tukey's test corrections for multiple comparisons, *P* < 0.05). NA (not available) represents missing samples/data due to low seed germination rates.

Supplementary Fig. 3

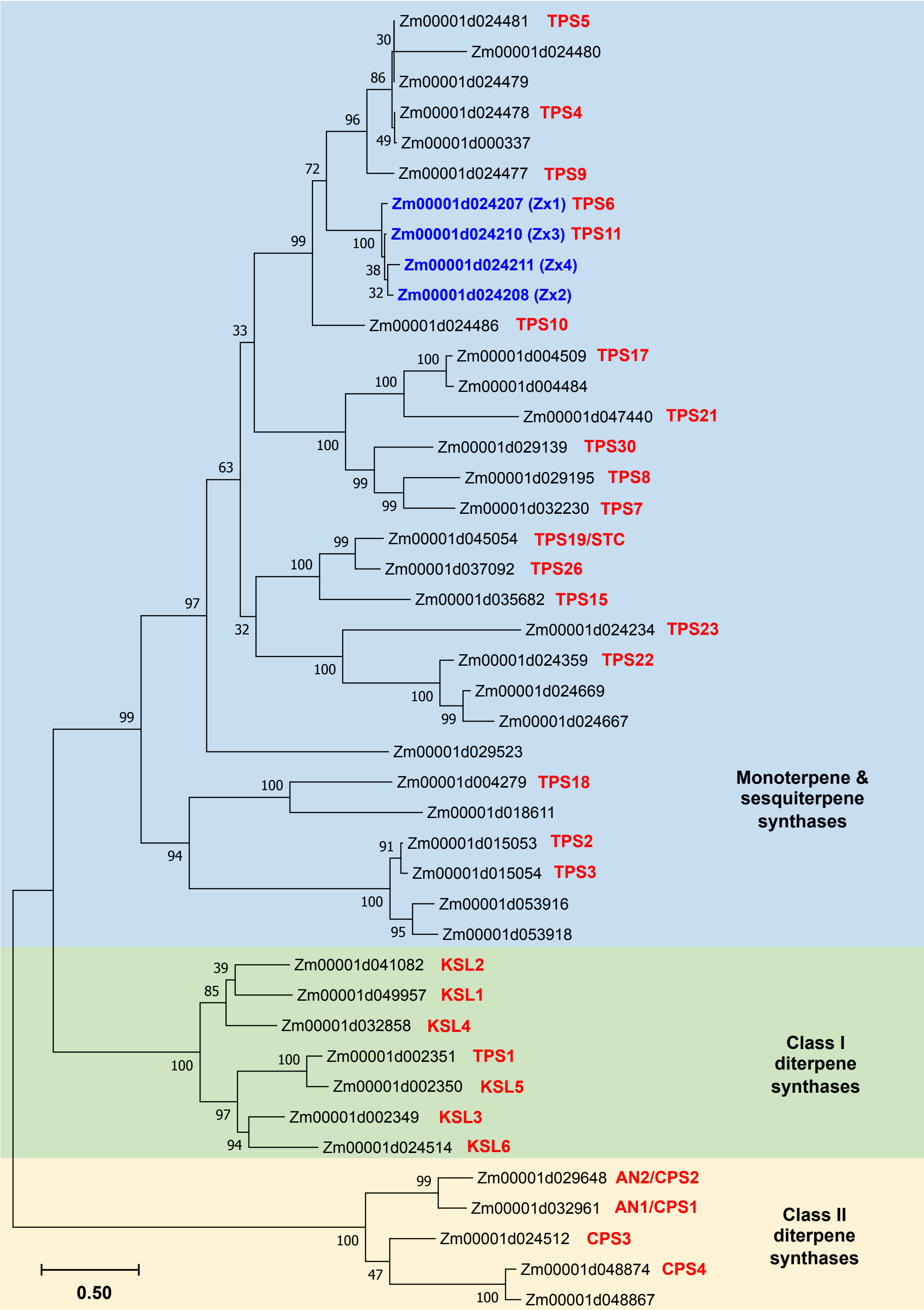


**Supplementary Fig. 3 | Maximum likelihood phylogenetic tree of 43 maize (*Zea mays*) monoterpene, sesquiterpene and diterpene synthases (B73 RefGen_V4).** Tree reconstruction was performed with the maximum likelihood algorithm using MEGA 7 program (www.megasoftware.net). Bootstrap values calculated from 1,000 iterations are indicated at the nodes. Protein sequences are listed in Supplementary Table 3

### Supplementary Fig. 4

```
B73 Zx1   MAAPTLTADGPRLGQQEMKKMSP-SFHPTLWGDFFLSYEAPTEAQEAQMREKAAVLKEEVRNMIKGSHDVPEIVDLIITL 79
B73 Zx2   MAAQTLTADGPRLGQQEMKKMSP-SFHPTLWGDFFLSYEAPTEAQEAEMRERAGVLREKVRSMIKGSHDVPEIVDLIITL 79
B73 Zx3   MAAPTLTTDGPRLGQQEMKKMSP-SFHPTLWGDFFLSYEAPTEAQEAEMRQRAEVLREEVRNMIKGSHDVPEIVDLIITL 79
B73 Zx4   MAALTQTTDGPRLGEQEMKKMSPASFHPTLWGDFFLSYEAPTETQEAQMREKAGVLKEEVRNMIKGSHDMPEIVDLIITL 80

B73 Zx1   QRLNLDYHYEDEINEKLTVVYKSNYDGGNLDLVSRRFYLLRKCGYDVSSDVFLKFKDQLGNFVEADTRSLLSLYNAAFLR 159
B73 Zx2   QRLNLDYHYEDEINEKLTVVYNSNYDGGNLVLVSRRFYLLRKCGYHVSS----------------DTRTLLSLYNAAYLR 143
B73 Zx3   QRLNLDYHYEDEINEKLAVVYNSNYDGGNLDLVSRRFYLLRKCGYHVSSDVFLNFKDQYGNFIEVDTRSLLSLYNAAYLR 159
B73 Zx4   QRLNLDYHYEDEITEKLTVVYNSNYDGGNLDLVSRQFYLLRKCGYHVSSDVFLNFKDQYGNFIGADTRSLLSLYNAAYLR 160

B73 Zx1   IHGETVLDEAISFTMRVLQDRLEHLESPLAEEVSSALDTPLFRRVGTLEMKDYIPIYEKDAKQNKSILEFAKLNFNLLQL 239
B73 Zx2   IHGETMLDEAISFTTRCLQDRLEHLESPIAEEVSSALDTPLFRRVGTLEMKDYIPIYEKDAKQNKSILEFAKLNFNLLQL 223
B73 Zx3   IHGETVLDEAISFTTRCLQDRLEHLESPIAEEVSSALDTPLFRRVGTLEMKDYIPIYEKDAKQNKSILEFAKLNFNLLQL 239
B73 Zx4   IHGETVLDEAISFTTRCLQDRLEHLESPIAEEVSSALDTPLFRRVGTLEIKDYIPIYEKDAKQNKSILEFAKLNFNLLQL 240

B73 Zx1   RYSSELKECTTWWKELRVESNLSFVRDRIVEVYFWMSGGCYDPQYSHSRIILTKIVAFITILDDTLDSHATSCESMQLAE 319
B73 Zx2   LYSSELKECTAWWKELCVESNLSFVRDRIVEVYFWMSGGCYDPQYSHSRIILTKIVAFITILDDTLDSLANSYESMQLAE 303
B73 Zx3   LYSSELKECTTWWKELRVESNLSFVRDRIVEVYFWMSGGCYDPQYSHSRIILTKIVAFITILDDTLDSHANSYESMQLAE 319
B73 Zx4   LYSSELKECTSWWKELRVESNLSFVRDRIVEVYFWMSGGCYDPQYSHSRIILTKIVAFITILDDTLDSHANSYESMQLAK 320
                                    RXR                                   DDXXD
B73 Zx1   AIERWDESAVSLLPEYMKDFYMYLLKTFSSFENELGPDKSYRVFYLKEAVKELVREYTKEIKWRDEDYVPKTLKEHLKVS 399
B73 Zx2   AVERIHEGFL--------------------------------------HVKELVREYTKEIKWRDEDYVPKTLKEHLKVS 345
B73 Zx3   AVERWDESAVSLLPEYMKDFYMYLLKTFSSFENELGPDKSYRVFYLKEAVKELVREYTKEIKWRDEDYVPKTLKEHLKVS 399
B73 Zx4   AVERWDETAVSLLPEYMKDFYMYLLKTFSSFENELGPDKSYRVFYLKEAVKELVREYTKEIKWRDEDYVPKTLKEHLKVS 400

B73 Zx1   LISIGGTLVLCSAFVGMGDVVTKKIMKWVMSDAELVKSFGIFVRLSNDIVSTKREQREKHCVSTVQCYMKQHEITMDEAC 479
B73 Zx2   LISIGGTLVLCSAFVGMGDVVTKKIMEWVMSDAELVKSFGIFVRLSNDIVSTKREQREKHCVSTVQCYMKQHELTMDEAC 425
B73 Zx3   LISIGGTLVLCSAFVGMGDVVTKKIMEWVMSDAELVKSFGIFVRLSNDIVSTKREQREKHCVSTVQCYMKQHELTMDEAC 479
B73 Zx4   LISIGGTLVLCSAFVGMGDVVTKKIMEWVMSDAELVKSFGIFVQLSNDIVSTKREQREKHCVSTVQCYMKQHELTMDEAC 480
                                                         (N,D)DXX(S,T)XXXE
B73 Zx1   EQIKELTEDSWKFMIEQGLALKEYPIIVPRTVLEFARTVDYMYKEADKYTVSHTIKDMLTSLYVKPVLM 548
B73 Zx2   EQIKELTEDSWKFMIEQGLALKEYPIIVPRTVLEFARTVDYMYKEADKYTVSHTIKDMLTSLYVKPVLM 494
B73 Zx3   EQIKELTEDSWKFMIEQGLALKEYPIIVPRTVLEFARTVDYMYKEADKYTVSHTIKDMLTSLYVKPVLM 548
B73 Zx4   EQIKELIEDSWKFMIEQGLALKEYPIIVPRTVLEFARTVDYMYKEADKYTVSHTIKDMLTSLYVKPVLM 549
```

**Supplementary Fig. 4 | Encoded amino acid sequence comparison of four B73 β-macrocarpene synthases (Zx1 to Zx4) present in *Zx* gene cluster I.** The alignment of predicted amino acid sequences encoded by B73 V4 genes *Zm00001d024207* (*Zx1*), *Zm00001d024208* (*Zx2*) *Zm00001d024210* (*Zx3*) and *Zm00001d024211* (*Zx4*) was constructed with the program MEGA7 (www.megasoftware.net) and the MUSCLE (codon) algorithm. The visualization was done with the program BIOEDIT ( http://www.mbio.ncsu.edu/BioEdit). Conserved amino acid motifs are indicated as RxR, DDxxD and (N,D)DXX(S,T)XXXE. **Note:** B73 Zx2 contains 2 deletion sites that together are predicted to negatively impact activity.

### Supplementary Fig. 5

```
W22 Zx1  MAAPTLTADGPRLGQQEMKKMSPSFHPTLWGDFFLSYEAPTEAQEAQMREKAGVLKEEVRNMIKGSHDVPEIVDLIITLQ 80
W22 Zx2  MAAQTLTADGPRLGQQE-MKMSPSFHPTLWGDFFLSYEAPTEAQEAEMRQRAEVLREEVRNMIKGSHDVPEIVDLIITLQ 79
W22 Zx3  MAAPTLTTDGPRLGQQETKKMSPSFHPTLWGDFFLSYEAPTEAQEAEMRQRAEVLREEVRNMIKGSHDVPEIVDLIITLQ 80
W22 Zx4  MAALTQTTDGPRLGEQEMKKMSPSFHPTLWGDFFLSYEAPTEAQEAQMREKAGVLKEEVRNMIKGSHDVPEIVDLIITLQ 80

W22 Zx1  RLNLDYHYEDEINEKLTVVYKSNYDGGNLDLVSRRFYLLRKCGYDVSIDVFLKFKDQLGNFVEADTRSLLSLYNAAFLRI 160
W22 Zx2  RLNLDYHYEDEINEKLTVVYNSNYDGGNLDLVSRRFYLLRKCGYHVSSDVFLNFKDQYGNFIEADTRTLLSLYNAAYLRI 159
W22 Zx3  RLNLDYHYEDEINEKLTVVYNSNYDGGNLDLVSRRFYLLRKCGYHVSSDVFLNFKDQYGNFIEADTRTLLSLYNAAYLRI 160
W22 Zx4  RLNLDYHYEDEITEKLTVVYNSNYDGGNLDLVSRRFYLLRKCGYHVSSDVFLNFKDQYGNFIEADTRSLLSLYNAAYLRI 160

W22 Zx1  HGETVLDEAISFTTRVLQDRLEHLESPLAEEVSSALDTPLFRRVGTLEMKDYIPIYEKDAKQNKSILEFAKLNFNLLQLR 240
W22 Zx2  HGETVLDEAISFTTRCLQDRLEHLESPIAEEVSSALDTPLFRRVGTLEMKDYIPIYEKDAKQNKSILEFAKLNFNLLQLL 239
W22 Zx3  HGETVLDEAISFTTRCLQDRLEHLESPIAEEVSSALDTPLFRRVGTLEMKDYIPIYEKDAKQNKLILEFAKLNFNLLQLL 240
W22 Zx4  HDETVLDEAISFTTRCLQDRLEHLESPIAEEVSSALDTPLFRRVGTLEMKDYIPIYEKDAKQNKSILEFAKLNFNLLQLL 240

W22 Zx1  YSSELKECTTWWKELRVESNLSFVRDRIVEVYFRMSGGCYDPQYSHSRIILTKIVAFITILDDTLDSHATSCESMQLAEA 320
W22 Zx2  YSSELKECTAWWKELRVESNLSFVRDRIVEVYFWMSGGCYDPQYSHSRIILTKIVAFITILDDTLDSHANSYESMQLAEA 319
W22 Zx3  YSSELKECTAWWKELRVESNLSFVRDRIVEVYFWMSGGCYDPQYSHSRIILTKIVAFITILDDTLDSHANSYESMQLAEA 320
W22 Zx4  YSSELKECTAWWKELRVESNLSFVRDRIVEVYFWMSGGCYDPQYSHSRIILTKIVAFITILDDTLDSHANSYESMQLAEA 320
                                  RXR      274                        DDXXD

W22 Zx1  IERWDESAVSLLPEYMKDFYMYLLKTFSSFENELGPDKSYRVFYLKEAVKELVREYTKEIKWRDEDYVPKTLKEHLKVSL 400
W22 Zx2  VERWDESAISLLPEYMKDFYMYLLKTFSSFENELGPDKSYRVFYLKEAVKELVREYTKEIKWRDEDYVPKTLKEHLKVSL 399
W22 Zx3  VERWDESAISLLPEYMKDFYMYLLKTFSSFENELGPDKSYRVFYLKEAVKELVREYTKEIKWRDEDYVPKTLEEHLKVSL 400
W22 Zx4  VERWDETAVSLLPEYMKDFYMYLLKTFSSFENELGPDKSYRVFYLKEAVKELVREYTKEIKWRDEDYVPKTLKEHLKVSL 400

W22 Zx1  ISIGGTLVLCSAFVGMGDVVTKKIMKWVMSDAELVKSFGIFVRLSNDIVSTKREQREKHCVSTVQCYMKQHELTMDEACE 480
W22 Zx2  ISIGGTLVLCSAFVGMGDVVTKKIMEWVMSDAELVKSFGIFVRLSNDIVSTKREQREKHCVSTVQCYMKQHELTMDEACE 479
W22 Zx3  ISIGGTLVLCSAFVGMGDVVTKKIMEWVMSDAELVKSFGIFVRLSNDIVSTKREQREKHCVSTVQCYMKQHELTMDEACE 480
W22 Zx4  ISIGGTLVLCSAFVGMGDVVTKKIMEWVMSDAELVKSFGIFVRLSNDIVSTKREQREKHCVSTVQCYMKQHELTMDEACE 480
                                                      (N,D)DXX(S,T)XXXE

W22 Zx1  QIKELTEDSWKFMIEQGLALKEYPIIVPRTVLEFARTVDYMYKEADKYTVSHTIKDMLTSLYVKPVLM 548
W22 Zx2  LIKELTEDSWKFMIEQGLALKEYPIIVPRTVLEFARTVDYMYKEADKYTVSHTIKDMLTSLYVKPVLM 547
W22 Zx3  QIKELTEDSWKFMIEQGLALKEYPIIVPRTVLEFARTVDYMYKEADKYTVSHTIKDMLTSLYVKPVLM 548
W22 Zx4  QIKELTEDSWKFMIEQGLALKEYSIIVPRTVLEFARTVDYMYKEADKYTVSHTIKDMLTSLYVKPVLM 548
```

**Supplementary Fig. 5 | Encoded amino acid sequence comparison of four W22 β-macrocarpene synthases.** The alignment of predicted amino acid sequences encoded by the W22 genes *Zm00004b038503* (*Zx1*), *Zm00004b038504* (*Zx2*), *Zm00004b038505* (*Zx3*), and *Zm00004b038506* (*Zx4*) was constructed with the program MEGA7 (www.megasoftware.net) and the MUSCLE (codon) algorithm. The visualization was done with the program BIOEDIT (http://www.mbio.ncsu.edu/BioEdit). Conserved amino acid motives are indicated as RxR, DDxxD and (N,D)DXX(S,T)XXXE. Unlike W22 Zx2 to Zx4, W22 Zx1 contains a R instead of a W at position 274 as marked with a red dashed square.

Supplementary Fig. 6

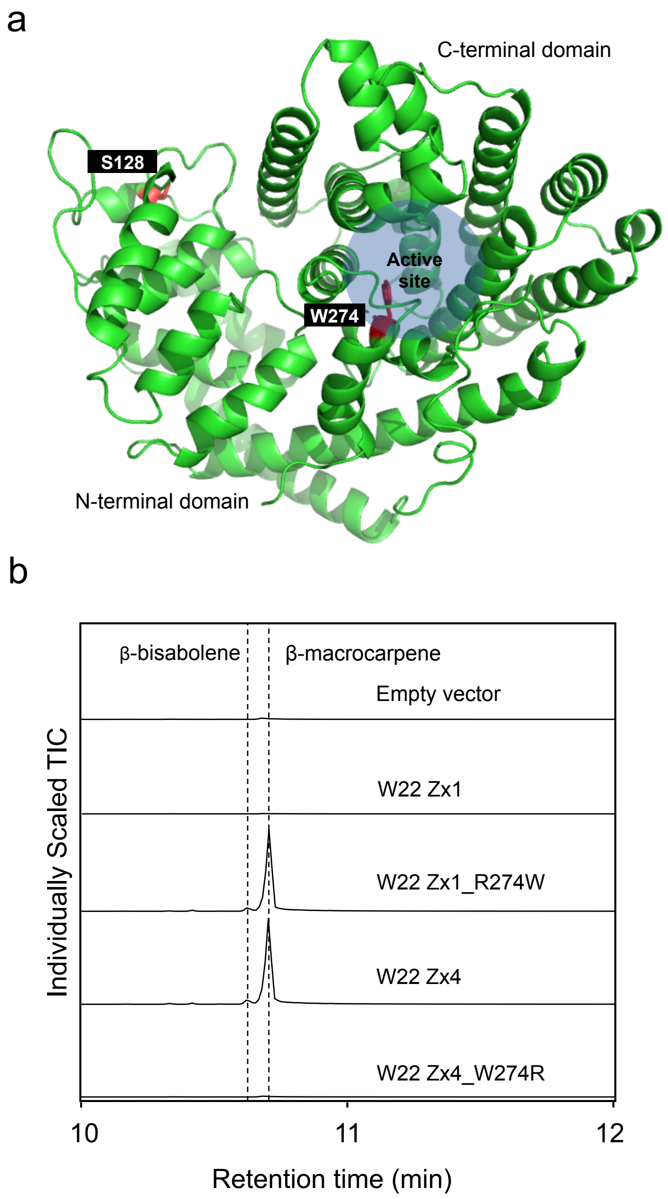


**Supplementary Fig. 6 | A single nucleotide polymorphism (SNP) commonly present in maize germplasm results in a loss of function *zx1* mutation. a**, Homology modeling of B73 β-macrocarpene synthase Zx1 based on the template 5eat.1.A (5-*epi*-aristolochene synthase from *N. tabacum*). **b**, GC-MS total ion chromatograms (TIC) are shown for the volatiles emitted from *Agrobacterium*-mediated transient expression in *N. benthamiana* leaves of the naturally occurring W22 Zx1 (Zm00004b038503) mutant, wild type W22 Zx4 (Zm00004b038506), site directed repair (R274W) of W22 Zx1 restores catalytic activity while site directed mutagenesis (W274R) of W22 Zx4 destroys catalytic activity. An empty pLife33 vector was used for the negative control. Of the analyzed maize inbreds, 35 of 237 harbor the null mutation in zx1, namely SNP_10_56448050 (A to G) (Supplementary Table 5).

Supplementary Fig. 7

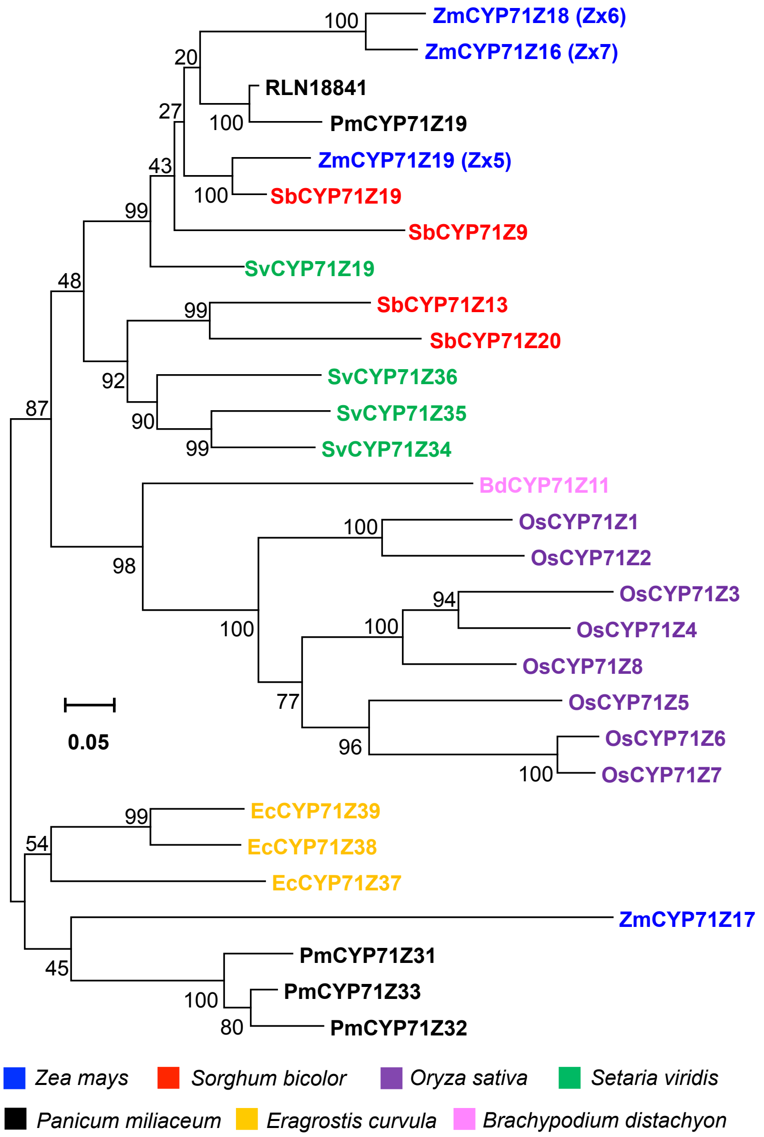


**Supplementary Fig. 7 | Phylogenetic tree of CYP71Z subfamily**. Cytochrome P450s from grass species *Zea mays (Zm), Sorghum bicolor (Sb), Oryza sativa (Os), Setaria viridis (Sv), Panicum miliaceum (Pm), Eragrostis curvula (Ec),* and *Brachypodium distachyon (Bd)* with amino acid sequence identity >55% were included (ZmCYP71Z19 as a seed query). Tree reconstruction was performed with the maximum likelihood algorithm using MEGA 7 program (www.megasoftware.net). Bootstrap values calculated from 1000 iterations are indicated at the nodes. The protein accession numbers are listed in the Supplementary Table 3.

Supplementary Fig. 8

```
CYP71Z19   MEQKVLVAVGVAVLLVVVLSKLKSVLVTKPKLNLPPGPWTLPLIGSHHLVTSPSIYRAMRDLAQKYGPLMMIRLGEVPT 80
CYP71Z18   MEDKVLIAVG-TVAVVAVLSKLKS-AVTKPKLNLPPGPWTLPLIGSIHHVSNPLPYRAMRELAHKHGPLMMLWLGEVPT 78
CYP71Z16   MEDKVLLAVA-MVALIAVLSKLKSLLETKPKLNLPPGPWTIPLIGSIHHLVSSPLPYRAMRELAHKHGPLMMLWLGEVPT 79

CYP71Z19   LVVSSPEAAQAITKTHDAFADRHMNTTIGVLTFNGTDLVFGPYGERWRQLRKICVLELFSVARVQSFQRIREEEVARFM 160
CYP71Z18   LVVSSPEAAQAITKTHDVSFADRHINSTVDILTFNGMDMVFGSYGEQWRQLRKLSVLELLSARVQSFQRIREEEVARFM 158
CYP71Z16   LVVSSPEAAQAITKTHDVTFADRHMNSTVDILTFNGDIVFGTYGEQWRQLRKLSVLELLSVARVQSFQRIREEEVARFM 159

CYP71Z19   QSLAASAG---TVNLSKMISRFINDTFVRECIGSRCKYQDEYLDAFDTAVRQTSVLTVADLFPSSRIMQAVGTAPRNALK 237
CYP71Z18   RSLAASASAGATVDLSKMISSFINDTFVRESIGSRCKYQDELAALDTAIRVAAELSVGNIFPSSRVLQSLSTARRKAIA 238
CYP71Z16   RNLAASAGAGATVDLSKMISSFINDTFVRESIGSRCKQDEYLDAIHTGIRVAAELSVANLFPSSRLLQSLSTARRKVA 239

CYP71Z19   CRNRITRILEQIIREKVEAMGRGEKTAHEGLIGVLLRLQKEANLPTLLTNDTIVALMFDLFGAGSDTSSTTNWCITELI 317
CYP71Z18   SRDEMARILGQIIRETKSMDQGDKTSNESMISVLLRLQKAGLPIELTDNVVMALMFDLFGAGSDTSSTTLTWCMTEVL 318
CYP71Z16   ARDEMARILGQIIRETKEAMVGDKASNESMISVLLRLQKEAGLPIELTDIVMALMFDLFGAGSDTSSTTLTWCMTMEI 319

CYP71Z19   RHPAAMAKAQAEVREAFKGKARIISEDDLAGAGLSYLKLVIKEALRMHCPLPLLLPRLCRETCQVMGYDIPKGTAVFINV 397
CYP71Z18   RYPATMAKAQAEVREAFKGK-TTITEDDLSTANLRYLKLVVKEALRLHCPVPLLLPRKCRACQVMGYDIPKGTCVFVNV 397
CYP71Z16   RYPATMAKAQAEVREAFKGK-TTITEDDLSRANLSYLKLVVKEALRLHCPVPLLPRKCRETCQIMGYDIPKDTCVLVNV 398

CYP71Z19   WAVCRDAKYWEDPEEFRPERFEDTNLEYNYKGTNYEFLPFGSGRRMCPGANLGLGNIELALASLLYHYDWKLPDGVKPQD 477
CYP71Z18   WAICRDPRYWEDAEEFKPERFENSNLDY-KGTYYEYLPFGSGRRMCPGANLGVANELALASLLYHFDWKLPSGQEPKD 475
CYP71Z16   WAICRDSRYWEDADEFKPERFENSSLDY--KGTSHEYLPFGSGRRMCPGGNLGVANMELALASLLYHFDWKLPSGQEPKD 476

CYP71Z19   VQVWEGPGLIAKKKTGLLLRPVTCIAFACSSG 509
CYP71Z18   VDVWEAAGLVAKKNIGLVLHPVSHIAPVNA-- 505
CYP71Z16   VDVWEAAGLVGRKNAGLVLHPVSRFAPVNA-- 506
```

**Supplementary Fig. 8 | Encoded amino acid sequence comparison of three B73 CYP71Z subfamily P450s present in zealexin gene cluster II.** The alignment of predicted amino acid sequences encoded by the B73 *ZmCYP71Z19* (*Zx5, Zm00001d014121*), *ZmCYP71Z18* (Zx6, *Zm00001d014134*), and *ZmCYP71Z16* (*Zx7, Zm00001d014136*) with the program MEGA7 (www.megasoftware.net) and the MUSCLE (codon) algorithm. The visualization was performed with the program BIOEDIT (http://www.mbio.ncsu.edu/BioEdit). B73 Zx5 shares 71.5-72.1% protein sequence identity to both Zx6 and Zx7, while Zx6 and Zx7 have 89.3% protein sequence identity to each other. The protein accession numbers are listed in the Supplementary Table 3.

Supplementary Fig. 9

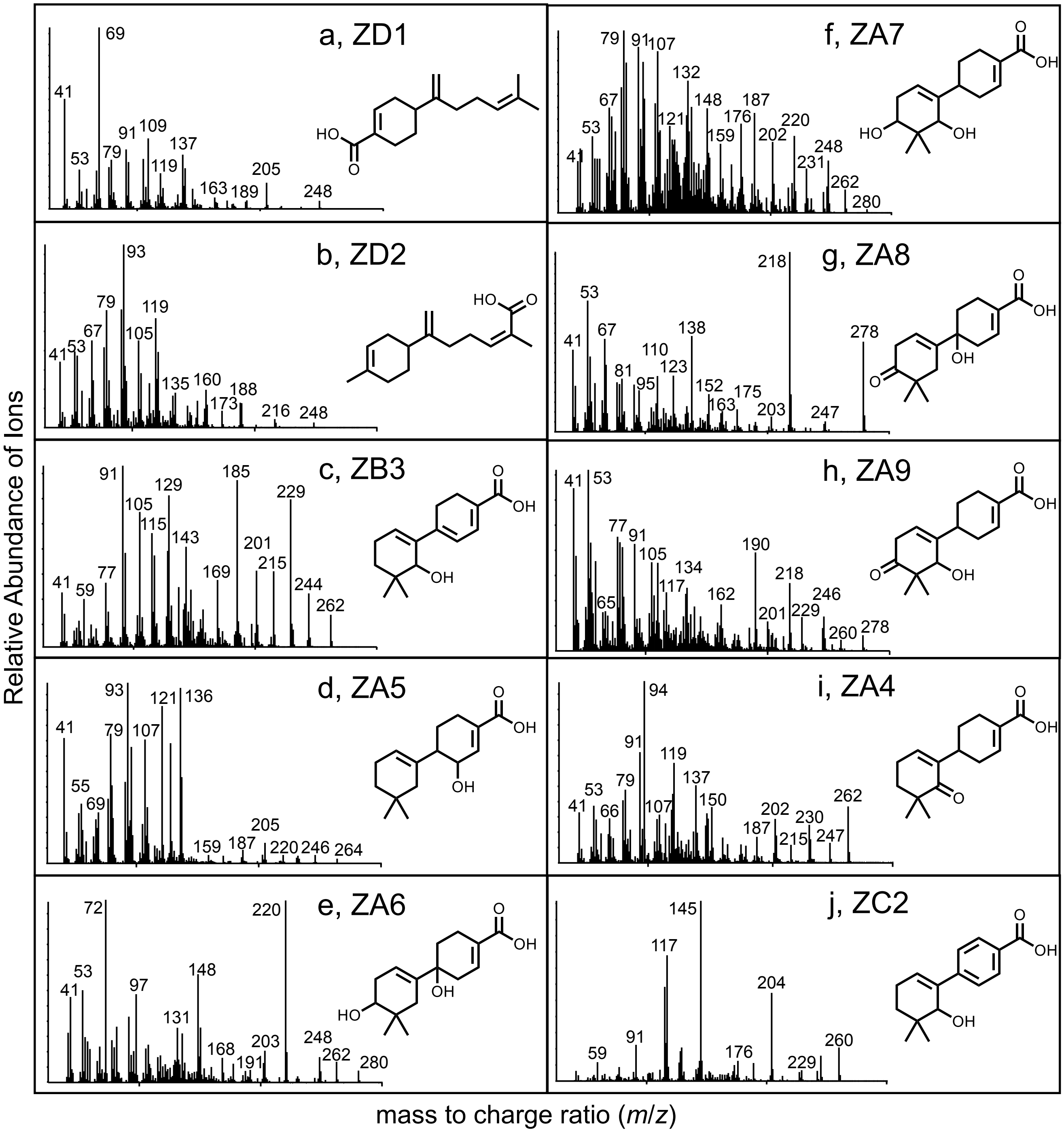


**Supplementary Fig. 9 | Reference electron ionization (EI) spectra of zealexins identified in the current study following methyl ester derivatization.** Maize metabolites purified, identified by NMR, derivatized with trimethylsilyldiazomethane and analyzed by electron ionization (70 eV). Two β-bisabolene derivatives include: **a**, ZD1 [4-(6-methylhepta-1,5-dien-2-yl)cyclohex-1-ene-1-carboxylic acid] and **b**, ZD2 [(E)-2-methyl-6-(4-methylcyclohex-3-en-1-yl)hepta-2,6-dienoic acid]. Six β-macrocarpene derivatives include **c**, ZB3 [6'-hydroxy-5',5'-dimethyl-[1,1'-bi(cyclohexane)]-1,1',3-triene-4-carboxylic acid]; **d**, ZA5 [2-hydroxy-5',5'-dimethyl-[1,1'-bi(cyclohexane)]-1',3-diene-4-carboxylic acid]; **e**, ZA6 [1,4'-dihydroxy-5',5'-dimethyl-[1,1'-bi(cyclohexane)]-1',3-diene-4-carboxylic acid]; **f**, ZA7 [4',6'-dihydroxy-5',5'-dimethyl-[1,1'-bi(cyclohexane)]-1',3-diene-4-carboxylic acid]; **g**, ZA8 [1-hydroxy-5',5'-dimethyl-4'-oxo-[1,1'-bi(cyclohexane)]-1',3-diene-4-carboxylic acid]; **h**, ZA9 [6'-hydroxy-5',5'-dimethyl-4'-oxo-[1,1'-bi(cyclohexane)]-1',3-diene-4-carboxylic acid]. **i**, ZA4 and **j**, ZC2 follow from Christensen et al. (2018)[1] and Suzuki et al (2007)[2], respectively. Reference spectra for β-macrocarpene, ZA1, ZA2, ZA3 and ZB1 follow from the original description of ZmTPS6/11[3] and zealexins[4]. Combined spectra were used to identify enzyme products of *Agrobacterium*-mediated transient *N. benthamiana* co-expression assays.

References: [1]Christensen *et al*. (2018) *Planta*. 247(4):863-873; [2]Suzuki R, Iijima M, Okada Y, Okuyama T (2007) *Chem Pharm Bull* (Tokyo) 55: 153–155; [3]Kollner *et al*. (2008). *J Biol Chem* 283: 20779–20788; [4]Huffaker *et al*. (2011) *Plant Physiol*. 156(4):2082-2097

### Supplementary Fig. 10

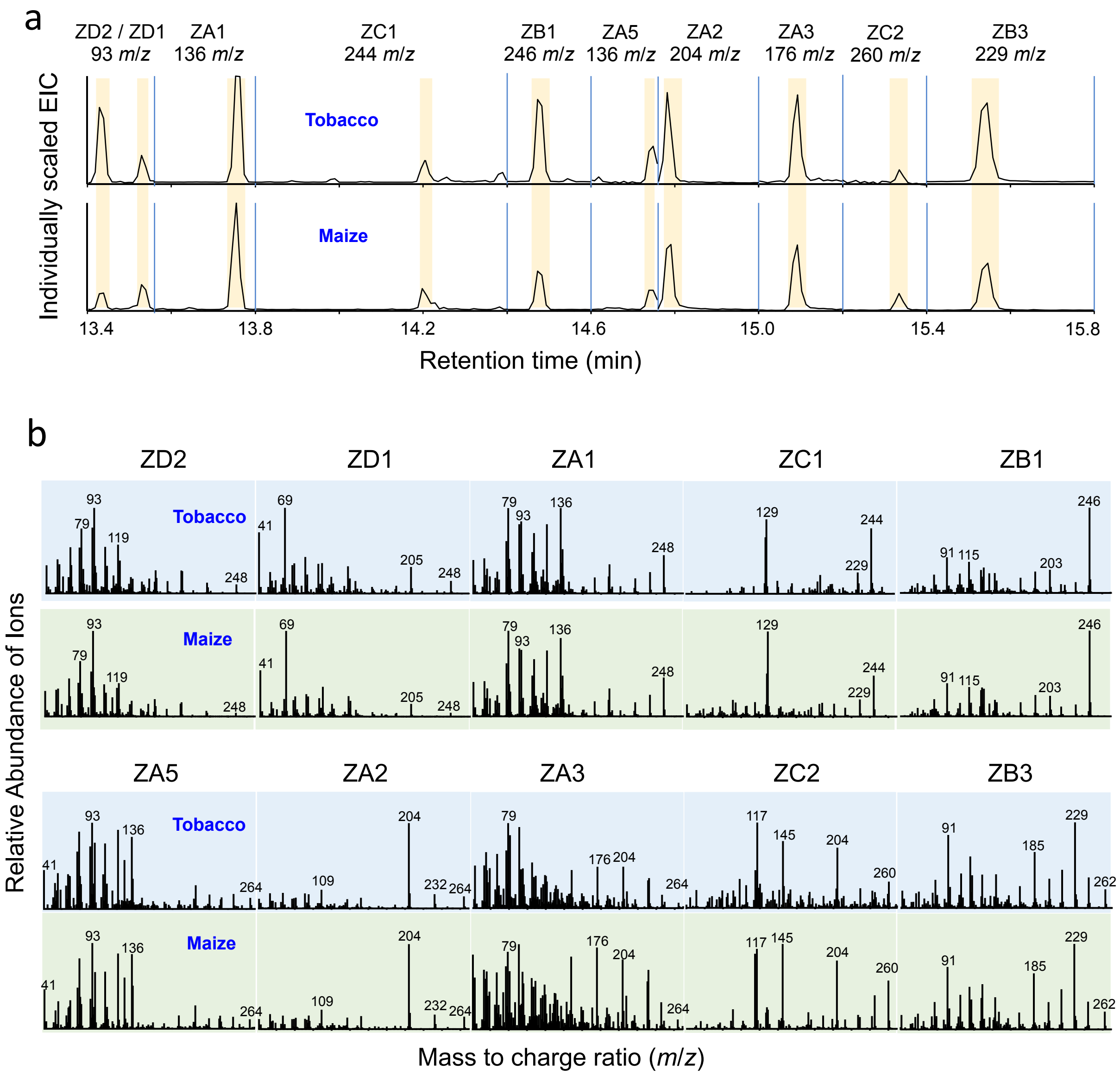


**Supplementary Fig. 10 | Representative GC-MS retention times, EI diagnostic (*m/z*) ions, and full EI spectra of zealexins present in plant extracts from transient expression assays in *N. benthamiana* and fungal-elicited maize stems. a**, Representative GC-MS extracted ion chromatograms (EIC) showing diagnostic EI (*m/z*) ions and relative retention times of zealexin methyl ester derivatives detected in extracts *Agrobacterium*-mediated transient *N. benthamiana* (tobacco) co-expression assays and fungal-elicited maize stems. **b**, EI mass spectra of zealexins from (**a**) as methyl esters. Mass spectra highlighted with blue are from transient expression assays in *N. benthamiana* (tobacco) and those with green are from elicited maize stems. Reference metabolite EIC traces and EI spectra were derived from the following enzyme pairs expressed in *N. benthamiana*: ZD1 and ZD2 (SaMonoTPS + Zx6); ZA1(Zx3 + Zx6); ZB1 and ZC1 (Zx3 + Zx6 + Zx8); ZA5, ZA2, ZA3, ZC2, and ZB3 (Zx3 + Zx6 + Zx8 + Zx10).

Supplementary Fig. 11

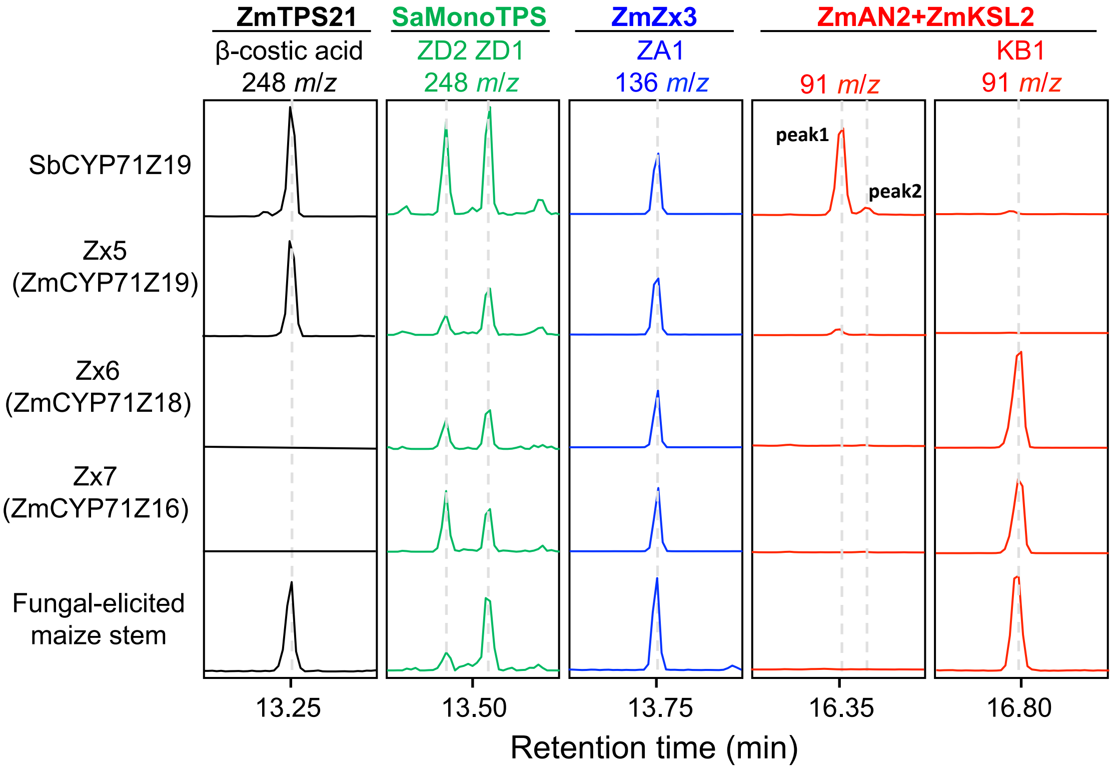


**Supplementary Fig. 11 | The sorghum homolog of ZmCYP71Z19 (Zx5) oxidizes diverse terpene olefins.** GC-MS extracted ion chromatograms (EIC) of preparations derived from *Agrobacterium*-mediated transient *N. benthamiana* co-expression assays of Mo17 TPS21, SaMonoTPS, Zx3 or ZmAN2 + ZmKSL2 with SbCYP71Z19, ZmCYP71Z19, ZmCYP71Z16 or ZmCYP71Z18 showing production of β-costic acid (*m*/*z* 248), ZD2 (*m*/*z* 248), ZD1 (*m*/*z* 248), ZA1 (*m*/*z* 136), *ent*-kaur-15-17-ol (*m*/*z* 91, peak 1)[1], *ent*-kaur-15-17-al (*m*/*z* 91, peak 2)[2], and KB1 (*m*/*z* 91) as methyl ester derivatives. Four independent experiments were preformed and showed similar results. References: [1]Zhang *et al.* Chemical constituents of *Aristolochia constricta*: antispasmodic effects of its constituents in guinea-pig ileum and isolation of a diterpeno-lignan hybrid. *J Nat Prod*. 2008, 71:1167-72. [2]Su WC, Fang JM, Cheng YS. Abietanes and kauranes from leaves of *Cryptomeria japonica*. *Phytochemistry*. 1994, 35:1279–1284.

.

Supplementary Fig. 12

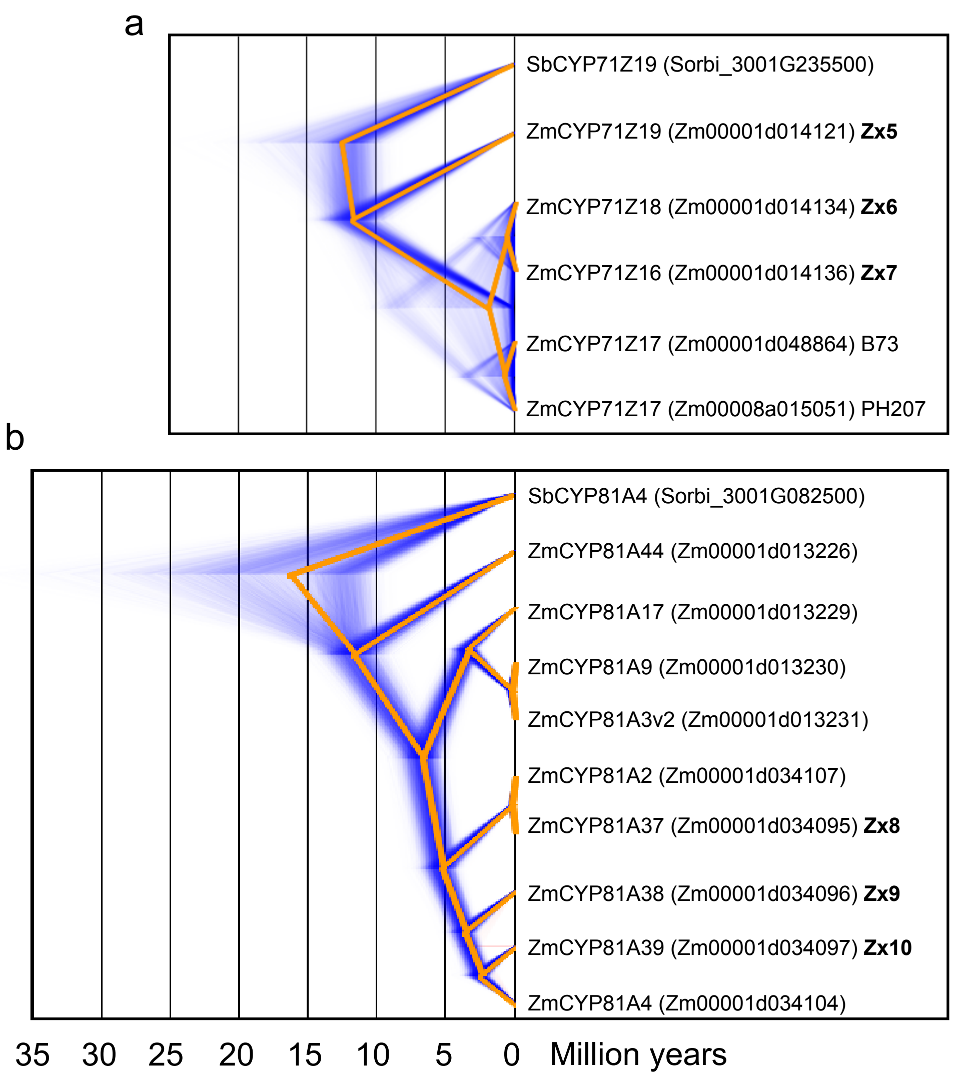


**Supplementary Fig. 12 | DensiTree visualizations of the relationships among the maize *CYP71Z* gene subfamily (A) and the *CYP81A* gene subfamily (B) in the B73 RefGen v4 reference genome.** The *ZmCYP71Z* gene subfamily appears to have arisen from a recent duplication of the *ZmCYP71Z19* (*Zx5*: *Zm00001d014121*) gene. The *ZmCYP81A* gene subfamily appears to have arisen from a more ancient duplication of the *ZmCYP81A4* (*Zm00001d034104*) gene. Orange lines show the consensus tree and blue shading shows uncertainty in the age estimates of nodes in the tree.

Supplementary Fig. 13

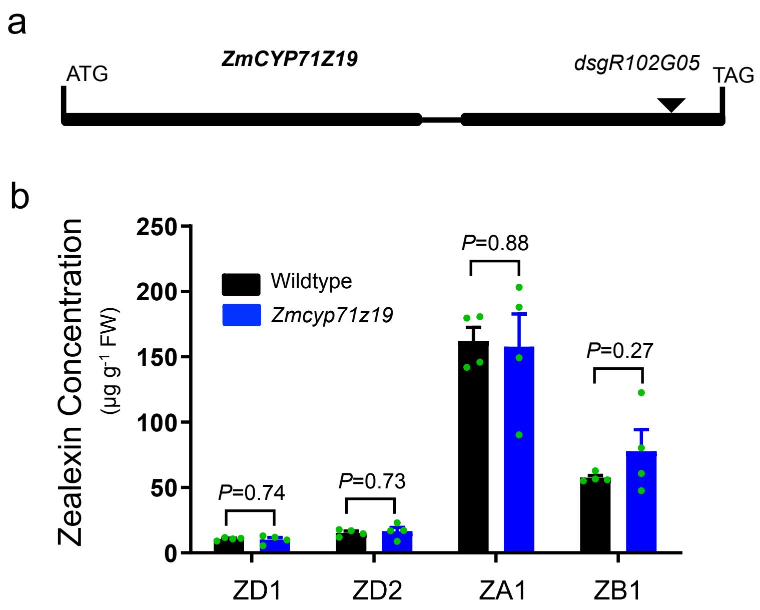


**Supplementary Fig. 13 | Characterization of the W22 *zx5* (*Zmcyp71z19*) *dsg*-transposon mutant supports an endogenous role of Zx6 and Zx7 in zealexin pathway redundancy**. **a**, The *Dsg* is inserted in exon 2 of the W22 *Zmcyp71z19* gene. **b**, Average quantity of ZD1, ZD2, ZA1 and ZB1 from W22 wild type siblings and *Zmcyp71z19* plants treated with heat-killed *F. venenatum* hyphae for 3 days. Error bars indicate mean ± s.e.m. (*n* = 4 biologically independent replicates). *P* values represent Student's *t* test, two-tailed distribution, equal variance.

Supplementary Fig. 14

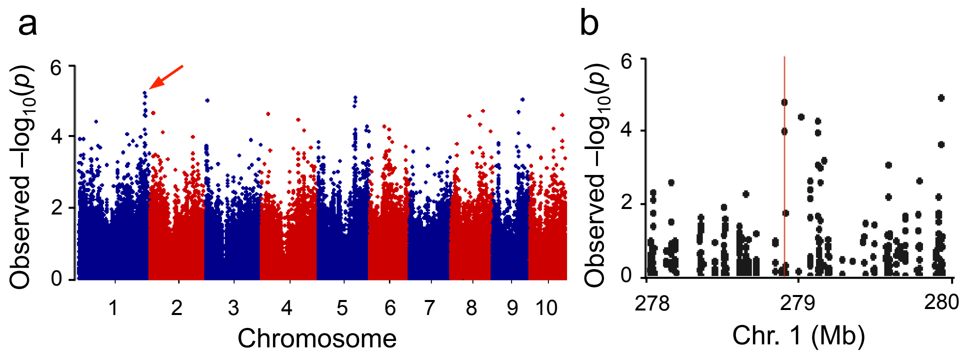


**Supplementary Fig. 14 | An association analysis using the Goodman diversity panel provides additional support for the third zealexin gene cluster. a**, Manhattan plot of the Goodman diversity panel association analysis using ratio of ZB1 to ZA1 as a mapping trait. 35 days after planting 258 Goodman diversity lines were wounded, stem elicited with heat-killed *F. venenatum* hyphae, harvested after 3 days and analyzed by LC-MS. Negative log10-transformed *P* values from the compressed mixed linear model are plotted on the y axis. 246,477 SNP markers identified (red line arrow) the largest log10-transformed *P* located on Chr. 1 (B73 RefGen_v2). **b,** Local Manhattan plot surrounding the peak on Chr. 1.

Supplementary Fig. 15

```
                                                                    Proline-rich region
B73 CYP81A37_Zx8      METANVVILSFFFIFAIHRLFNRLHSRSTKMKPLPPGPLAIPVLGHLYLLLEKNMHHLFT  60
Mo17 CYP81A37_Zx8     METAYVVILSFFFTFAIHRLLNRRHSRNMNTKQLPPGPLAIPVLGHLH-LLEKKIHDSFT  59
B73 CYP81A38_Zx9      METANVVILS-FFIFAIHRLFNRLHSRSTKMKPLPPGPLAIPVLGHLYLLLEKNIHHMFT  59
Mo17 CYP81A38_Zx9     METANVIILSFFFIFAIHRLLNRLHNRSTKMKQLPPGPLAIPVLGHLYLLLEKNIHHLFT  60
B73 CYP81A39_Zx10     METAYVVVLSFFFIVAIHRLLNRGESKNMSKQPLPPGPRAIPVLGHLH-LLEKPYHLAFM  59
Mo17 CYP81A39_Zx10    METAFVVVLSFLFIVAIHRLLNRGESKNMSKQPLPPGPRAIPVLGHLH-LLEKPYHLAFM  59

B73 CYP81A37_Zx8      RLAARYGPVLYLRLGSRNAVVVSSVDCARECFTEHDVTFANRPTFPTLHLMTYGGTTVGT  120
Mo17 CYP81A37_Zx8     RLAARYGPVLYVRLGSRDAVVVSSVDCARECFTEHDVTFANRPSFPTLHLMTYGGTTVGT  119
B73 CYP81A38_Zx9      RLAARYGPVLYLRLGSRNAVIVSSVDCARECFTEHDVTFANRPTFPTLHLMTYGGTTVGT  119
Mo17 CYP81A38_Zx9     RLAARYGPVLYLRLGSRNAVVVSSVDCARECFTEHDVTFANRPTFPTLHLMTYGGTTVGT  120
B73 CYP81A39_Zx10     RLAARYGPVFSLQLGSRAAVVVSTADCARECLAEHDATFANRPTYPTLHLMTYGGATIGH  119
Mo17 CYP81A39_Zx10    RLAARYGPVFSLQLGSRAAVVVSTADCARECLAEHDATFANRPTYPTLHLMTYGGATIGH  119

B73 CYP81A37_Zx8      CAYGPYWRHIRRVITVHLLSALRVR-SMVPAIEAEVRAMARSMYRAAAAAPCGTGAAKVE  179
Mo17 CYP81A37_Zx8     CAYGPYWRHIRRVITVHLLSAQRVR-AMVPAIEAEVRAMARSMYRAAVTAPGGAGAAKVE  178
B73 CYP81A38_Zx9      CAYGPYWRHIRRVITVHLLSALRVR-SMVPAIEAEVRAMARSMYRAAAAAPCGTGAAKVE  178
Mo17 CYP81A38_Zx9     CAYGPYWRHIRRVITVHLLSALRVR-SMVPAIEAEVRAMARSMYRAAAAAPCGTGAAKVE  179
B73 CYP81A39_Zx10     SAYGAHWRHVRRVVSLHLLSPQRVLSSMVPTITAEVRAMARRMYRAAAG-----GAARIE  174
Mo17 CYP81A39_Zx10    SAYGAHWRHVRRVVSLHLLSPQRVLSSMVPTITAEVRAMARRMYRAAAG-----GAARIE  174

B73 CYP81A37_Zx8      LRGRLFEVALSALMETVAQSKTSRSSMGGAADTGMSPEAQEFKESMDVMIPLLGTANMWD  239
Mo17 CYP81A37_Zx8     LRGMLFELALSALMETVAKTKTSRT--VDAADTGLSPEAREFKESMDVMVPLLGTANMWD  236
B73 CYP81A38_Zx9      LRGRLFEVALSALMETVAQSKTSRSSMGGAADTGMSPEAQEFKESMDVMVPLLGTANMWD  238
Mo17 CYP81A38_Zx9     LRGRLFEVALSALMETVAQSKTSRSSMGGAADTGMSPEAQEFKESMDVMVPLLGTANMWD  239
B73 CYP81A39_Zx10     LKGRLLELSLSTLMETIAQTKTYRP--ADDADTDMSPETQEFKQLLDAIISLLGLTNVLE  232
Mo17 CYP81A39_Zx10    LKGRLLELSLSTLMETIAQTKTYRP--ADDADTDMSPETQEFKQLLDAIISLLGLTNVLE  232

B73 CYP81A37_Zx8      FLPVLQRFDVFGVKNKMAAAVSGRDAFFRRLIDEERRRLDDGVESENKSMMAVLLTLQKS  299
Mo17 CYP81A37_Zx8     FLPVLQRFDVFGVKNKIAAAVSSRDAFFRRLIDAERRRLDGGVKSEDKSMMAVLLTLQKS  296
B73 CYP81A38_Zx9      FLPVLQRFDVFGVKNKMAAAVSGRDAFFRRLIDEERRRLDDGVESENKSMMAVLLTLQKS  298
Mo17 CYP81A38_Zx9     FLPVLQRFDVFGVKNKMAAAVSGRDAFFRRLIDEERRRLDDGVQSENKSMMAVLLTLQKS  299
B73 CYP81A39_Zx10     FLPVLQTFDVLGLKKKIAAAAGRRDAFFQRLIDAERRRLDDGVEGQKKSMLAVLLALQKS  292
Mo17 CYP81A39_Zx10    FLPVLQTFDVLGLKKKIAAAAGRRDAFFQRLIDAERRRLDDGVEGQKKSMLAVLLALQKS  292

B73 CYP81A37_Zx8      EPENYSDDMIMSLCFSMFSAGTETTATTAEWAMSLLLNNPEVLKKAQGEIDAHVGNSRLL  359
Mo17 CYP81A37_Zx8     EPENYNDDMIMSLCFSMFSAGTETTATTAEWAMSLLLNHPEVLKKAQGQIDAYVGNSRLL  356
B73 CYP81A38_Zx9      EPENYSDDMIMSLCFSMFSAGTETTATTAEWAMSLLLNNPEVLKKAQGEIDAHVGNSRLL  358
Mo17 CYP81A38_Zx9     EPENYSDDMIMSLCFSMFSAGTETTATTAEWAMSLLLNNPEVLKKAQGEIDAHVGNSRLL  359
B73 CYP81A39_Zx10     EPENYTDEKIMALCFSMFIAGTDSTATAMEWAMSLLLNHPEVLKKAREEIDAHVGSSRLL  352
Mo17 CYP81A39_Zx10    EPENYTDEKIMALCFSMFIAGTDSTATAMEWAMSLLLNHPEVLKKAREEIDAHVGSSRLL  352
                                    O2-binding region (A/G)GX(D/E)T(T/S)

B73 CYP81A37_Zx8      GADDMPHLPYLQCVLTETLRLYPVFPMLIAHESTADCKVGGHHVPSGTMLLTNAYAIHRD  419
Mo17 CYP81A37_Zx8     CADDMPHLPYLQCILTETLRLYPIIPMLIAHESTADCKVGGHHVPSGTMLLVNAYAIHRD  416
B73 CYP81A38_Zx9      GADDMPHLPYLQCVLTETLRLYPVFPMLIAHESTADCKVGGHHVPSGTMLLTNAYAIHRD  418
Mo17 CYP81A38_Zx9     GADDMPHLPYLQCVLTETLRLYPVFPMLIAHESTADCKVGGHHVPSGTMPLTNVYAIHRD  419
B73 CYP81A39_Zx10     GADDVPSLGYLHCVLNETLRLYPVGPTLIPHESTADCTVGGYRVPSGTMLLVNVYAIHRD  412
Mo17 CYP81A39_Zx10    GADDVPSLGYLHCVLNETLRLYPVGPTLIPHESTADCTVGGYRVPSGTMLLVNVYAIHRD  412

B73 CYP81A37_Zx8      PAAWTEPDAFRPERFEDG--SAEGKLLIPFGMGRRKCPGETMALRTLGLVLGTLIQCFDW  477
Mo17 CYP81A37_Zx8     PAAWMDSTVFRPERFEDGSASAEGRLLIPFGMGRRKCPGETMALRTLGLVLGTLIQCFDW  476
B73 CYP81A38_Zx9      PAAWTEPDAFRPERFEDG--SAEGKLLIPFGMGRRKCPGETMALRTLGLVLGTLIQCFDW  476
Mo17 CYP81A38_Zx9     PAAWTEPDAFRPERFEDG--SAEGKLLIPFGMGRRKCPGETMALRTLGLVLGTLIQCFDW  477
B73 CYP81A39_Zx10     PATWPDPDVFRPERFEDGGGSAEGRLLIPFGMGRRKCPGETMALQIMGLALGTMIQCFDW  472
Mo17 CYP81A39_Zx10    PATWPDPDVFRPERFEDGGGSAEGRLLIPFGMGRRKCPGETMALQTMGLALGTMIQCFDW  472
                                                  Heme binding PFGXGRRXCXG

B73 CYP81A37_Zx8      ATVGG--VPKVDMTEASGLTLPRAVPLEAMCKPRQAMLDVLQNL  519
Mo17 CYP81A37_Zx8     ATVGG--VPKVDMTEASGLTLPRAVPLEAMCKPRQAMLDVLQNL  518
B73 CYP81A38_Zx9      ATVGG--VPKVDMTEASGLTLPRAVPLEAMCKPRQAMLDVLQKL  518
Mo17 CYP81A38_Zx9     ATVGG--VPKVDMTEASGLTLPRAVPLEAMCKPRQAMLDVLQKL  519
B73 CYP81A39_Zx10     GAVGGGGAPKVDMTQGGGLTLPRAVPLEAMCKPRQVMLDVLQKL  516
Mo17 CYP81A39_Zx10    GAVGGGEAPKVDMTQGGGLTLPRAVPLEAMCKPRQVMLDVLEKL  516
```

**Supplementary Fig. 15 | Encoded amino acid sequence comparison of zealexin gene cluster III CYP81A subfamily P450s in B73 and Mo17 inbreds.** The alignment was constructed based on predicted amino acid sequences of ZmCYP81A37, **Zx8** (B73, Zm00001d034095; Mo17, deduced from genomic sequence no assigned ID), ZmCYP81A38, **Zx9** (B73, Zm00001d034096; Mo17, Zm00014a026557) and ZmCYP81A39, **Zx10** (B73, Zm00001d034097; Mo17, Zm00014a026556) with the program MEGA7 (www.megasoftware.net) and the MUSCLE (codon) algorithm. The visualization was done with the program BIOEDIT (http://www.mbio.ncsu.edu/BioEdit). B73 Zx8 shares 99% and 72% amino acid identity with B73 Zx9 and B73 Zx10, respectively.

Supplementary Fig. 16

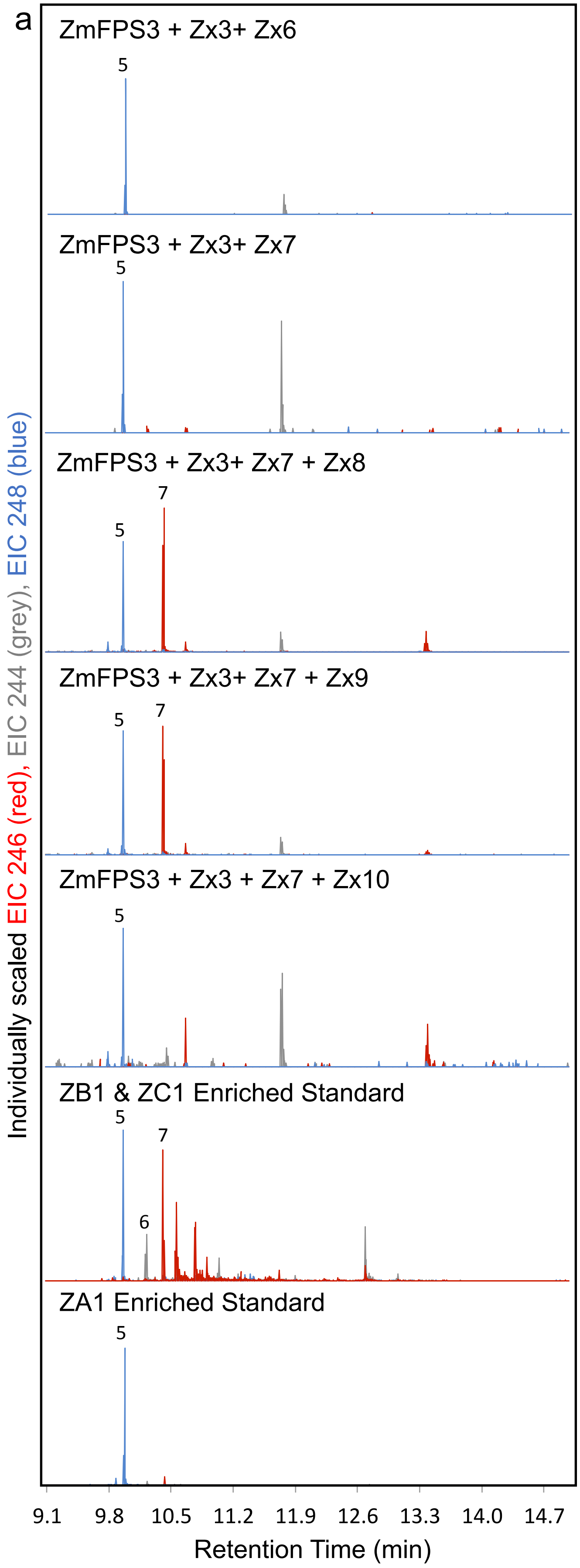


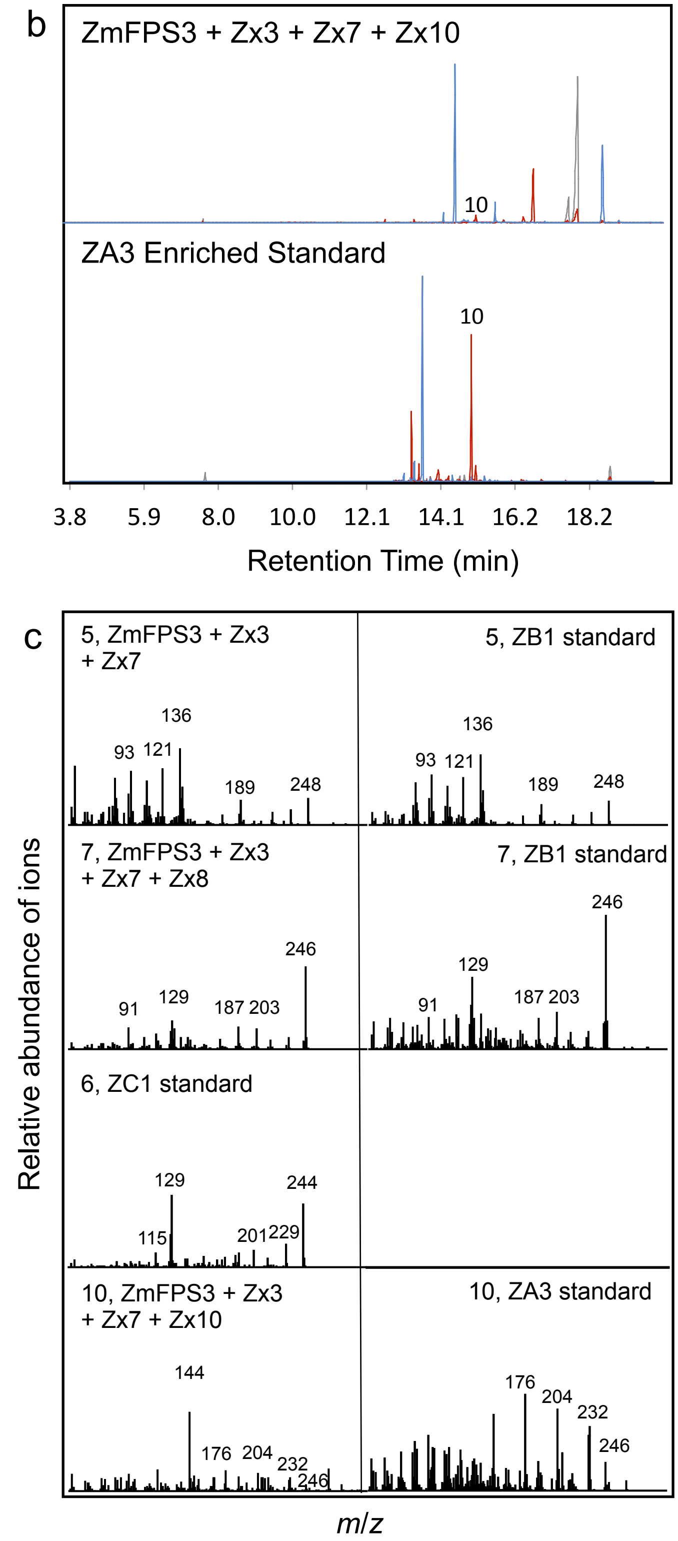


**Supplementary Fig. 16 | Heterologous co-expression of early zealexin enzymes with Zx8, Zx9 and Zx10 in *E. coli* supports roles in pathway expansion. a**, GC-MS extracted ion chromatograms (EIC) of ZmFPS3 and ZmCPR2 co-expressed with Zx3 + Zx6, Zx3 + Zx7, Zx3 + Zx7 + Zx8, Zx3 + Zx7 + Zx9 and Zx3 + Zx7 + Zx10. Following unified numbering from Supplementary Fig. 1, compound 5 is ZA1, compound 7 is ZB1 and compound 6 is ZC1. Enriched zealexin standards are derived from root extracts and all were runs analyzed on a HP5-MS column. **b**, GC-MS extracted ion chromatograms (EIC) of ZmFPS3 and ZmCPR2 co-expressed Zx3, Zx7, and ZX10 run on a separate GC column (DB-35) and MS instrument. The ZA3 enriched standard was a Mo17 root extract. Compound 10 is ZA3. **c**, All compounds in (**a**) and (**b**) were analyzed as methyl ester derivatives.

Supplementary Fig. 17

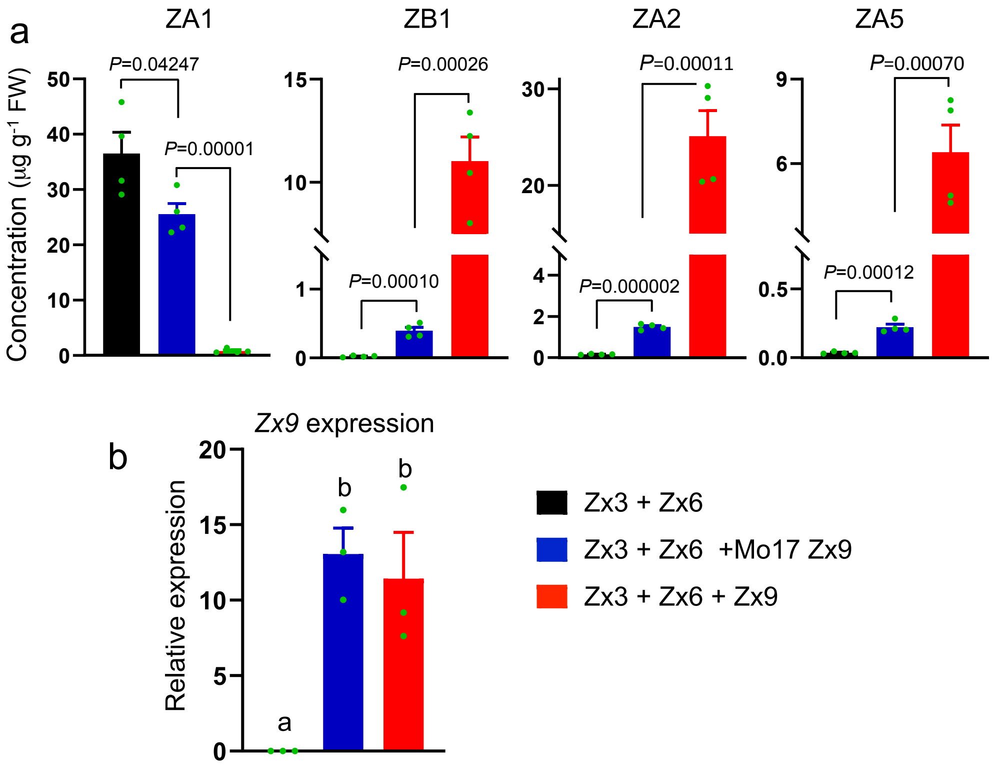


**Supplementary Fig. 17 | Mo17 Zx9 (ZmCYP81A38) retains residual catalytic activity towards ZA1. a,**Transient expression of Zx3 + Zx6 with Zx9, Mo17 Zx9, or empty vector in *Nicotiana benthamiana* demonstrates that Mo17 Zx9 has residual enzymatic activity in converting ZA1 to ZB1, ZA2 and ZA5. Zealexins extracted from 5-day *Agrobacterium*-inoculated leaves were measured with GC-MS. Error bars indicate mean ± s.e.m. (*n* = 4 biologically independent replicates). Two-tailed *P*-values from Student's *t* test (unpaired) are shown above bars to indicate significant differences. **b,** qrtPCR showing *Zx9* expression from transient assays of Zx3+Zx6 with Zx9, Mo17 Zx9, or empty vector in *Nicotiana benthamiana.* Error bars indicate mean ± s.e.m. (*n* = 3 biologically independent replicates). Within plots, different letters (a–b) represent significant differences (ANOVA $P < 0.05$; Tukey's test corrections for multiple comparisons, $P <0.05$).

### Supplementary Fig. 18

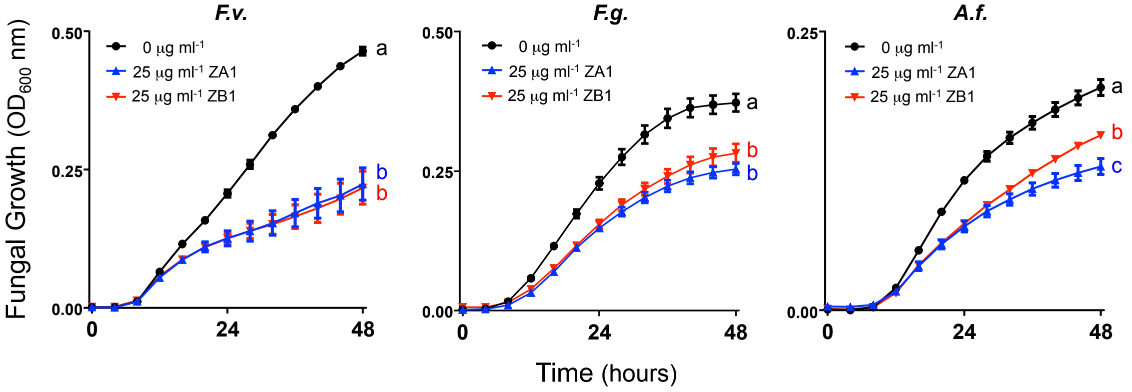


**Supplementary Fig. 18 | Zealexin gene cluster III product ZB1 displays antibiotic activity against maize fungal pathogens.** Fungal growth estimates measured at 600 nm (optical density) of *Fusarium verticillioides* (*F.v.*), *Fusarium graminearum* (*F.g.*) and *Aspergillus flavus* (*A.f.*) in liquid medium in the presence of a dimethylsulfoxide (DMSO) solvent control (0 µg $ml^{-1}$; black circles), ZA1 (25 µg $ml^{-1}$; blue arrows), or ZB1 (25 µg $ml^{-1}$; red arrows). Error bars indicate mean ± s.e.m. ($n$ = 6 biologically independent replicates) and different letters (a–c) represent significant differences (one-way ANOVA followed by Tukey's test corrections for multiple comparisons, $P < 0.05$).

Supplementary Fig. 19

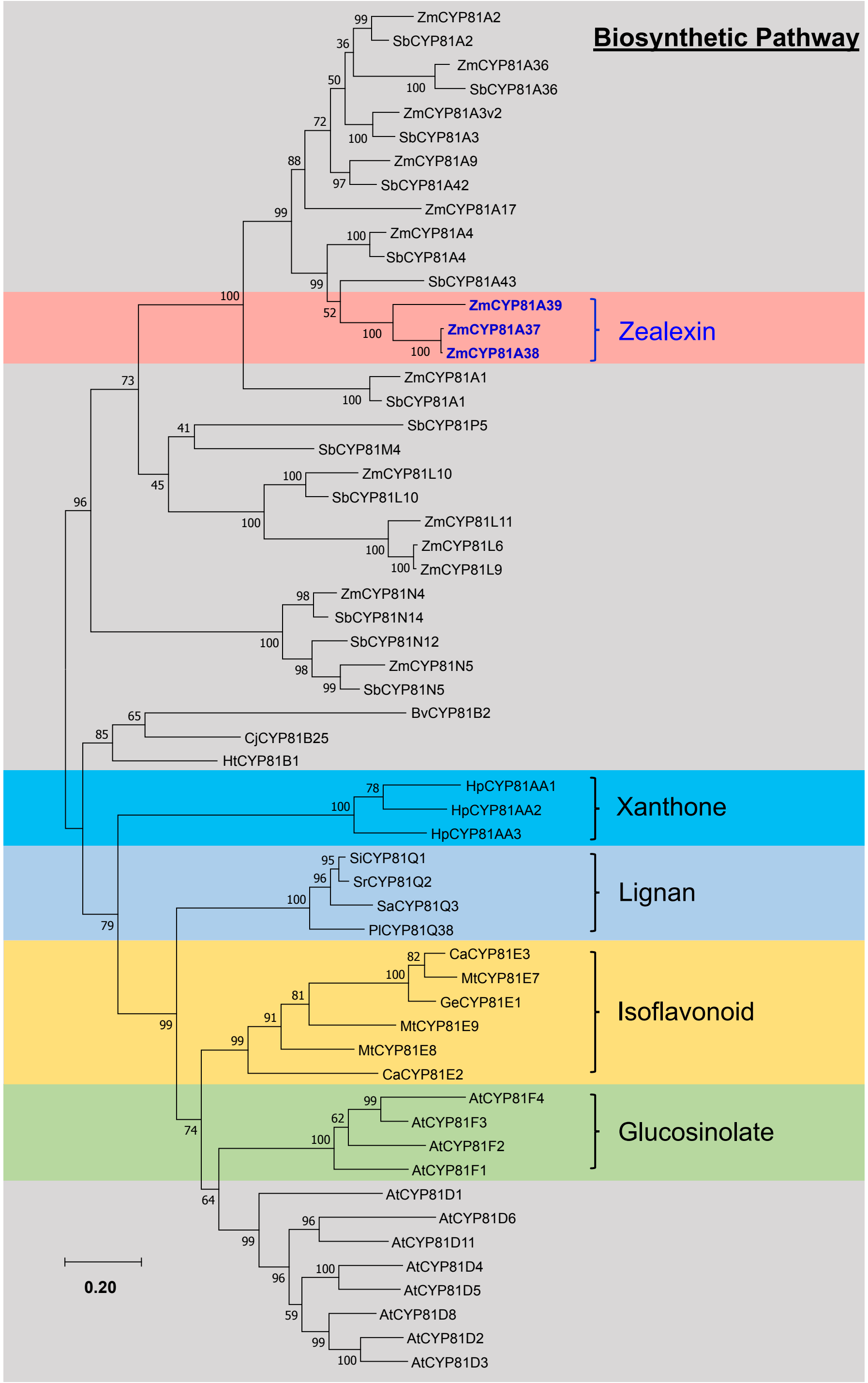


**Supplementary Fig. 19 | Zealexin gene cluster III expands the known roles of CYP81 enzymes to include sesquiterpenoid defenses.** Phylogenetic analysis of *Zea mays* (Zm) CYP81 subfamily of P450s together with representative members of the CYP81 subfamily based on amino acid sequences from *Arabidopsis thaliana* (At), *Beta vulgaris* (Bv), *Cicer arietinum* (Ca), *Coptis japonica* (Cj), *Glycyrrhiza echinate* (Ge), *Helianthus tuberosus* (Ht), *Hypericum perforatum* (Hp), *Medicago truncatula* (Mt), *Phryma leptostachya* (Pl), *Sesamum alatum* (Sa), *Sesamum indicum* (Si), *Sesamum radiatum* (Sr), and *Sorghum bicolor* (Sb). Tree reconstruction was performed with the maximum likelihood algorithm using MEGA 7 program. Bootstrap values calculated from 1000 iterations are indicated at the nodes. The protein accession numbers and references to specific roles in glucosinolate, isoflavonoid, lignin, and xanthone are listed in the Supplementary Table 3.

Supplementary Fig. 20

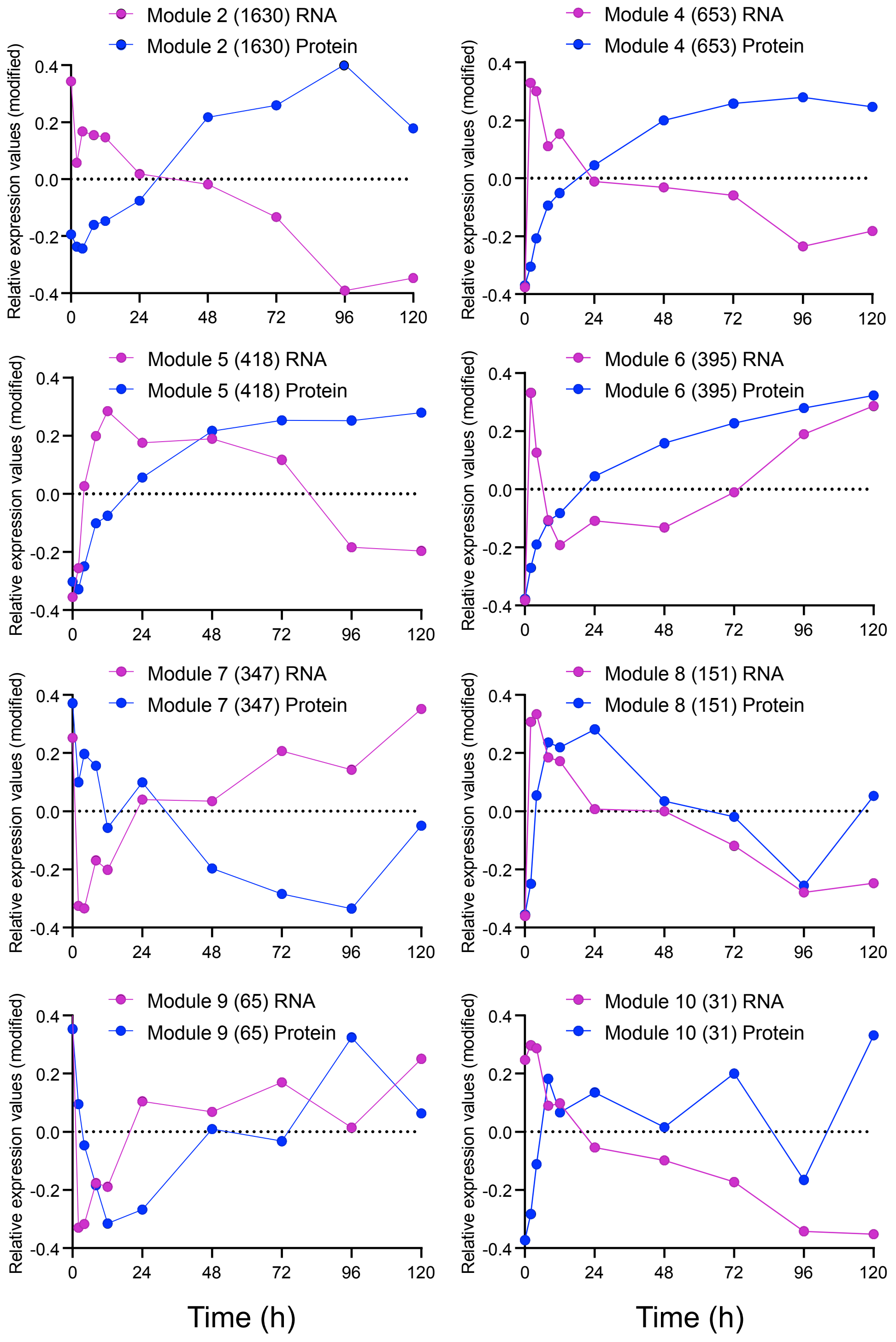


**Supplementary Fig. 20 | Weighted Gene Co-Expression Network Analysis (WGCNA) of W22 transcriptome and proteome changes following *Fusarium* elicitation yields 10 module eigengenes dominated by co-suppression, co-activation and dysregulated patterns.** A ten point 120 h *F. venenatum* elicitation time course in W22 stems utilized 38 day old plants. Graphs represent eight ordered RNA and protein module eigengenes (modules 2,4-10) which are the first principal components and summarize expression patterns for each module. Modules 1 and 3 are shown in Fig. 5 and not redrawn here. The vertical axes indicate expression values relative to the mean expression across all time points, which are indicated in the horizontal axes. RNA and protein fold changes compared to the 0 h time point were scaled and modified using a rank-order normalization. Numbers in parentheses denote the number of genes in each module (purple lines, RNA; blue lines, protein).

Supplementary Fig. 21

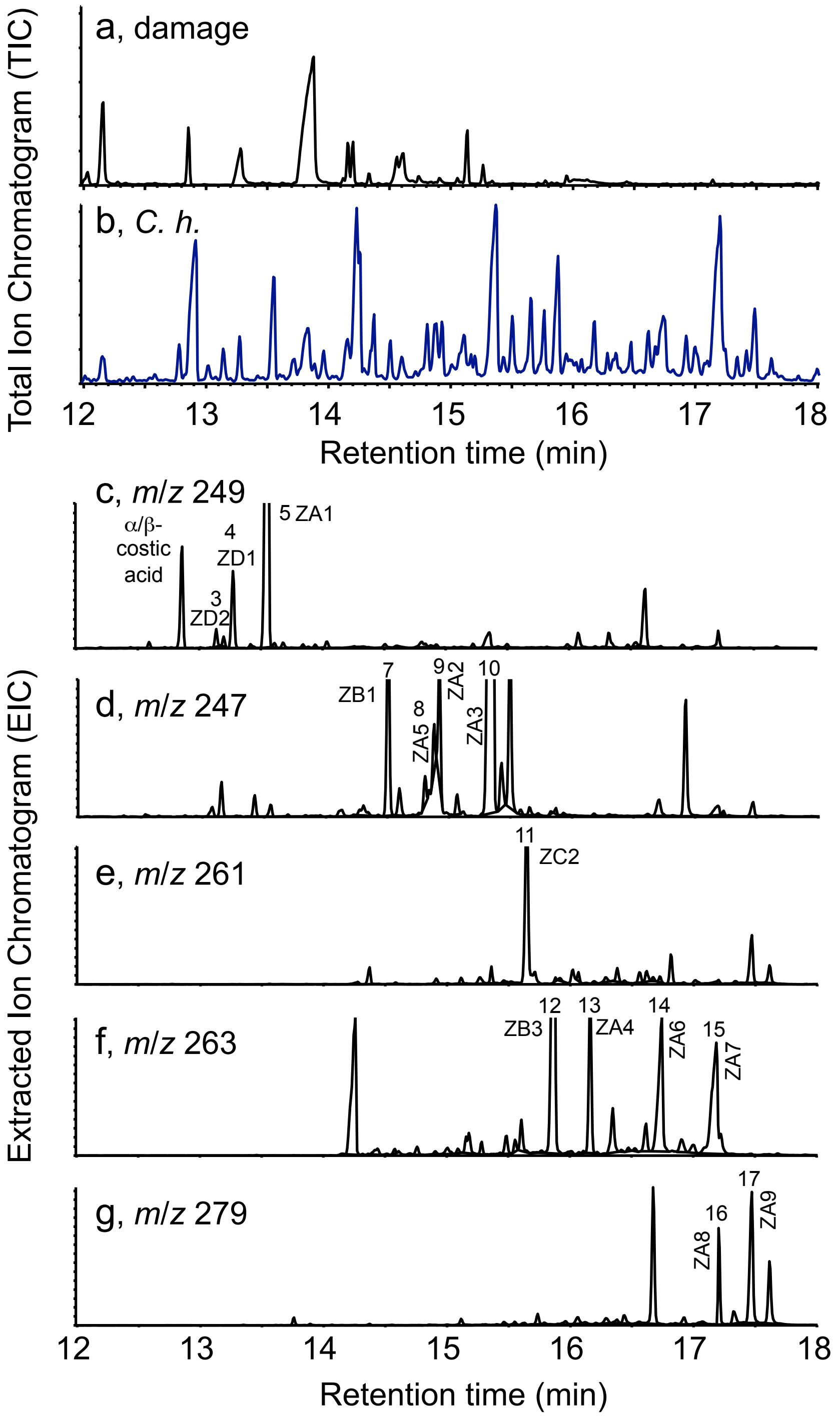


**Supplementary Fig. 21 | A complex blend of zealexins accumulate following stem infection with a necrotrophic fungal pathogen. a,** GC/(+)CI-MS total ion chromatograms (TIC) of extracts from representative maize stems 3 days after wounding or **b**, inoculation with *Cochliobolus heterostrophus* (*C. h.* 100 μl of $10^7$ spores $ml^{-1}$) analyzed as methyl ester derivatives. **c-g**, Representative EIC traces of complex oxygenated zealexins following SLB infection. With reference to compound numbering in Supplementary Fig. 1, representative retention times (RT) in order and GC/(+)CI-MS *m/z* ions are as follows (1) β-bisabolene $[M+H]^+$ *m/z* 205, RT 9.81 min (not shown); (2) β-macrocarpene $[M+H]^+$ *m/z* 205; RT, 9.85 min (not shown) **c**, α/β-costic acids *m/z* 249, RT, 12.86 min; (3) ZD2 $[M+H]^+$ *m/z* 249, RT 13.14 min; (4) ZD1 $[M+H]^+$ *m/z* 249, RT 13.27 min; (5) ZA1 $[M+H]^+$ *m/z* 249, RT 13.55 min; (6) ZC1 $[M+H]^+$ *m/z* 245, RT 14.12 (not shown) **d**, (7) ZB1 $[M+H]^+$ *m/z* 247, RT 14.51; (8) ZA5 $[M+H]^+$ *m/z* 265 (fragment $[M\text{-}H_2O]^+$ *m/z* 247 shown) RT 14.88; (9) ZA2 $[M\text{-}H_2O]^+$ *m/z* 247, RT 14.93; (10) ZA3 $[M\text{-}H_2O]^+$ *m/z* 247, RT 15.32. **e**, (11) ZC2 $[M+H]^+$ *m/z* 261, RT 15.65. **f**, (12) ZB3 $[M+H]^+$ *m/z* 263, RT 15.87; (13) ZA4 $[M+H]^+$ *m/z* 263, RT 16.17; (14) ZA6 $[M\text{-}2H_2O]^+$ *m/z* 245 (fragment $[M\text{-}H_2O]^+$ *m/z* 263 shown), RT 16.75; (15) ZA7 $[M\text{-}2H_2O]^+$ *m/z* 245 (fragment $[M\text{-}H_2O]^+$ *m/z* 263 shown), RT 17.20. **g**, (16) ZA8 $[M+H]^+$ *m/z* 279, RT 17.21; (17) ZA9 $[M+H]^+$ *m/z* 279, RT 17.48. In each section (**a-g**), the y axis denotes relative abundance of ions.

Supplementary Fig. 22

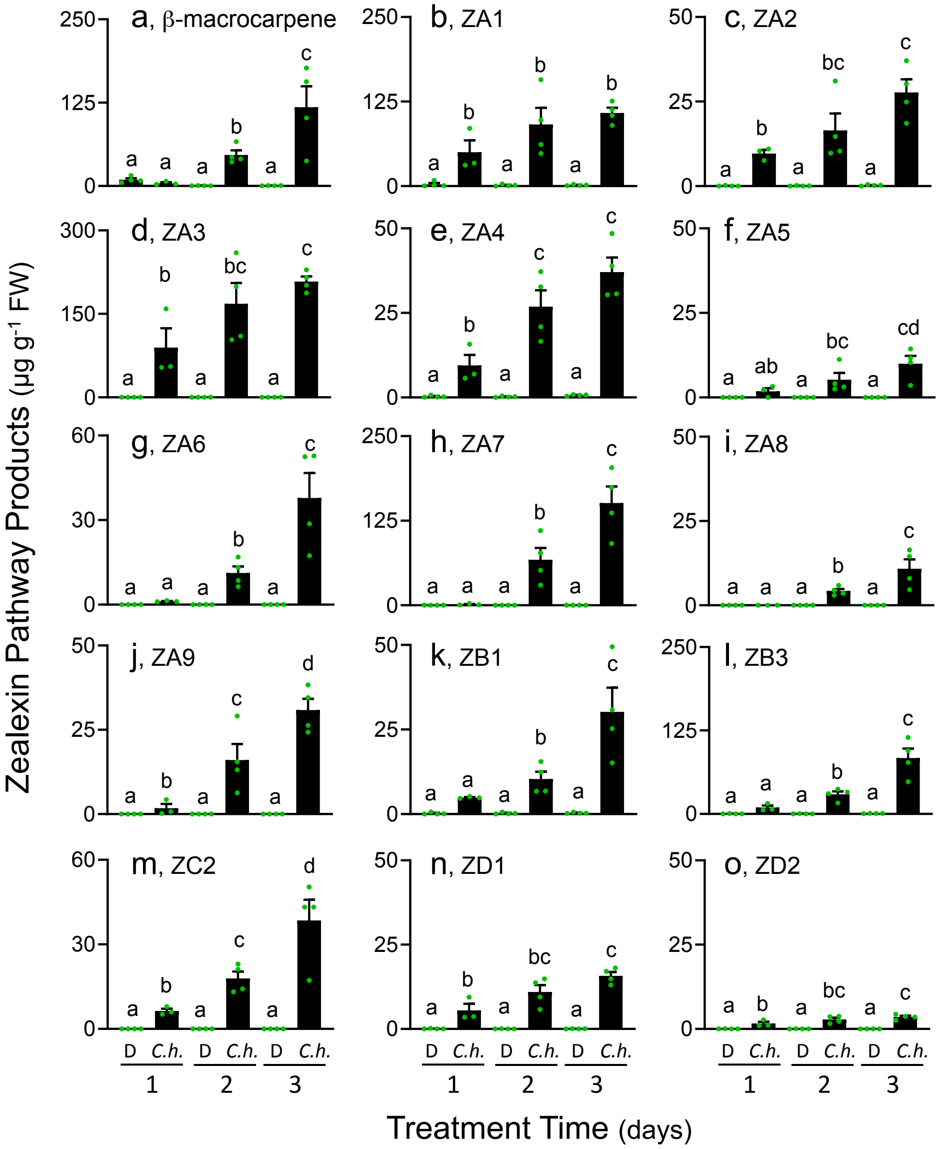


**Supplementary Fig. 22 | Quantification of zealexin accumulation following stem infection with *C. heterostrophus* supports predominant endproducts.** Time course of zealexin accumulation 1, 2 and 3 d after maize stems were either damaged and treated with 100 µl $H_2O$ (D) or inoculated with *C. heterostrophus* (*C. h.* 100 µl of $10^7$ spores $ml^{-1}$) analyzed as GC/(+)CI-MS. **a,** β-macrocarpene, **b**, ZA1; **c**, ZA2; **d**, ZA3; **e**, ZA4; **f**, ZA5; **g**, ZA6; **h**, ZA7; **i**, ZA8; **j**, ZA9; **k**, ZB1; **l**, ZB3; **m**, ZC2; **n**, ZD1; **o**, ZD2. Error bars in indicate mean ± s.e.m. (*n* = 3-4 biologically independent replicates). Within plots, different letters (a–d) represent significant differences (one-way ANOVA followed by Tukey's test corrections for multiple comparisons, *P* < 0.05).

Supplementary Fig. 23

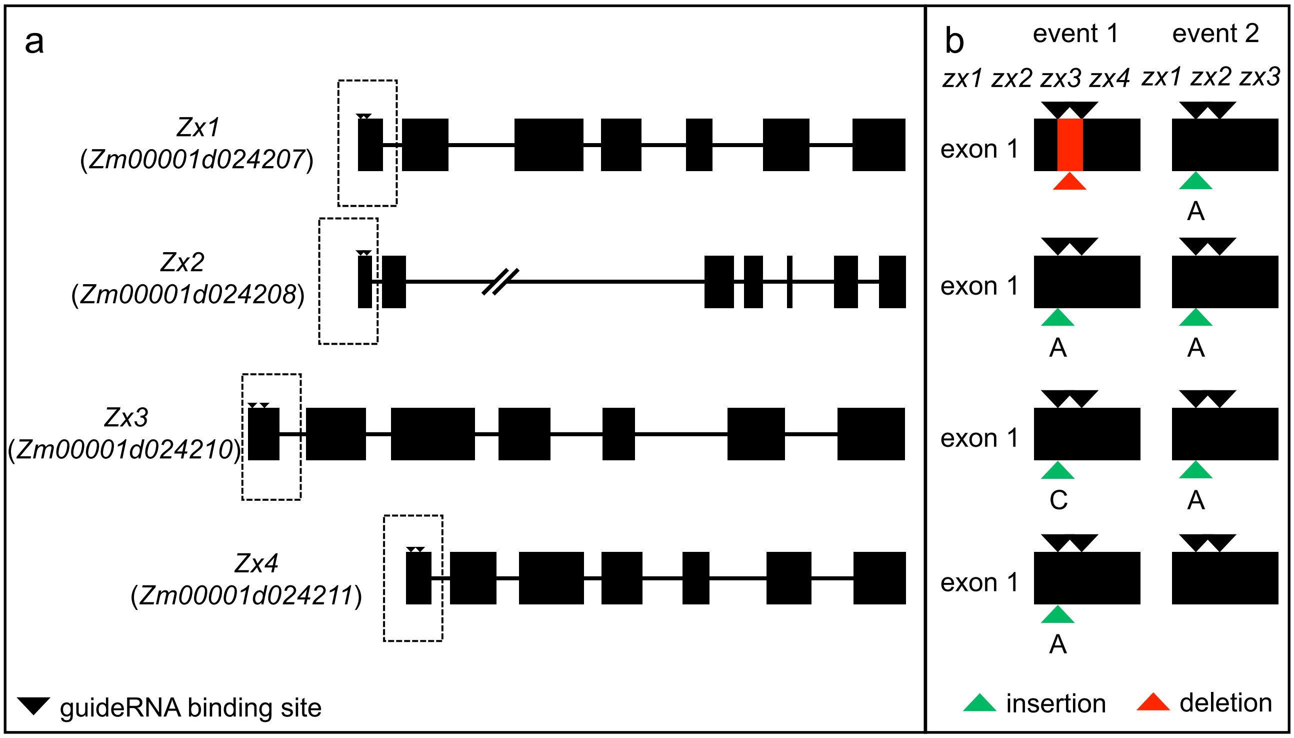


c

***Zx1***

WT ACTGCAGATGGTCCTCGCCTCGGCCAGCAGGAGATGAAGAAGATGTC

*zx1 zx2 zx3 zx4* ACTGCAGATGGTCCTCG------------------------------

*zx1 zx2 zx3* ACTGCAGATGGTCCTCGaCCTCGGCCAGCAGGAGATGAAGAAGATGTC

***Zx2***

WT ACTGCAGATGGTCCTCGCCTCGGCCAGCAGGAGATGAAGAAGATGT

*zx1 zx2 zx3 zx4* ACTGCAGATGGTCCTCGaCCTCGGCCAGCAGGAGATGAAGAAGATGT

*zx1 zx2 zx3* ACTGCAGATGGTCCTCGaCCTCGGCCAGCAGGAGATGAAGAAGATGT

***Zx3***

WT ACTACAGATGGTCCTCGCCTCGGCCAGCAGGAGATGAAGAAGATGT

*zx1 zx2 zx3 zx4* ACTACAGATGGTCCTCGcCCTCGGCCAGCAGGAGATGAAGAAGATGT

*zx1 zx2 zx3* ACTACAGATGGTCCTCGaCCTCGGCCAGCAGGAGATGAAGAAGATGT

***Zx4***

WT ACTACAGATGGTCCTCGCCTCGGCGAGCAGGAGATGAAGAAGATGT

*zx1 zx2 zx3 zx4* ACTACAGATGGTCCTCGaCCTCGGCGAGCAGGAGATGAAGAAGATGT

*zx1 zx2 zx3* ACTACAGATGGTCCTCGCCTCGGCGAGCAGGAGATGAAGAAGATGT

**Supplementary Fig. 23 | Graphical representation of the zealexin biosynthetic pathway genes *Zx1 to Zx4* and corresponding triple (*zx1 zx2 zx3*) and quadruple (*zx1 zx2 zx3 zx4*) mutants using CRISPR–Cas9. a**, Gene structure of maize β-macrocarpene synthase genes *Zx1*, *Zx2*, *Zx3* and *Zx4*. Small arrows on top of exon one denote gRNA 1 and gRNA 2-binding sites, respectively. **b**, Graphical representation of the first exon of *Zx1 to Zx4* and their respective mutations in triple and quadruple mutants created by CRISPR–Cas9. **c**, Partial sequence of *Zx1*, *Zx2*, *Zx3* and *Zx4* showing gRNA-binding site (underlined) and the respective mutations (--- deletion, green insertion of adenine or cytosine).

Supplementary Fig. 24

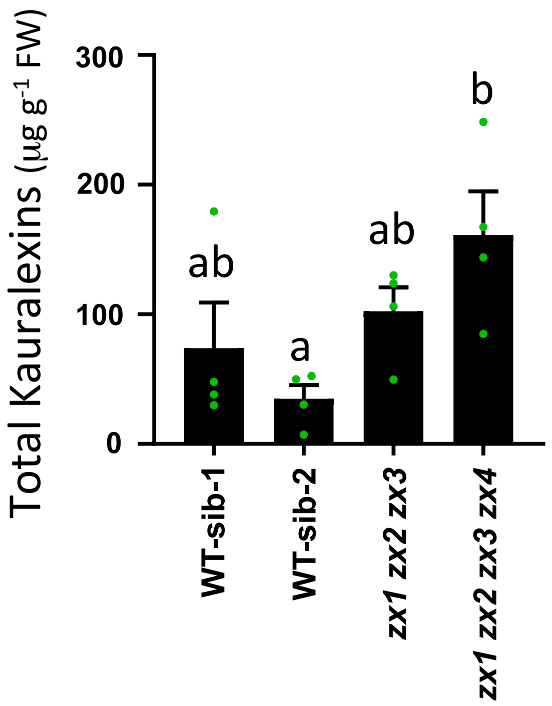


**Supplementary Fig. 24 | Levels of total kauralexins present in zealexin triple (*zx1 zx2 zx3*) and quadruple (*zx1 zx2 zx3 zx4*) mutant plants in response to *Fusarium graminearum*.** CRISPR/Cas9 derived triple and quadruple mutants and the respective wildtype siblings (WT-sib) were grown for 25 days and stem-inoculated with *Fusarium graminearum* (10 μL of 1.5 x $10^5$ conidia $ml^{-1}$) for 10 days. Kauralexins were measured by GC-MS. Error bars in the bar chart indicate mean ± s.e.m ($n$ = 4 biologically independent replicates). Within plots, different letters (a–b) represent significant differences (one-way ANOVA followed by Tukey's test corrections for multiple comparisons, $P < 0.05$).

Supplementary Fig. 25

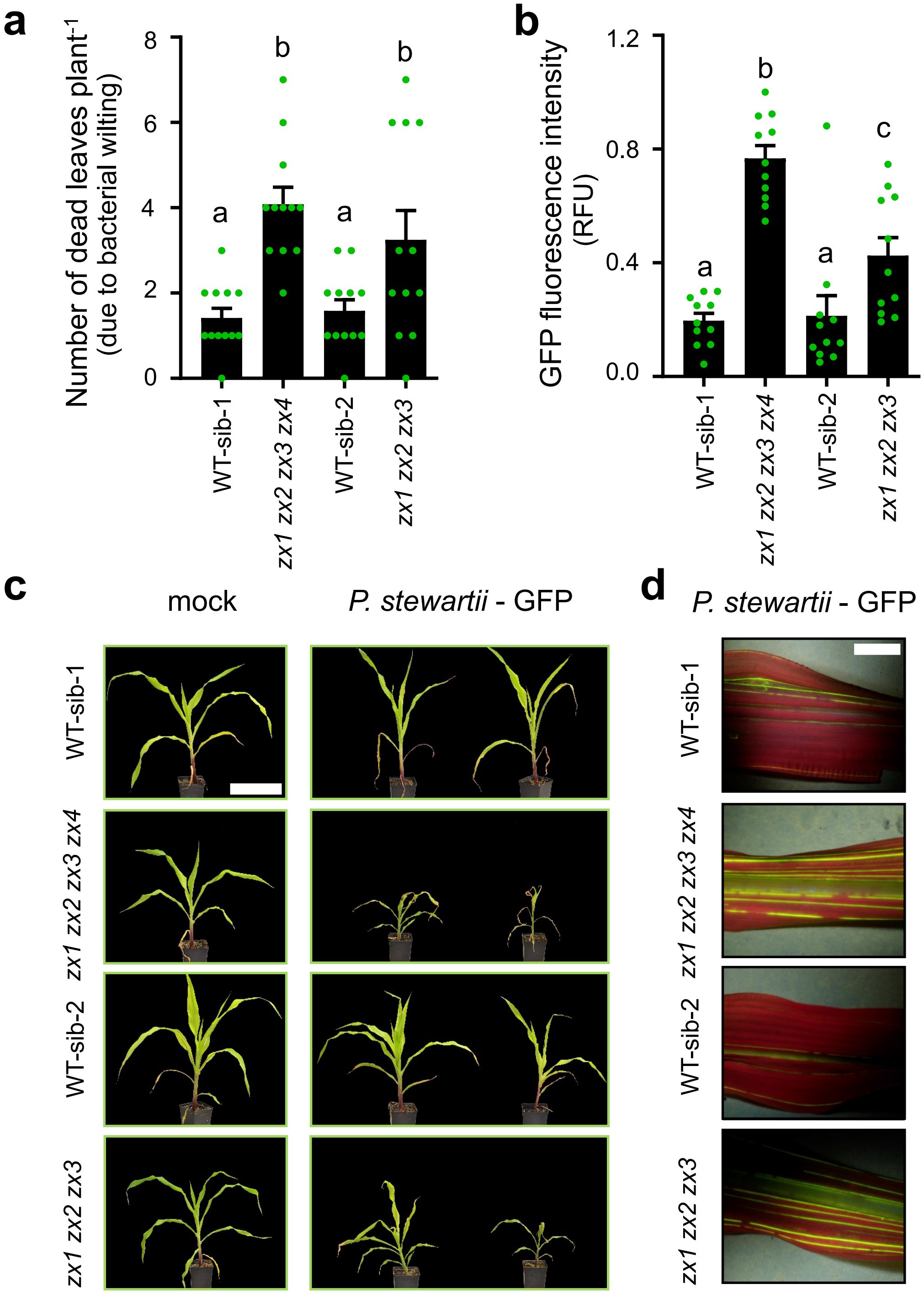


**Supplementary Fig. 25 | Zealexin deficient triple and quadruple mutant plants display increased susceptibility to Stewarts wilt (*Pantoea stewartii*), a xylem-dwelling bacterium**. **a**, Quantification of wilting symptoms in *P. stewartii* infected plants of triple (*zx1 zx2 zx3*), quadruple (*zx1 zx2 zx3 zx4*) and the respective wild type siblings (WT-sib). **b**, Quantification of relative GFP fluorescence in *P. stewartii* infected plants. GFP fluorescence intensity was normalized to the highest fluorescence value and is shown as relative fluorescence units (RFU). Maize seedlings were infected with *P. stewartii*-GFP (DC283-GFP) or mock inoculated (controls, shown in **c**); GFP fluorescence was examined after 5 days post inoculation (dpi) while wilting was examined at 16 dpi. In **a** and **b**, error bars indicate mean ± s.e.m. (*n* = 11-12 biologically independent replicates). Within plots, different letters (a–c) represent significant differences (one-way ANOVA followed by Tukey's test corrections for multiple comparisons; $P < 0.05$). **c**, Representative mock-inoculated or *P. stewartii* infected WT-sib-1, *zx1 zx2 zx3 zx4*, WT-sib-2, and *zx1 zx2 zx3* plants. Bar = 20 cm. **d**, Representative blue light illuminated leaf sections of *P. stewartii* infected WT-sib-1, *zx1 zx2 zx3 zx4*, WT-sib-2, and *zx1 zx2 zx3* plants. Bar = 1 cm.

Supplementary Fig. 26

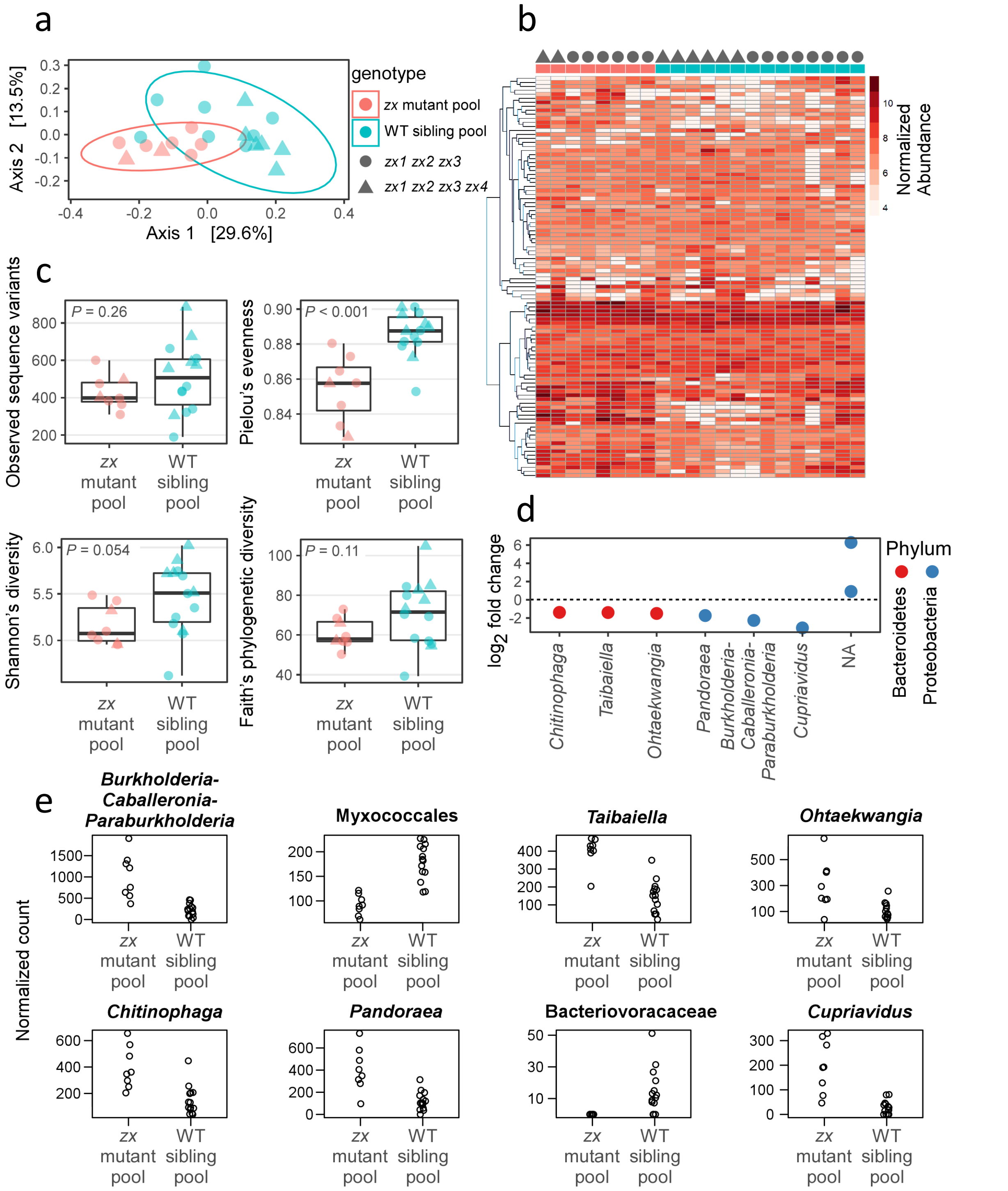


**Supplementary Fig. 26 | Reduced zealexin production substantially alters the bacterial microbiome associated with corn roots, lowering overall evenness and altering the abundances of particular taxa. a**, Principal coordinates plot of bacterial communities associated with corn roots. Parenthetical percentages indicate the percentage of variation explained by each axis. **b**, heatmap showing normalized counts in each sample for the 100 most abundant bacterial sequence variants associated with corn roots. Rows correspond to sequence variants, and were hierarchically clustered. c, diversity indices for bacterial communities associated with corn roots. *P* values represent Student's *t* test, two-tailed distribution. **d**, Log2 fold-changes for sequence variants, identified to genus and phylum, that differed significantly in abundance between wild type vs. zealexin knockout plants (DESeq with Wald test, $P < 0.05$). **e**, normalized counts across samples, for those sequence variants shown in **d**.

**Supplementary Table 6: β-macrocarpene synthase derived zealexins purified and elucidated from fungal elicited maize tissue**. All NMR data was acquired in 2.5-mm NMR tubes (Norell) at 22 °C using a 5-mm TXI cryoprobe (Bruker Corporation) and a Bruker Avance II 600 console (600 MHz for 1H and 151 MHz for 13C). One- and two-dimensional $^{1}$H and $^{13}$C NMR spectroscopy with heteronuclear multiple-bond correlations (HMBC) and assigned carbon numbers (C) were used for structural elucidation. In all figures, correlated spectroscopy (COSY) correlations are shown in red bonds. Zealexin NMR data was collected in $CDCl_3$ (ZA5, ZB3 methyl ester), acetonitrile-($d_3$) (ZA8, ZA9) and benzene ($d_6$) (ZD1, ZD2, ZA6 methyl ester, ZA7 methyl ester) and the spectra were referenced to the following chemical shifts $^{1}$H 7.26 ppm and $^{13}$C 77.4 ppm for $CDCl_3$; $^{1}$H 7.16 ppm and $^{13}$C 128.1 ppm for benzene-$d_6$; and $^{1}$H 1.94 ppm and $^{13}$C 1.4 ppm for acetonitrile-$d_3$. NMR spectra were processed using Bruker Topspin 2.0 and MestReNova (Mestrelab Research) software packages. For each drawn compound structure, all carbons (C) are numbered. Coupling constants are given in Hertz [Hz]. In plant tissues all detected zealexins occur as free carboxylic acids. In select cases, for example ZB3, ZA6 and ZA7, final purification prior to NMR utilized methyl ester derivatives.

**Supplementary Table:** **6a.** Summarized NMR spectral data for ZD1
**6b.** Summarized NMR spectral data for ZD2
**6c.** Summarized NMR spectral data for ZA5
**6d.** Summarized NMR spectral data for ZB3 (methyl ester)
**6e.** Summarized NMR spectral data for ZA6 (methyl ester)
**6f.** Summarized NMR spectral data for ZA7 (methyl ester)
**6g.** Summarized NMR spectral data for ZA8
**6h.** Summarized NMR spectral data for ZA9

**Supplementary Table: 6a.** Summarized NMR spectral data for ZD1

4-(6-methylhepta-1,5-dien-2-yl)cyclohex-1-ene-1-carboxylic acid

| Position | δ $^{13}$C [ppm] | δ $^{1}$H [ppm] | J coupling constants [Hz] | HMBC correlations |
|---|---|---|---|---|
| 1 | 72.2 | - | | |
| 2 | 31.5 | 2H 1.82, 1.71 | 1.82, 1H, m<br>1.71, 1H, m | C4, C5 |
| 3 | 21.9 | 2H 2.32 | m | |
| 4 | 130.1 | - | | |
| 5 | 137.9 | 1H 6.88 | t J=1.54 | C1, C3, C4, C15 |
| 6 | 37.1 | 2H 2.55, 2.21 | 2.55, 1H, q J=3.1<br>2.21 1H m | C7, C12 |
| 7 | 168.3 | - | | |
| 8 | 51.7 | 2H 2.18 | d *J* = 2.0 | C9, C10, C11, C13, C14 |
| 9 | 34.4 | - | | |
| 10 | 200.9 | - | | |
| 11 | 40 | 2H 2.30 | 2.30, 2H, dd *J* =4.9, 1.5 | |
| 12 | 123 | 1H 6.01 | t J=1.6 | C1, C7, C8, C11 |
| 13 | 27.9 | 3H 1.01 | s | C8, C9, C10, C11, C14 |
| 14 | 28.3 | 3H 1.02 | s | C8, C9, C10, C11, C13 |
| 15 | 168 | - | | |

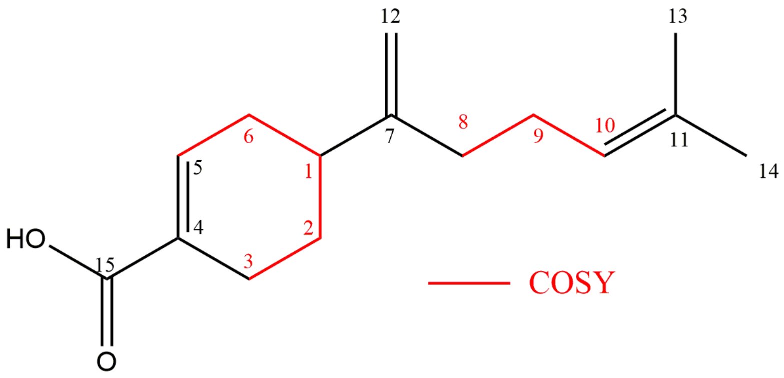

**Supplementary Table: 6b.** Summarized NMR spectral data for ZD2

(E)-2-methyl-6-(4-methylcyclohex-3-en-1-yl)hepta-2,6-dienoic acid

| Position | δ $^{13}C$ [ppm] | δ $^{1}H$ [ppm] | J coupling constants [Hz] | HMBC correlations |
|---|---|---|---|---|
| 1 | 39.6 | 1H 1.98 (m) | m | C3, C8 |
| 2 | 28.2 | 2H 1.67, 1.38 | 1.67, 1H, m<br>1.38, 1H, m | C1, C4 |
| 3 | 30.9 | 2H 1.91, 1.85 | 1.91, 1H<br>1.85, 1H | C1, C2, C4, C5 |
| 4 | 133.4 | - | | |
| 5 | 120.8 | 1H 5.42 | 5.42, 1H, m | C1, C3, C6, C15 |
| 6 | 31.2 | 2H 2.04, 1.98 | 2.04, 1H, m<br>1.98, 1H, m | C1 |
| 7 | 152.9 | - | | |
| 8 | 33.1 | 2H 1.89 | m | C1, C7, C9, C10, C12 |
| 9 | 27.3 | 2H 2.02 | m | C7, C8, C10, C11 |
| 10 | 144.7 | 1H 7.01 | s | C8, C9, C11, C14, C13 |
| 11 | 127.6 | - | | |
| 12 (14) | 107.8 | 2H 4.83, 4.71 | 4.83, 1H, s<br>4.71, 1H, s | C1, C7, C8 |
| 13 | 173.8 | - | | |
| 14 (12) | 11.8 | 3H, 1.76 | s | C8, C10, C11, C13 |
| 15 | 23.5 | 3H, 1.64 | s | C3, C4, C5 |

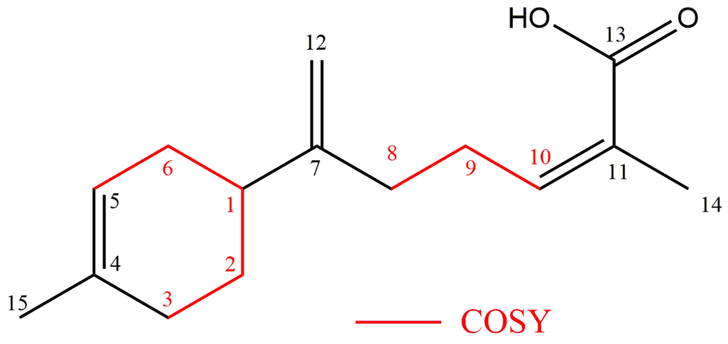

**Supplementary Table: 6c.** Summarized NMR spectral data for ZA5

2-hydroxy-5',5'-dimethyl-[1,1'-bi(cyclohexane)]-1',3-diene-4-carboxylic acid

| Position | δ $^{13}C$ [ppm] | δ $^{1}H$ [ppm] | J coupling constants [Hz] | HMBC correlations |
|---|---|---|---|---|
| 1 | 50.2 | 1H 2.04 | m | |
| 2 | 25.7 | 2H 1.77, 1.60 | 1.77, 1H, m<br>1.60, 1H m | C1, C3, C4, C6 |
| 3 | 24.6 | 2H 2.41, 2.27 | 2.41, 1H, dd J=18.2, 5.3<br>2.27, 1H, m | C1, C4, C5 |
| 4 | 136.6 | - | | |
| 5 | 142.8 | 1H 7.02 | s | C1, C3, C4, C7, C15 |
| 6 | 68.5 | 1H 4.25 | d J=9.3 | C4, C5, C7 |
| 7 | 136.3 | - | | |
| 8 | 39 | 2H 1.71 | d J=8.4 | C7, C14 |
| 9 | 29.1 | - | | |
| 10 | 35.2 | 2H 1.35 | t J=6.5 | C8, C11, C12, C13, C14 |
| 11 | 23.3 | 2H 2.07 | m | C7, C12 |
| 12 | 123.6 | 1H 5.57 | s | C1, C8, C10, C11 |
| 13 | 27.9 | 3H 0.91 | s | C7(w), C8, C9, C10, C14 |
| 14 | 28.7 | 3H 0.93 | s | C7(w), C8, C9, C10, C13 |
| 15 | 171.8 | - | | |

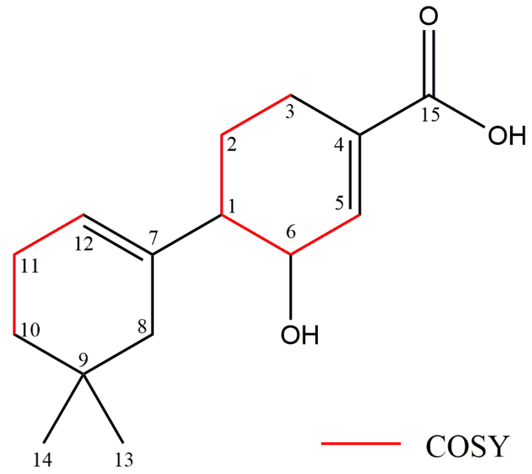

**Supplementary Table: 6d.** Summarized NMR spectral data for ZB3 (methyl ester)

6'-hydroxy-5',5'-dimethyl-[1,1'-bi(cyclohexane)]-1,1',3-triene-4-carboxylic acid methyl ester

| Position | δ $^{13}$C [ppm] | δ $^{1}$H [ppm] | J coupling constants [Hz] | HMBC correlations |
|---|---|---|---|---|
| 1 | 142.6 | - | | |
| 2 | 24.3 | 2H 2.52, 2.44 | 2.52, 1H,<br>2.44, 1H, m | C1, C3, C4, C6 |
| 3 | 22 | 2H 2.52 | m | C1, C2, C4, C5 |
| 4 | 126.3 | - | | |
| 5 | 134.6 | 1H 7.11 | d, J=6.13 | C1, C3, C4, C6, C15 |
| 6 | 118.4 | 1H 6.37 | d, J=5.85 | C3, C4, C7 |
| 7 | 137.6 | - | | |
| 8 | 71.9 | 1H 3.92 | br s | C1, C7, C9, C10, C11, C12 |
| 9 | 33.8 | - | | |
| 10 | 28.8 | 2H 1.26, 1.68 | 1.26, 1H,<br>1.68, 1H, m | C1, C8, C9, C11, C12 |
| 11 | 24.2 | 2H 2.46, 2.62 | 2.46, 1H,<br>2.62, 1H, m | C8, C9, C10, C12 |
| 12 | 129.3 | 1H 6.12 | t, J=4.17 | C1, C10, C11, C8 |
| 13 | 23.7 | 3H 0.84 | s | |
| 14 | 26.4 | 3H 1.07 | s | |
| 15 | 167.9 | - | | |
| 16 | 51.7 | 3H 3.76 | s | C15 |

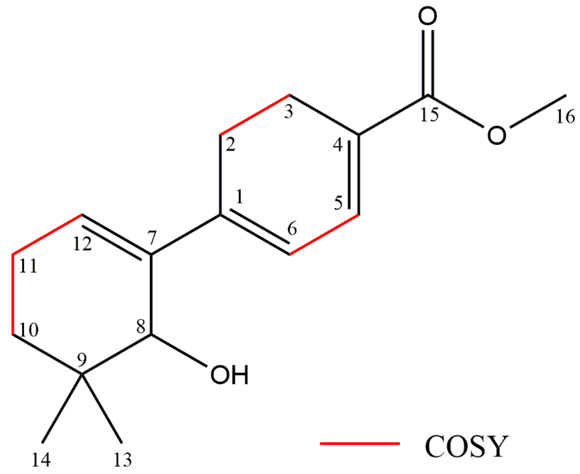

**Supplementary Table: 6e.** Summarized NMR spectral data for ZA6 (methyl ester)

1,4'-dihydroxy-5',5'-dimethyl-[1,1'-bi(cyclohexane)]-1',3-diene-4-carboxylic acid methyl ester

| Position | δ $^{13}C$ [ppm] | δ $^{1}H$ [ppm] | J coupling constants [Hz] | HMBC correlations |
|---|---|---|---|---|
| 1 | 71.2 | - | | |
| 2 | 31.4 | 2H 1.42 | m | C1, C3, C4, C7, C6 |
| 3 | 22.4 | 2H 2.49 | m | C1, C2, C4, C5 |
| 4 | 130 | - | | |
| 5 | 136.1 | 1H 6.92 | m | C1, C15 |
| 6 | 37.2 | 2H 2.07, 1.92 | 2.07, 1H, m<br>1.92, 1H, m | C2, C4, C5, C7 |
| 7 | 141.3 | - | | |
| 8 | 36.6 | 2H 1.81, 1.64 | 1.81, 1H, m<br>1.64, 1H, m | C7, C9, C10, C12, C13, C14 |
| 9 | 34 | - | | |
| 10 | 73.1 | 1H 3.24 | 3.24, 1H, m | C8, C9, C13, C14 |
| 11 | 32.1 | 2H 2.12, 1.82 | 2.12, 1H, m<br>1.82, 1H, m | C7, C9, C10, C12 |
| 12 | 116.7 | 1H 5.33 | 5.33, 1H, m | C8, C10, C11 |
| 13 | 26.3 | 3H 0.84 | s | C1, C7, C8, C9, C10, C14 |
| 14 | 21.7 | 3H 0.86 | s | C9, C10, C13 |
| 15 | 170.8 | - | | |
| 16 | 50.9 | 3H 3.45 | s | |

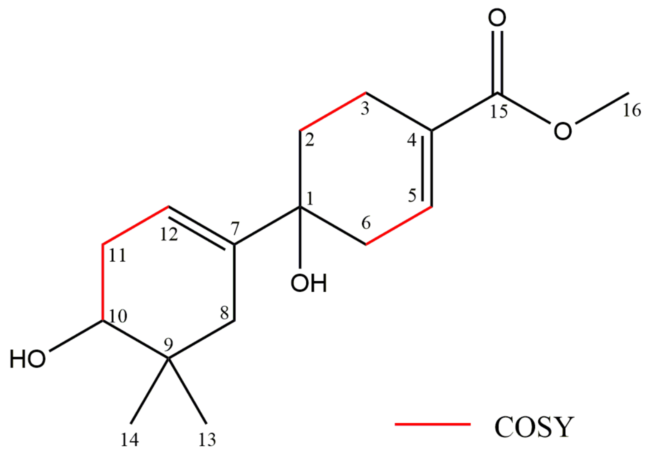

**Supplementary Table: 6f.** Summarized NMR spectral data for ZA7 (methyl ester)

4',6'-dihydroxy-5',5'-dimethyl-[1,1'-bi(cyclohexane)]-1',3-diene-4-carboxylic acid methyl ester

| Position | δ $^{13}C$ [ppm] | δ $^{1}H$ [ppm] | J coupling constants [Hz] | HMBC correlations |
|---|---|---|---|---|
| 1 | 36.7 | 1H 2.13 | 2.13, 1H, m | C2, C6 |
| 2 | 27.9 | 2H 1.65, 1.21 | 1.65, 1H, m<br>1.21, 1H, m | C1, C3, C4, C6, C7 |
| 3 | 24.9 | 2H 2.51, 2.31 | 2.51, 1H, m<br>2.31, 1H, m | C1, C2, C4, C5 |
| 4 | 130.3 | - | | |
| 5 | 139.3 | 1H 7.06 | 7.06, 1H, m | C1, C2, C6, C15 |
| 6 | 31.3 | 2H 2.04, 1.90 | 2.04, 1H, m<br>1.90, 1H, m | C1, C3, C4, C5 |
| 7 | 142 | - | | |
| 8 | 75.8 | 1H 3.38 | 3.38, 1H, s | C1, C7, C9, C10, C11, C12, C13, C14 |
| 9 | 39.1 | - | | |
| 10 | 69.8 | 1H 3.63 | 3.63, 1H, dd J=8.8, 5.9 | C8, C9, C11, C12, C13, C14 |
| 11 | 32.4 | 2H 2.12, 1.75 | 2.12, 1H, m 1.75, 1H, m | C10, C12, C17 |
| 12 | 120.4 | 1H 5.04 | 5.04, 1H,<br>ddd J=4.7, 2.9, 1.0 | C1, C7, C8, C10, C11 |
| 13 | 21.9 | 3H 1.04 | s | C8, C9, C10, C14 |
| 14 | 17.8 | 3H 0.75 | s | C8, C9, C10, C13 |
| 15 | 167.3 | - | | |
| 16 | 50.88 | 3H 3.47 | s | |

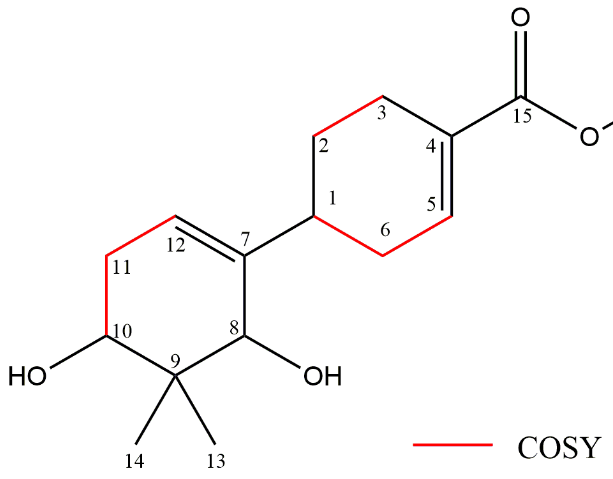

**Supplementary Table: 6g.** Summarized NMR spectral data for ZA8

1-hydroxy-5',5'-dimethyl-4'-oxo-[1,1'-bi(cyclohexane)]-1',3-diene-4-carboxylic acid

| Position | δ $^{13}$C [ppm] | δ $^{1}$H [ppm] | J coupling constants [Hz] | HMBC correlations |
|---|---|---|---|---|
| 1 | 72.2 | - | | |
| 2 | 31.5 | 2H 1.82, 1.71 | 1.82, 1H, m<br>1.71, 1H, m | C4, C5 |
| 3 | 21.9 | 2H 2.32 | m | |
| 4 | 130.1 | - | | |
| 5 | 137.9 | 1H 6.88 | t J=1.54 | C1, C3, C4, C15 |
| 6 | 37.1 | 2H 2.55, 2.21 | 2.55, 1H, q J=3.1<br>2.21 1H m | C7, C12 |
| 7 | 168.3 | - | | |
| 8 | 51.7 | 2H 2.18 | d *J* = 2.0 | C9, C10, C11, C13, C14 |
| 9 | 34.4 | - | | |
| 10 | 200.9 | - | | |
| 11 | 40 | 2H 2.30 | 2.30, 2H, dd *J* =4.9, 1.5 | |
| 12 | 123 | 1H 6.01 | t J=1.6 | C1, C7, C8, C11 |
| 13 | 27.9 | 3H 1.01 | s | C8, C9, C10, C11, C14 |
| 14 | 28.3 | 3H 1.02 | s | C8, C9, C10, C11, C13 |
| 15 | 168 | - | | |

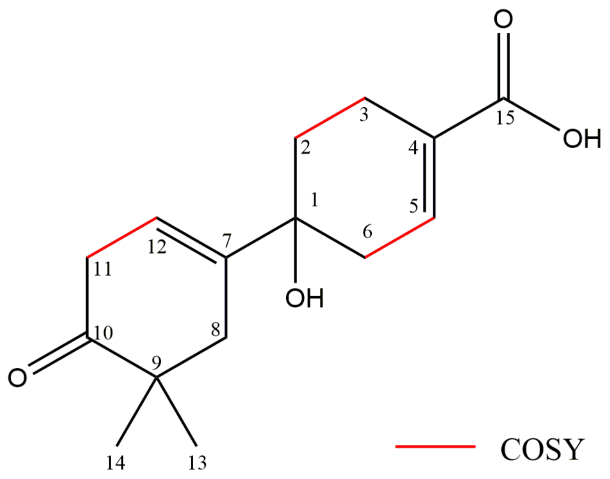

**Supplementary Table: 6h.** Summarized NMR spectral data for ZA9

6'-hydroxy-5',5'-dimethyl-4'-oxo-[1,1'-bi(cyclohexane)]-1',3-diene-4-carboxylic acid

| Position | δ $^{13}C$ [ppm] | δ $^{1}H$ [ppm] | J coupling constants [Hz] | HMBC correlations |
|---|---|---|---|---|
| 1 | 37.4 | 1H 2.63 | br m | C7 |
| 2 | 27.3 | 2H 1.97, 1.55 | 1.97, 1H, m<br>1.55, 1H, m | C1, CC4, C6 |
| 3 | 24.5 | 2.32 1H m, 2.26 1H m | 2.32, 1H, m<br>2.26, 1H, m | C6 |
| 4 | 130.4 | - | | |
| 5 | 139.8 | 1H 6.99 | br m | C1, C3, C15 |
| 6 | 30.8 | 2H 2.42, 2.25 | 2.42, 1H, m<br>2.25, 1H, m | C2, C4, C5 |
| 7 | 167.4 | - | | |
| 8 | 74.6 | 1H 4.00 | d J=6.2 | C7, C11, C12, C13, C14 |
| 9 | 38.4 | - | | |
| 10 | 199.6 | - | | |
| 11 | 48.3 | 2h 2.39, 2.10 | 2.39, 1H, m<br>2.10, 1H, m | C8, C9, C10, C13, C14 |
| 12 | 124.1 | 1H 5.69 | br s | C1, C8, C11 |
| 13 | 26.4 | 3H 0.97 | s | C8, C9, C10, C11, C14 |
| 14 | 23.3 | 0.98 3H s | s | C8, C9, C10, C11, C13 |
| 15 | 168.1 | - | | |

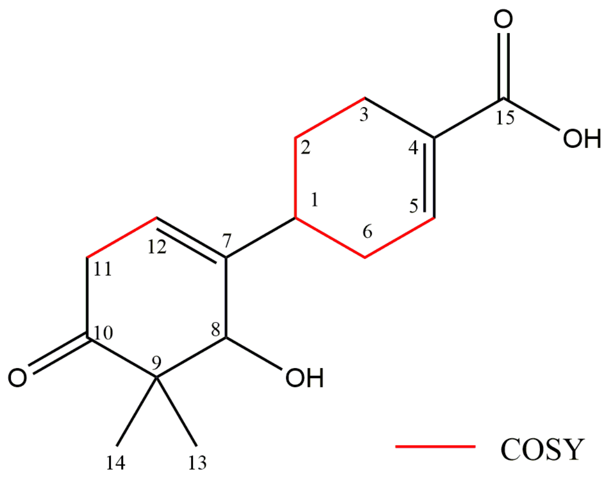
